# Chemical interplay and complementary adaptative strategies toggle bacterial antagonism and co-existence

**DOI:** 10.1101/2021.01.11.426172

**Authors:** Carlos Molina-Santiago, David Vela-Corcía, Daniel Petras, Luis Díaz-Martínez, Alicia Isabel Pérez-Lorente, Sara Sopeña-Torres, John Pearson, Andrés Mauricio Caraballo-Rodríguez, Pieter C. Dorrestein, Antonio de Vicente, Diego Romero

**Affiliations:** Instituto de Hortofruticultura Subtropical y Mediterránea “La Mayora”, Universidad de Málaga-Consejo Superior de Investigaciones Científicas (IHSM-UMA-CSIC), Departamento de Microbiología, Universidad de Málaga, Bulevar Louis Pasteur 31 (Campus Universitario de teatinos), 29071, Málaga, Spain; University of California San Diego, Scripps Institution of Oceanography, La Jolla, USA; University of California San Diego, Collaborative Mass Spectrometry Innovation Center, La Jolla, USA; Nano-imaging Unit, Andalusian Centre for Nanomedicine and Biotechnology, BIONAND, Málaga, Spain

**Author notes:** Corresponding author: Diego Romero.

## Abstract

Bacterial communities are in a continuous adaptive and evolutionary race for survival. A myriad of molecules that kill, defend, or mediate communication between bacterial cells of different lineages shape the final structure of the microbial community. In this work we expand our knowledge on the chemical interplay and specific mutations that modulate the transition from antagonism to co-existence between two plant-beneficial bacteria, *Pseudomonas chlororaphis* PCL1606 and *Bacillus amyloliquefaciens* FZB42. We reveal that the bacteriostatic activity of bacillaene produced by *Bacillus* relies on an interaction with the protein elongation factor FusA and how mutations in this protein lead to tolerance to bacillaene and other protein translation inhibitors. Additionally, we describe how the unspecific tolerance to antimicrobials associated with mutations in the glycerol kinase GlpK is provoked mainly by a decrease of *Bacillus* cell membrane permeability among other pleiotropic cellular responses. We conclude that nutrient specialization and mutations in basic biological functions are bacterial evolutive and adaptive strategies that lead to the coexistence of two primary competitive bacterial species rather than their mutual eradication.

## Introduction

Microbes living in multispecies communities are continuously interacting and competing for scarce resources such as nutrients and space, which in the end are key determinants of the evolution and the success of a community^1^. Thus, an understanding of how bacterial species interact, communicate, and evolve to coexist or to defeat competitors is a major interest in microbial ecology^2^. Specifically, the plant environment often represents a competitive ecological niche where microbes are in a continuous fight for nutrients either secreted by the plant or found in the soil^3–5^. The activation of specific metabolic pathways and the production and secretion of signaling compounds, siderophores, antibiotics, and quorum-sensing molecules are bacterial factors that mediate inter- and intraspecies interactions as well as inter-kingdom communication^6–9^.

*Bacillus* and *Pseudomonas* are two of the most predominant bacterial genera found in plant microbiomes^10,11^, and their beneficial effects on plants in the fight against fungal and bacterial pathogens via direct antagonism or indirectly via the activation of plant defense mechanisms (ISR) have been deeply studied^12,13^. However, how *Pseudomonas* and *Bacillus* strains coexist has only been partially described. Separate studies have revealed: i) the relevance of sporulation and biofilm formation in the protection of *Bacillus* against *Pseudomonas* competition; and ii) molecules of *Pseudomonas* that inhibit cell differentiation of *Bacillus* and the relevance of the type VI secretion system (T6SS) as an offensive tool in close cell-to cell contact^14,15^. Despite the major role that these processes play in bacterial interactions, it is well known that bacteria produce a vast arsenal of toxins and secondary metabolites that play critical roles in the modulation of antagonistic interactions^16–18^.

*Bacillus amyloliquefaciens* FZB42 (Bamy) is a Gram-positive soil bacterium with outstanding potential for the production of non-ribosomal secondary metabolites^19^, e.g., fengycins, surfactins, bacillaene, bacillomycin D, bacillibactin, difficidin, bacilysin, and other unknown compounds, that participate in diverse biological and communicative processes^20^. In fact, it has been proposed that 8.5% of the Bamy genome is devoted to non-ribosomal biosynthesis of secondary metabolites, highlighting the relevance of these molecules to the lifestyle of this bacterium^19^. Among pseudomonads, *Pseudomonas chlororaphis* PCL1606 (Pcl) is a well-known bacterium that was isolated from the rhizosphere of avocado trees^21^ and has shown antifungal and antimicrobial activities thanks to its production of several small molecules, including pyrrolnitrins, hydrogen cyanide, and 2-hexyl-propyl-resorcinol (HPR)^22^. In addition, genomic sequence analysis has determined that Pcl produces siderophores such as pyochelin, pyoverdine, and achromobactin^22,23^.

In this work we have studied the development of bacterial interspecies interactions by investigating how the chemical interplay between Pcl and Bamy is modulated over time. We have found that, in the short-term, Pcl inhibits the growth of Bamy via the combined effects of molecules regulated by the two-component system GacA-GacS while the production of the bacteriostatic compound bacillaene protects Bamy from Pcl advance. Mutations in the specific target of the antimicrobial compounds or enzymes produced by Bamy and Pcl, with the consequent pleiotropic effect, arise as the most favorable strategies leading to the co-existence of both bacterial species. Our results serve to decipher the intricacies of bacterial communication and the evolution of antagonistic interactions that support stable mixed communities.

## Results

### The interaction between Pcl and Bamy progresses from antagonism to coexistence

Pairwise interaction experiments are routinely used as the starting point for exploring microbial antagonism or alternative ways of bacterial communication. In a previous study we found that different growth media inflicted variations in the resulting interaction between *Bacillus subtilis* and *Pseudomonas chlororaphis*^15^. The two strains come into close contact in lysogeny broth (LB) medium, while in King’s B medium, which promotes the production of secondary metabolites by *Pseudomonas* strains^24^, the two colonies were physically separated by a typical inhibition halo (Suppl. Figure 1a). This result was reproducible in the interaction between Pcl with Bamy, a closely related species to *B. subtilis* (Figure 1a top). Pcl and Bamy are strong producers of a myriad of secondary metabolites, and they are natural inhabitants of the rhizosphere where they contribute to the health and yield of plants in different ways. For these reasons, we selected these strains to characterize the mechanism of the interspecies chemical interplay.

**Figure 1.**
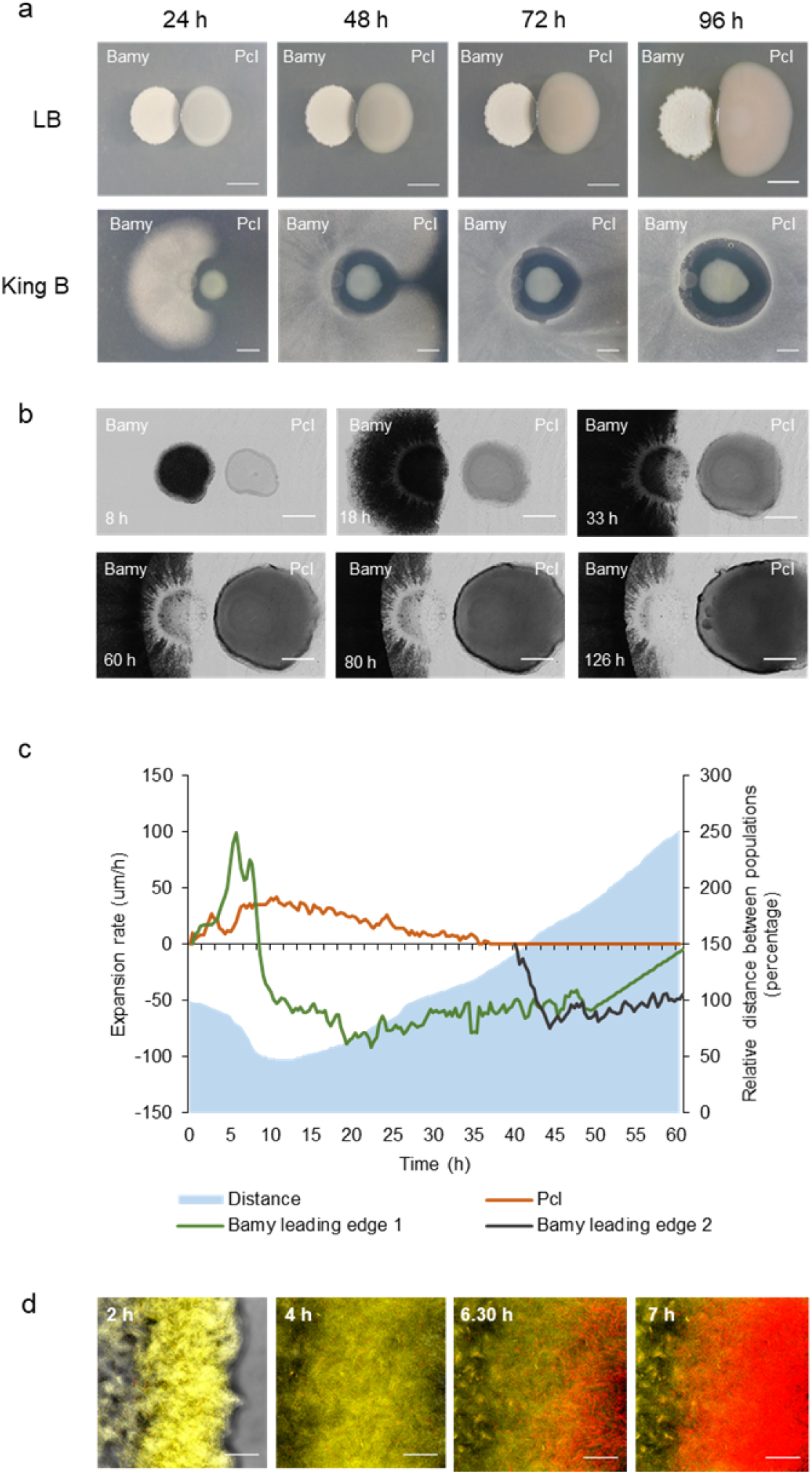
Pcl and Bamy mutually exclude in King B medium. a) Pairwise interaction time lapses between Bamy (left) and Pcl (right) on LB (top) and King B (bottom) media at 24 h, 48 h, 72 h, and 96 h. Scale = 5 mm. b) Time-lapse microscopy of the pairwise interaction between Pcl and Bamy over 126 h. Scale = 2 mm. c) Expansion rates of the Bamy and Pcl leading edges and distance between both strains during the interaction. The orange line represents the Pcl leading edge, and the green and black lines represent Bamy leading edges 1 and 2, respectively. The blue area represents the distance between the two populations during the entire interaction. d) CLSM time-course experiment of the interaction area using propidium iodide (PI) to identify cell death. Bamy is fluorescently labelled with YFP, and PcI is observed in red. Scale = 40 µm.

Time-course analysis of the Pcl-Bamy interaction in King’s B medium suggested mutual antagonism (Figure 1a bottom), an idea that we explored at the cellular level via time-lapse microscopy analysis of their interaction (Figures 1b-c and Suppl. Movie 1). At the initial stages of colony growth, the expansion rate of Bamy was 5-fold faster than that of Pcl (Figure 1c). After 9 hours of interaction, Bamy growth was arrested, and the leading Pcl-proximal edge showed initial signs of cell death (Figure 1b and d) while Pcl maintained the same displacement rate. After 30 hours, Pcl arrived at the “death zone” of the Bamy colony and progressively reduced the expansion rate, concomitant with a second wave of cell death in the inner areas of the Bamy colony (which was not in physical contact with Pcl) (Figure 1c). The interaction stabilized after 60 hours, and no further inhibition of Bamy colony growth or Pcl expansion was observed. At later time points (90 h to 120 h), small colonies of Bamy grew in the inhibition area (macroscopic images in Figure 1a – 96 h and Figure 4a), and spontaneous clones emerged from the leading edge of the original Pcl colony, which were capable of reaching the inhibition zone (Figure 1b – 120 h and Figure 4a). These results were consistent with our initial hypothesis of mutual exclusion between the two species. In addition, the colony behavior at cellular level and the particular emergence of spontaneous mutants in each strain collectively suggested that Pcl might secrete bactericidal molecules active against Bamy, while bacteriostatic molecules from Bamy seemed to be more relevant in this interspecies interaction.

### Pcl uses the two-component system GacA-GacS to atomize its antimicrobial activity

The previous findings furthered the study of the interaction at two different stages: short-term (24 h to 48 h), to delineate the chemical interplay mediating the antagonism between the two strains; and long-term (96 h to 144 h), to define the genetic changes that lead to their adaptation and coexistence in this adverse chemical environment. We previously reported the relevance of extracellular matrix-related genes in the interaction of *Bacillus* with *Pseudomonas* in LB medium^15^. Thus, we first analyzed gene expression changes at 24 hours to determine whether or not biofilm-related pathways were differentially expressed and to seek putative antimicrobials potentially induced during the interaction (Figure 2a-b, Suppl. Figures 2-4, Suppl. Table 1 and 2). The number of differentially expressed genes in Pcl (304 induced and 59 repressed, p-value <0.05) was larger than that in Bamy (73 induced and 58 repressed, p-value <0.05) (Suppl. Figure 2b). KEGG pathway and GO term analyses showed that pathways related to central metabolism and the cell membrane were mostly altered in Pcl (Figure 2a, Suppl. Figures 2c and 3a). Specifically, we found induction of: i) secondary metabolites, e.g., achromobactin or other gene clusters predicted to participate in the production of secondary metabolites according to AntiSmash^25^; and ii) the type II secretion system (T2SS) and many efflux pumps. Interestingly and contrary to the interaction between *B. subtilis* and Pcl in LB medium^15^, the expression of the T6SS was downregulated, most likely due to the absence of close cell-to-cell contact (Figure 1a). Other downregulated genes were dedicated to the synthesis of the siderophore pyochelin and the antimicrobial pyrrolnitrin (Suppl. Table 1 and Figure 2c). In Bamy, extracellular matrix- and sporulation-related genes were not differentially expressed. The main pathways overexpressed were related to: i) sulfur and nitrogen metabolism and the phosphotransferase system (PTS) (Figure 2b); and ii) synthesis of the antimicrobials bacillaene and difficidin (Suppl. Table 2). Histidine metabolism (Figure 2b) and many of the genes related to membrane proteins and the glycerol uptake system were mostly repressed (Suppl. Table 2 and Suppl. Figures 3b-4).

**Figure 2.**
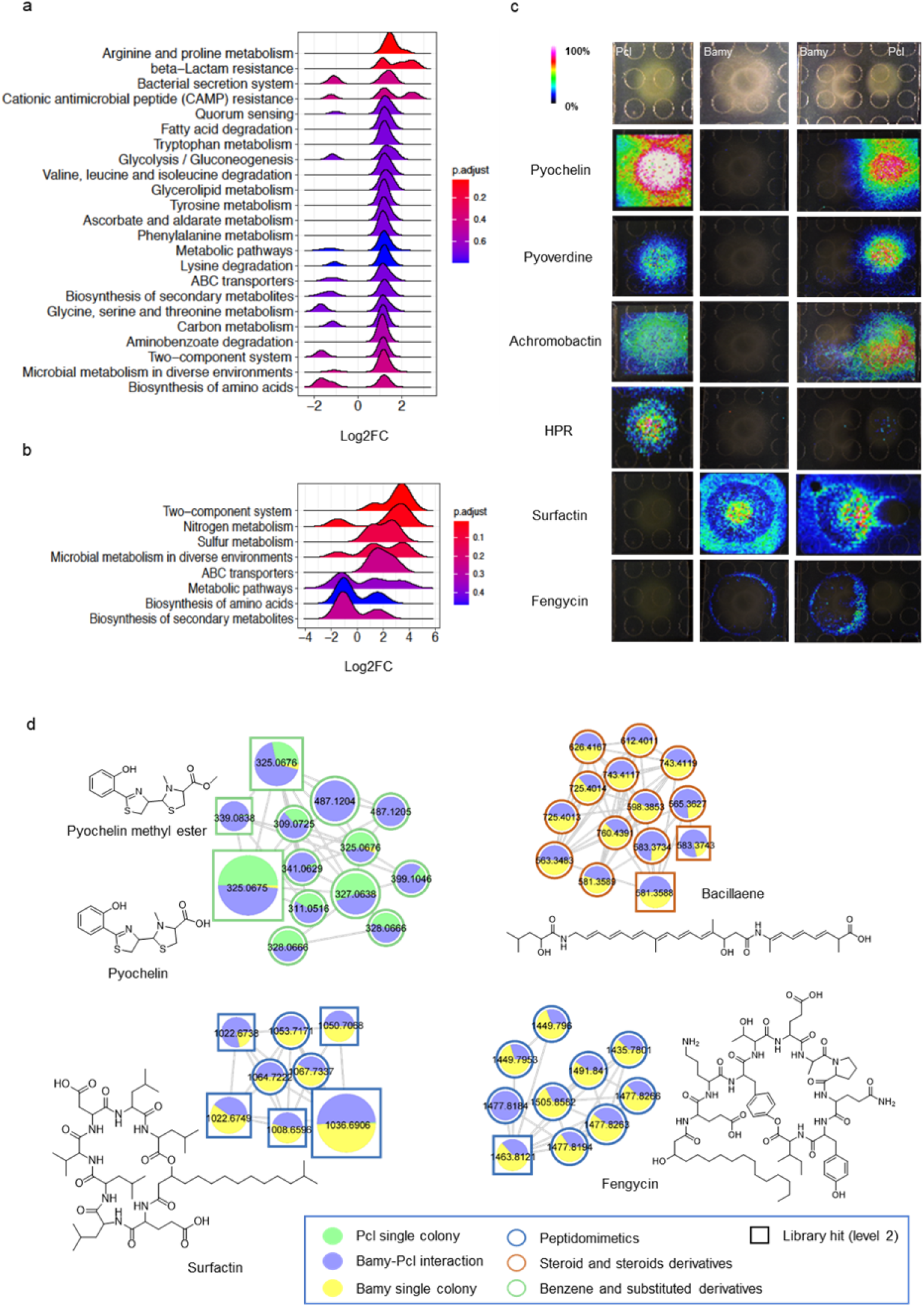
Production and spatial distribution of putative antimicrobials by Pcl and Bamy. a), b) KEGG pathways overexpressed in a) Pcl and b) Bamy after interaction for 24 h. c) MALDI-MSI heatmaps showing the spatial distribution of siderophores (pyoverdine, pyochelin, and achromobactin) and other secondary metabolites (HPR, surfactin, and fengycin) produced by Pcl and Bamy growing alone or in interaction. d) Molecular families of secondary metabolites detected from Pcl and Bamy. The chemical structures of the annotated features are based on spectral matches in GNPS libraries representative of specific molecular families. Border colors indicate ClassyFire classification. The sizes of the compounds are directly related to their abundance in the metabolome. Squares indicate a library hit level 2 through GNPS, and circles indicate unknown compounds based on GNPS searches.

**Figure 3.**
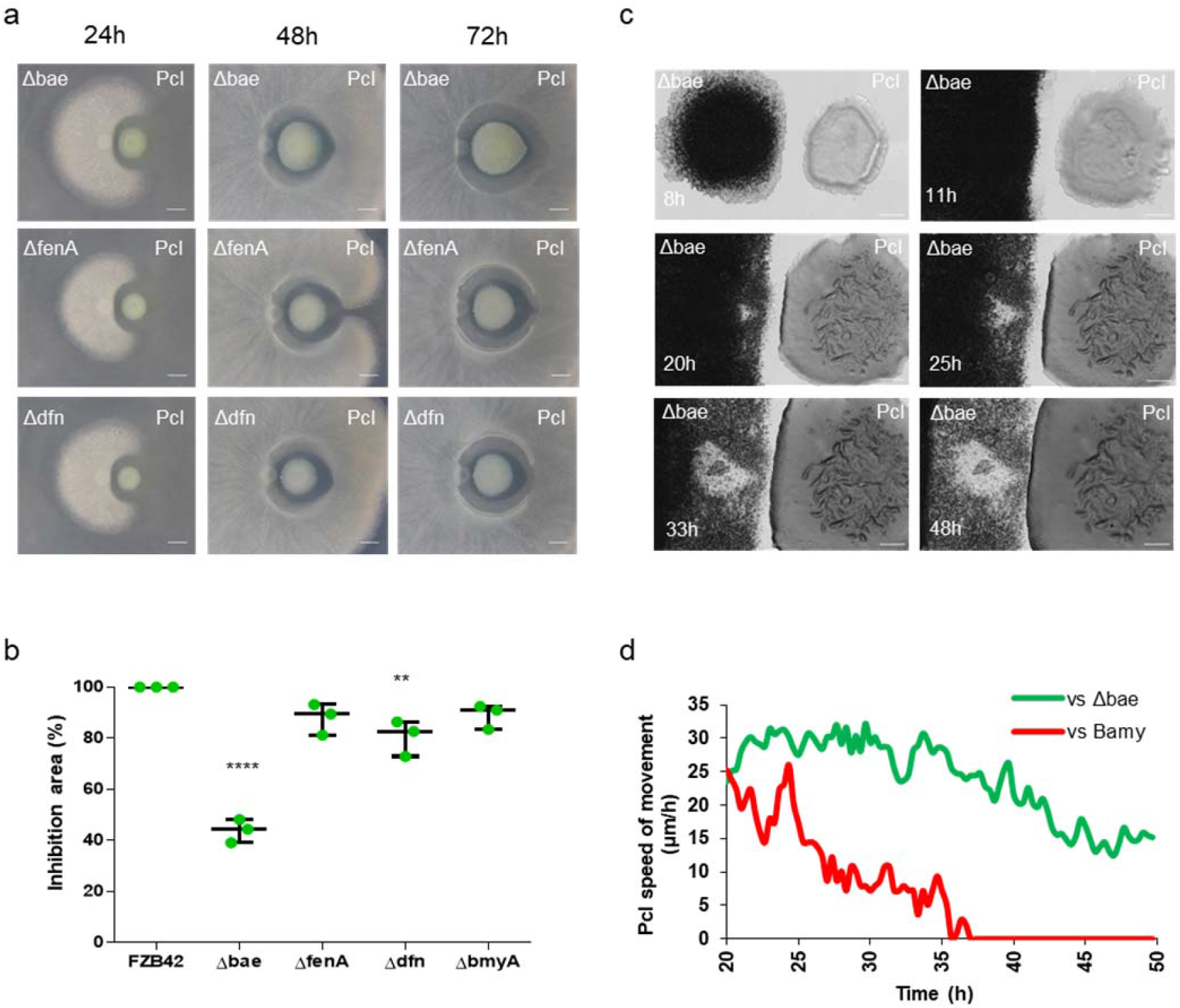
Bacillaene is a bacteriostatic compound produced by Bamy that arrests Pcl growth. a) Time-course pairwise interactions between Pcl and Bamy mutants in secondary metabolites bacillaene (Δbae), fengycin (ΔfenA), and difficidin (Δdfn) on King B medium at 24, 48, and 72 h. Scale = 5 mm. b) Measurement of the inhibition area percentage between Pcl and Bamy mutants at 72 h. Mean values of three biological replicates are shown, with error bars representing SEM. \*\**p*-value < 0.01; \*\*\*\**p*-value < 0.0001 (*t*-test). c) Time-lapse microscopy of the pairwise interaction between Pcl and Δbae over 48 h. Scale= 2 mm. d) Expansion rates of the Pcl leading edges in the interaction with WT Bamy (red line) and Δbae (green line) from 20−50 h.

In light of these results, we analyzed the presence and distribution of metabolites produced by each strain using matrix-assisted laser desorption ionization imaging mass spectrometry (MALDI-MSI) and liquid chromatography-tandem mass spectrometry (LC-MS/MS) non-targeted metabolomics along with the detection of putative inhibitory molecules (Figure 2c-d, and Suppl. Table 3). Although differentially expressed at the transcriptional level (Suppl. Table 1), the siderophores achromobactin, pyochelin, and pyoverdine were distributed around the Pcl colony in the inhibition area and in the inner colony during the interaction (Figure 2c-d). The fractionation analysis of Pcl cell-free supernatants confirmed that fractions containing pyochelin methyl ester (*m/z* 339.08), uniquely detected in the interaction (Figure 2d), HPR (*m/z* 237.18), or two additional unidentified molecules were the most potent inhibitors of Bamy growth (Suppl. Table 3). We were, however, unable to detect achromobactin (*m/z* 592.16) in any of the fractions tested. Unexpectedly, null single pyochelin mutant or even double and triple mutants in the remaining siderophores (achromobactin and pyoverdin) still antagonized Bamy at comparable levels to wild-type (WT) Pcl (Suppl. Figure 5). Diverse lines of evidences also discounted the involvement of HPR in the antagonistic interaction: i) downregulation of gene expression during the interaction; ii) lack of diffusion of the molecule from the Pcl colony as revealed by MALDI-MSI (Figure 2c); and iii) antagonistic activity of a null HPR mutant comparable to the WT (Suppl. Figure 5). Mutants in the T2SS, the most overexpressed efflux pump, and the other non-ribosomal peptide synthetase (NRPS) clusters predicted by AntiSmash still retained comparable antimicrobial activity to WT Pcl (Suppl. Figures 5-6). Untargeted metabolomics and feature molecular networking^26^ using the GNPS platform^27^ and MALDI-MSI experiments showed two additional molecular families produced exclusively by Pcl or primarily during the interaction with a distribution pattern consistent with putative involvement in the antagonism (Suppl. Figure 7a-b). *In silico* analysis with MolNetEnhancer and network annotation propagation (NAP)^28–30^ led us to classify these molecules as glycerophosphoethanolamines and benzene derivatives at the subclass level^30^. The complementary use of SIRIUS/ZODIAC^31^, CSI:FingerID^32^, and CANOPUS^31^ predicted molecular formulas, compound class, and putative chemical structure based on *in silico* MS/MS fragmentation trees. The results obtained for both molecular families were consistent with the MolNetEnhancer results and confirmed the presence of both phosphoethanolamines and benzene derivatives (Suppl. Figure 7a-b); however, the lack of more information on these molecules preclude us from reaching more definitive conclusions.

The combination of metabolomics and functional genetics suggested that Pcl antagonism is multifactorial rather than based on single and known antimicrobials. Thus, we built a library of transposon random mutants to alternatively identify putative candidates involved in this inhibitory activity (Suppl. Figure 8). Clones with no antimicrobial activity shared mutations in the gene RS_08425, which encodes GacS, a member of the GacA-GacS two-component system. Knock-out mutants in other candidate genes did inhibit antagonism. Pairwise interactions and time-lapse microscopy analyses of ΔGacS and Bamy showed no inhibition between the two strains and even overgrowth of Bamy on the leading edge of a ΔGacS colony (Suppl. Figure 7c-d and Suppl. Movie 2). Overall, we concluded that Pcl antagonism is more complex than anticipated and most likely involves several secondary metabolites and other metabolic derivatives, all under control of the environment-sensing GacA-GacS two-component system.

### The offensive actions of Bamy rely on the bacteriostatic compound bacillaene

Transcriptomic analysis revealed overexpression of bacillaene and difficidin by Bamy as a consequence of its interaction with Pcl (Suppl. Table 2), and non-targeted metabolomics confirmed bacillaene overproduction. MALDI-MSI analysis added surfactin and fengycin to the group of putative antimicrobials of Bamy that might mediate its antagonistic interaction with Pcl (Figure 2c-d). Fengycin and bacillomycin are well-known antifungal compounds with little or null antibacterial activity, and accordingly, pairwise interactions between strains single mutant for either of these molecules (ΔfenA or ΔbmyA) and Pcl provided comparable antagonistic results to that of WT Bamy (Figure 3a-b and Suppl. Figure 9). A surfactin mutant (ΔsrfA) showed reduced colony expansion (an expected finding) but antagonized Pcl as comparably to WT Bamy. The size of the inhibition area was, however, mildly reduced in the interaction with a difficidin mutant (Δdfn) (Fig. 3a) and strongly diminished (60% compared to the interaction with WT Bamy) in the absence of bacillaene (Δbae) (Figure 3b). Two additional findings supported the inhibitory contribution of bacillaene to the interaction: i) the disappearance of most of the inhibition halo after 72 hours; and ii) the larger size of the Pcl colony in comparison with those of the other interactions (Figure 3a). Time-lapse microscopy experiments confirmed that, indeed, the larger size of the Pcl colony was due to growth beyond the area initially colonized by the Δbae colony (Figure 3c and Suppl. Movie 3). A comparison of the speed of movement clearly showed that Pcl expanded constantly at 25-30 µm/h in the interaction with Δbae (Figure 3d), but that it completely ceased after 35 hours of interaction with WT Bamy. These results provide support for a defensive role of bacillaene produced by Bamy against the advance of Pcl; furthermore, the lack of massive cell death at the front line of the Pcl colony also led us to confirm that bacillaene exerts a bacteriostatic effect ^33^.

### Mutations in FusA overcome bacillaene-mediated antagonism

As revealed earlier via time-lapse microscopy analysis, Bamy colonies spontaneously emerged in the inhibition zone, and clones grew from the Pcl colony when the interaction stabilized after 90 hours (Figure 1b and Figure 4a and Suppl. Movie 1). To elucidate the genomic changes driving this microbial adaptation, we sequenced the genomes of three spontaneous Pcl clones and seven spontaneous Bamy clones. Comparison of the Pcl clones showed a variety of point mutations at different loci, and recurring changes were found in the locus PCL1606_RS29735 locus, which encodes translation factor elongation factor G (FusA) (Table 1). No inhibition halos were observed at late time points during pairwise interactions between these Pcl clones and WT Bamy, a pattern similar to that of the interaction between WT Pcl and Δbae (Suppl. Figure 10a). These results show that the speed of movement of Pcl clones, recorded via time-lapse microscopy experiments (Figure 4b and Suppl. Movie 4), was constant during the interaction and similar to that during the interaction between WT Pcl and Δbae, although forward movement decelerated relative to that observed in the complete absence of bacillaene (Figure 4c-d).

**Figure 4.**
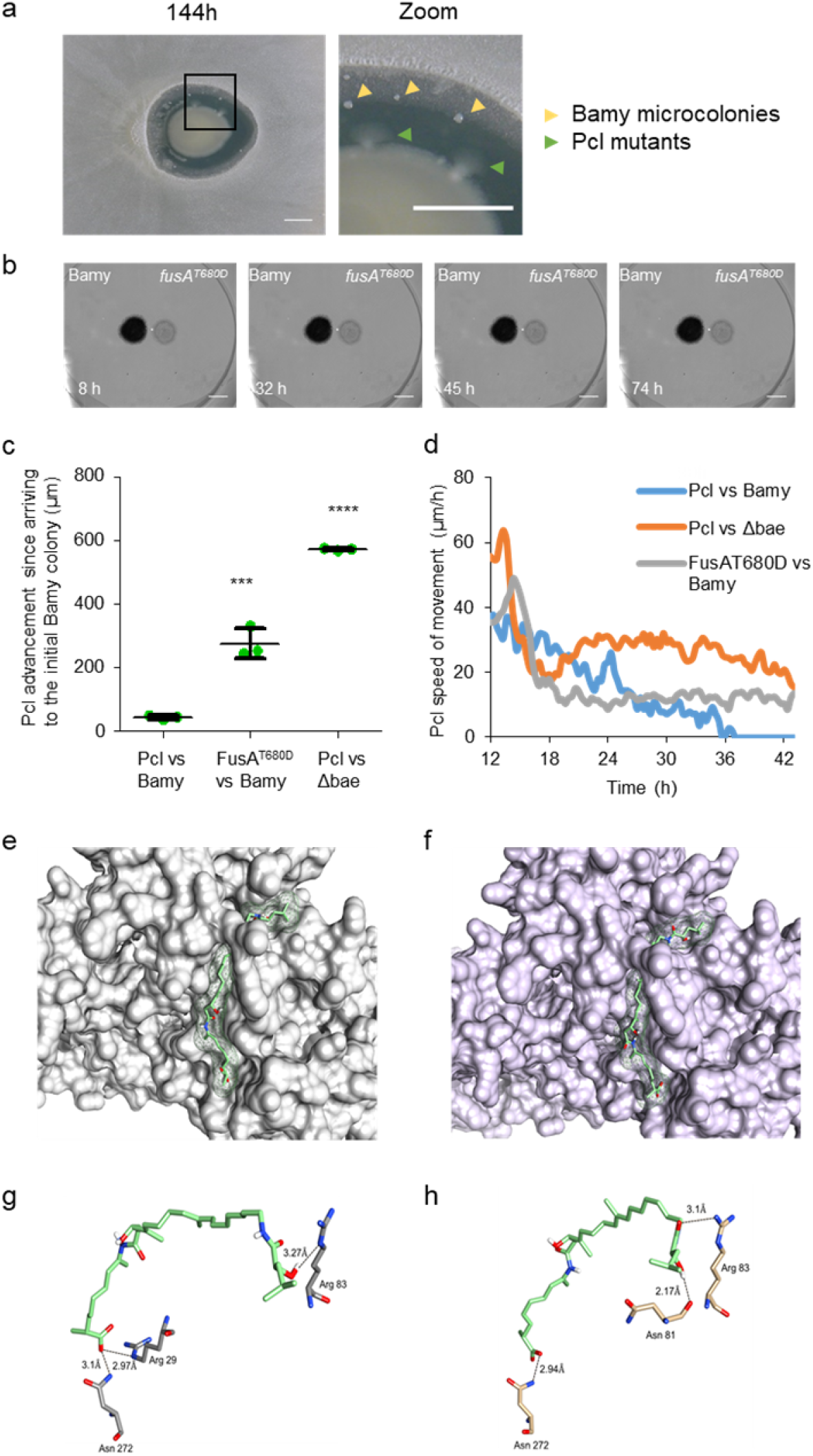
Mutations in Pcl FusA confer bacillaene resistance. a) Pairwise interaction between Pcl and Bamy after interaction for 144 hs. Magnification of the selected area permits visualization of Bamy microcolonies (yellow triangles) and Pcl subzones (green triangles) growing in the inhibition area. Scale = 5 mm. b) Time-lapse microscopy of the pairwise interaction between FusA^T680D^ and Bamy after 74 hs. Scale= 2 mm. c) Measurement of the Pcl colony movement within 24 h of their arrival at the initial position of a Bamy colony. Mean values of three biological replicates are shown, with error bars representing SEM. *** *p*-value < 0.001; **** *p*-value < 0.0001 (*t*-test). d) Expansion rates of the WT Pcl leading edges during interaction with WT Bamy (blue line), Δbae (orange line), and the FusA^T680D^ leading edge expansion rate during interaction with Bamy from 12−43 h. Molecular docking between bacillaene and e) FusA or f) FusA^T680D^ reveals a putative binding site formed by Arg29, Arg89 and Asn272. g) and h) Distances measured between the bacillaene molecule and side chains of these residues in FusA and FusA^T680D^.

**Table 1.** Alleles of fusA identified in Pcl spontaneous mutants.

| Allele | Nucleotide Change | Amino acid Change | Resistance to Bamy |
| --- | --- | --- | --- |
| *fusA*<sup>T680D</sup> | g.2038A>G | p.T680D | + |
| *fusA*<sup>T680D</sup> | g.2038A>G | p.T680D | + |
| *fusA*<sup>K366N</sup> | g.1098G>T | p.K366N | + |

Previous studies have demonstrated that *Staphylococcus aureus* FusA is the target of the antibiotic fusidic acid^34,35^. In addition, FusA has also been revealed to be involved in tolerance of the aminoglycoside antimicrobials gentamicin, amikacin, and tobramycin by *P. aeruginosa*^36^. FusA is highly conserved, and its three-dimensional structure has been elucidated, enabling *in silico* molecular docking studies of the WT protein or mutated proteins in the presence and absence of bacillaene. Molecular modelling of WT FusA and FusA^T680D^ showed slight structural and conformational changes in the most probable bacillaene binding pocket (Suppl. Movie 5). Molecular docking of bacillaene and WT FusA resulted in 48 identifiable clusters in eight different sites in the protein. The top-score cluster exhibited higher full fitness and lower free energy values of -5458.2 and −7.8 kcal mol^-1^, respectively, compared with those of other potential binding sites. In its interaction with WT FusA, bacillaene formed one hydrogen bond with the secondary amine of Arg29 (2.97 Å), one with the secondary amine of Arg89 (2.1 Å), and one with the primary amine of Asn272 (3.101Å), suggesting a favorable binding site (Figure 4e and g). However, molecular docking of bacillaene and FusA^T680D^ resulted in the identification of 9 clusters in two different sites in the protein. The top-score cluster exhibited higher full fitness and free energy values of −2187.4 and −7.6 kcal mol^-1^, respectively, compared with those of WT FusA. In this interaction, three hydrogen bonds were predicted: one hydrogen bond with the hydroxyl group of Asn81 (2.17 Å), one with the secondary amine of Arg83 (3.10 Å), and one with the primary amine of Asn272 (2.94 Å) (Figure 4f and h), suggesting a lower affinity for bacillaene with the mutant FusA and, thus, reduced inhibitory activity.

Based on our model, we hypothesized that mutations in FusA would also decrease its affinity for fusidic acid, thus promoting higher tolerance to this molecule (Suppl. Figure 10b-c). The top-score cluster obtained via molecular docking of WT FusA with fusidic acid revealed full fitness and free energy values of −3257.2 and −8.7 kcal mol^-1^, respectively, with five hydrogen bonds predicted: one with the hydroxyl group of Glu98 (2.21 Å), one with the hydroxyl group of Thr89 (2.18 Å), two with the primary and secondary amines of Arg101 (3.027 Å and 3.13 Å), and one with the hydroxyl group of Thr402 (2.98 Å), suggesting a favorable binding site. The same analysis with the FusA^T680D^ model exhibited a top-score cluster with full fitness and free energy values of −2180.025 and −6.98 kcal mol^-1^, respectively. Three hydrogen bonds were predicted: one with the secondary amine of Arg101 (2.54 Å) and two with Thr402 (one with the hydroxyl group (2.51 Å) and one with its secondary amine (2.33 Å)). These results are consistent with those of bacillaene, providing strong evidence of its mechanism of action. Interestingly, we also noticed that mutations in FusA provided protection against other antibiotics that target the translation machinery. Minimal inhibitory concentration (MIC) experiments using the aminoglycoside antibiotics kanamycin and gentamicin, which target ribosomal subunits, confirmed that these single amino acid mutations in FusA resulted in higher levels of resistance, i.e., two- and ten-fold, respectively (Table 2).

**Table 2.**
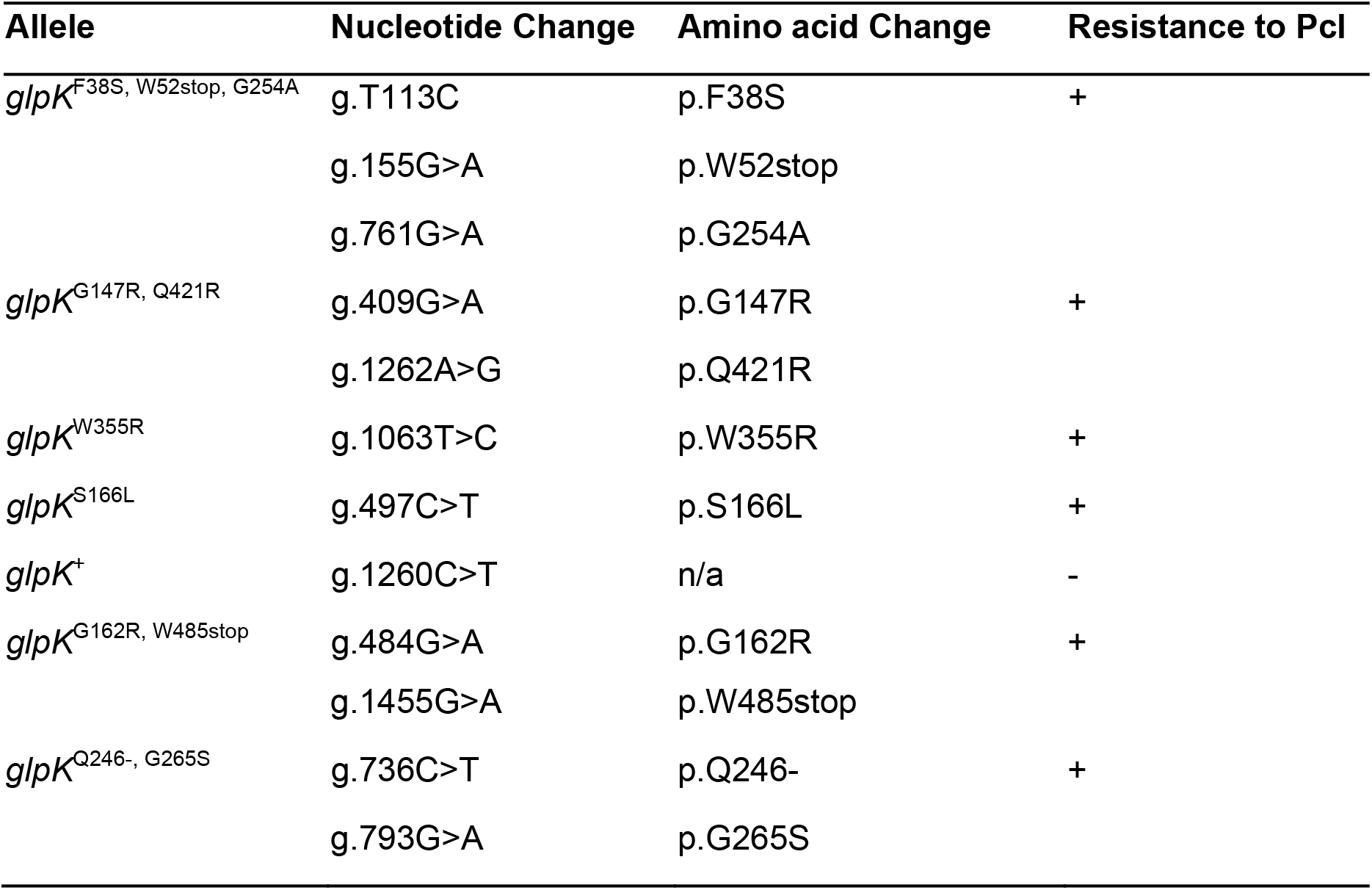
Alleles of glpK identified in Bamy mutants.

Based on these results, we conclude that the antimicrobial activity of bacillaene stems from its inhibition of protein translation via direct interaction with FusA and that point mutations in the protein, that preserve its function, provide resistance to bacillaene and other unrelated protein translation inhibitors.

### GlpK promotes pleiotropic cellular changes and unspecific antibiotic resistance

Genome sequence analysis and comparison from seven Bamy isolates showed only a few mutations at multiple loci (Figure 5a). Mutations in multiple residues were only found in *glpK* (Table 3). Two of the clones showed mutations that result in premature stop codons, one mutation was silent (no amino acid substitution), and the remaining mutations accumulated one- or two-nucleotide changes that resulted in amino acid substitutions. Phenotypically, these clones expanded less than the WT Bamy in King’s B medium and in glycerol-supplemented LB medium (Suppl. Figures 11 and 12), demonstrating the relevance and direct involvement of glycerol metabolism in the expansion of these colonies. In interactions with Pcl, the *glpK* mutants reached the Pcl colony, although initial signs of cell death were again observed at the Pcl-proximal colony edge (Figure 5b top). To confirm that these findings were associated with the loss of GlpK function, we constructed a *glpK* null strain via deletion of *glpK*, and this strain phenotypically mirrored the isolated spontaneous clones (Figure 5b bottom). Examination of the interaction between GlpK^S166L^ and WT Pcl at the cellular level via time-lapse microscopy experiments demonstrated partial inhibition of *Bacillus* growth at the Pcl-proximal leading edge of the colony (Figure 5c and Suppl. Movie 6). However, notable differences in the interaction between WT Bamy WT and Pcl included: i) lack of massive cell death in the leading edge of GlpK^S166L^, which permitted physical contact between the two colonies (Figure 5c); and ii) absence of regression of a secondary leading edge of GlpK^S166L^ strain after 40 hours of interaction (Suppl. Figure 13a).

**Figure 5.**
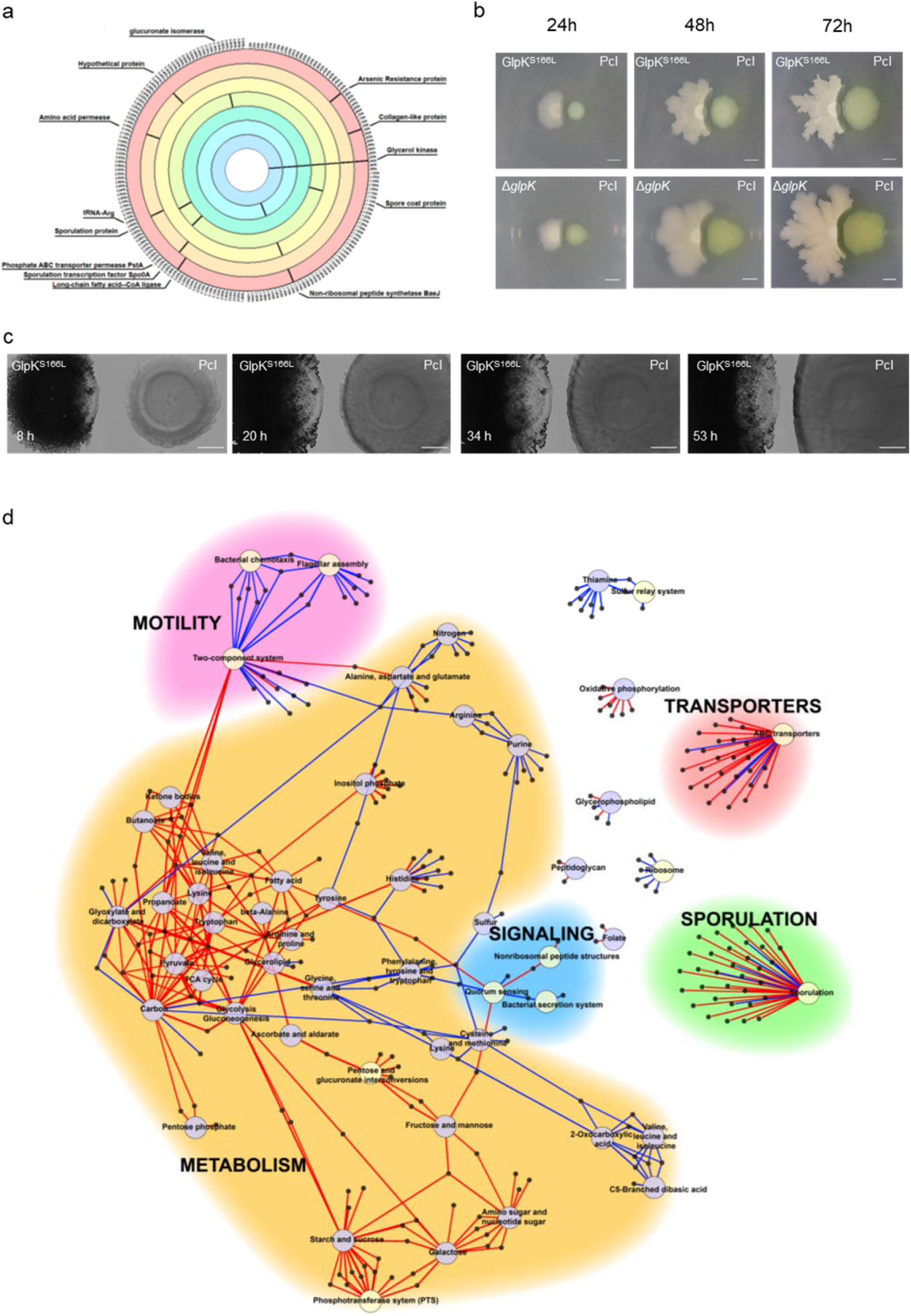
Pleiotropic changes protect Bamy *glpK* mutants from antibiosis of Pcl. a) Circular representation of the point mutations found in the genomes of Bamy clones with *glpK* mutations. Each ring represents one sequenced Bamy clone. b) Time-course pairwise interactions between Pcl and GlpK^S166L^ (top) and ΔglpK (bottom) at 24, 48, and 72 h. Scale = 5 mm. c) Time-lapse microscopy of the pairwise interaction between Pcl and GlpK^S166L^ over 48 h. Scale= 2 mm. d) Differentially expressed genes between Bamy and GlpK^S166L^ at 24 h clustered into different metabolic pathways. Larger circles indicate the main KEGG pathways, which are surrounded by arrows pointing to smaller circles that represent the differentially expressed genes. The color of the arrows indicates induction (red) or repression (blue). The color of the circles differentiates pathways involved in metabolism (light blue) from those not involved (light yellow).

We next wanted to determine the genetic basis of the phenotypic changes in the *glpK* mutants and the highest tolerance to Pcl antagonism. Comparative transcriptomic analysis of the GlpK^S166L^-mutant strain and WT Bamy revealed a vast number of differentially expressed genes related to central metabolism and sugar uptake (Figure 5d and Suppl. Table 5). As expected from *glpK* mutants incapable of metabolizing glycerol, the main carbon source in King’s B medium, we found overexpression of PTS and the pathways involved in the uptake of galactose, starch and sucrose, fructose, or mannose (Suppl. Figure 13b). Additionally, the TCA cycle, pentose phosphate pathway, and fatty acid metabolism were also induced, while the expression levels of genes involved in amino acid biosynthesis and metabolism were only mildly activated or repressed (Figure 5d and Suppl. Table 5). Reduced swarming motility (Suppl. Figure 13c) reflected repression of motility- and quorum sensing-related genes. The expression levels of genes associated to secretion systems and sporulation were induced compared with their levels in WT Bamy.

Complementary metabolomic analysis revealed a pronounced reduction of surfactin production, which is likely associated with the reduced swarming motility of *glpK* mutants, slight changes in bacillaene production, and increased fengycin production (Suppl. Figures 14-15). Corroborating the transcriptomic data, metabolomics analysis also indicated important changes in the metabolic pathways involved in fatty acid metabolism, specific changes in the composition of phosphoethanolamines of Bamy cells (Figure 5d, Figure 6a and Suppl. Table 5), and the absence of or decreases in the levels of glycerophosphoethanolamines and benzene derivatives (Suppl. Figure 16a), which are candidate inhibitory compounds produced by Pcl.

**Figure 6.**
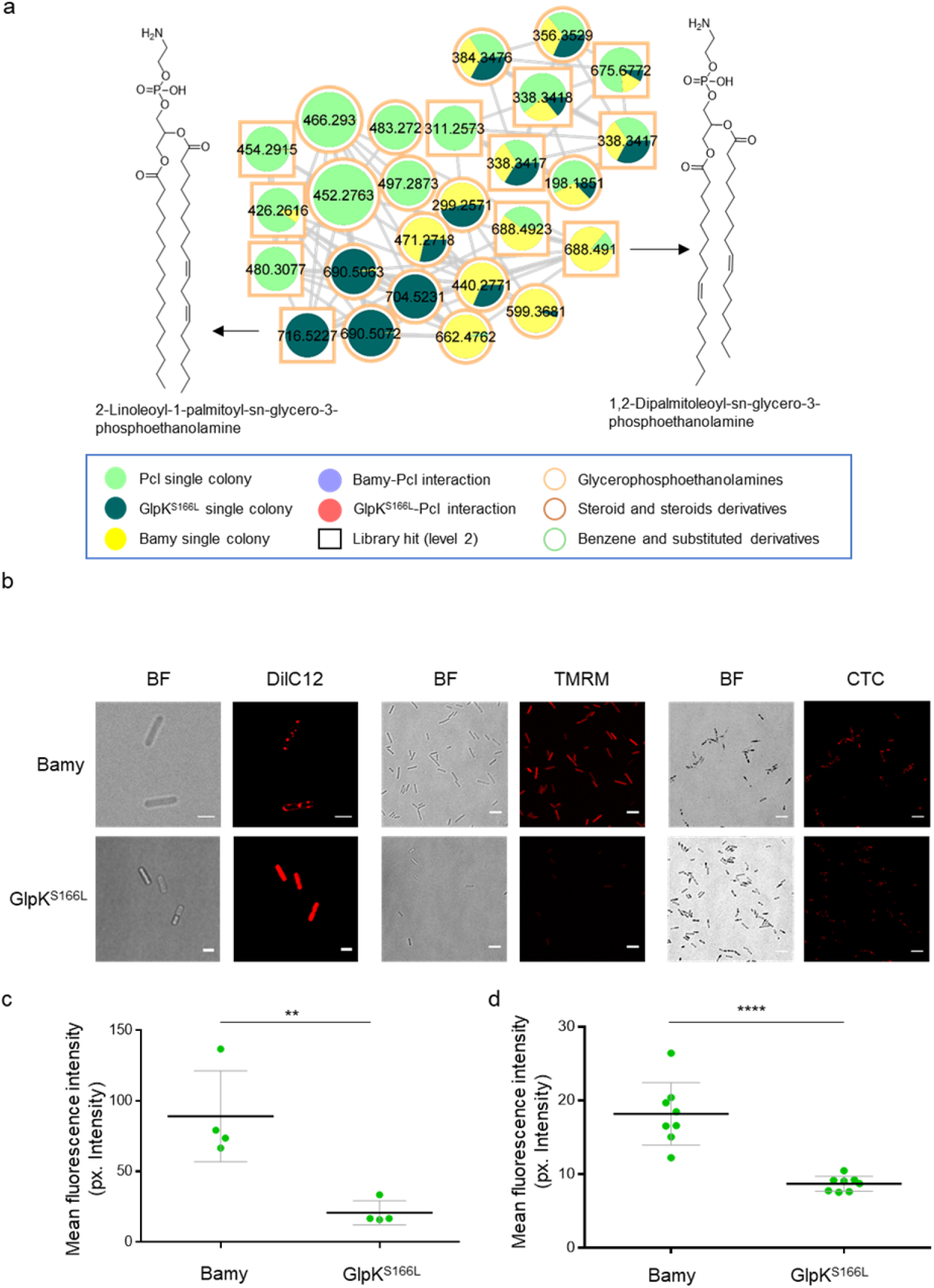
Mutations in *glpK* provoke changes in membrane rigidity and permeability. a) Glycerophosphoethanolamine cluster showing modifications in the abundance of different glycerophosphoethanolamine compounds between Pcl, Bamy, and GlpK^S166L^. Chemical structure of annotated features are based on spectral matches to GNPS libraries, as representative of these molecular families. Border color indicates ClassyFire classification. The sizes of the compounds are directly related to their abundance in the metabolome. Squares indicate a library hit level 2 through GNPS, while circles indicate unknown compounds based on GNPS. b) Membrane staining of Bamy and GlpK^S166L^ with (left panel) DilC12 dye to analyze differences in membrane fluidity (scale = 2 µm), (middle panel) TMRM straining for membrane potential analysis (scale 5 µm), and (right panel) CTC staining to measure respiration changes (scale = 5 µm). c) Quantification of the TMRM signal in Bamy and GlpK^S166L^. d) Quantification of the CTC signal in Bamy and GlpK^S166L^ cells. Mean values of three independent replicates are shown. Error bars represent the SEM. For each experiment and sample, at least three fields-of-view were measured. Statistical significance in the TMRM and CTC experiments was assessed via two-tailed independent t-tests at each time-point (\*\**p-value* < 0.01).

We predicted that changes in the overall fatty acid content of *glpK* mutant cells could alter the functionality of cell membrane lipids, possibly underlying the higher protection against Pcl chemical aggression. Confocal microscopy analysis of cultures treated with the membrane dye DilC12 revealed the presence of highly fluorescent foci, which correspond to regions with increased fluidity, distributed along the membrane of WT Bamy cells. However, the membranes of GlpK^S166L^ cells stained uniformly with no signs of patches of dye accumulation (Figure 6b left and 6c), which indicated higher rigidity and, thus, reduced permeability. We also observed significant decreases in membrane potential and the respiratory activity of GlpK^S166L^ cells in tetramethylrhodamine methyl ester (TMRM) and 5-cyano-2,3-ditolyl-tetrazolium chloride (CTC) experiments, respectively (Figure 6b center and right, Figure 6d, and Suppl. Figure 16b). These results reveal that GlpK confers an unexplained tolerance to antibacterial compounds produced by Pcl; thus, mutations in this gene are selected as the most favorable evolutionary trait in the face of diverse chemical aggression^37,38^.

## Discussion

Bacterial interactions are normally considered from an antagonistic perspective, and ongoing effort is dedicated to identifying the specific inhibitory metabolites. However, bacterial interactions are diverse, and the involvement of specific mechanisms largely relies on cell-to-cell distance and the environment (Figure 7). In close cell-to-cell interactions, injection structures, e.g., the T6SS^15,39–41^, ensure the efficient delivery of the inhibitory compounds, while in long-distance interactions, diffusible molecules are more important^42,43^. Inhibitory molecules are not always produced or concentrated at lethal levels, and alternative modes of action that induce changes in gene expression patterns or in modulation of the behavior of the competitor have emerged^44,45^. Thus, interspecies interactions are not classified as beneficial or detrimental based on specific time frames; however, these interactions represent a fascinating evolutionary process where competitors attempt to adapt and survive in the presence of other species in the same ecological niche. In this work, we provide substantial insight into the evolution of interspecies interactions by studying the mechanism of the chemical interplay between Pcl and Bamy, two plant-beneficial bacteria^2,16,20,22^. Understanding of this interaction is also important from a biotechnological perspective, given that their beneficial effects can be potentiated or inhibited depending on how the strains interact in the short term and adapt and evolve to ensure subsistence and/or coexistence in the long term.

**Figure 7.**
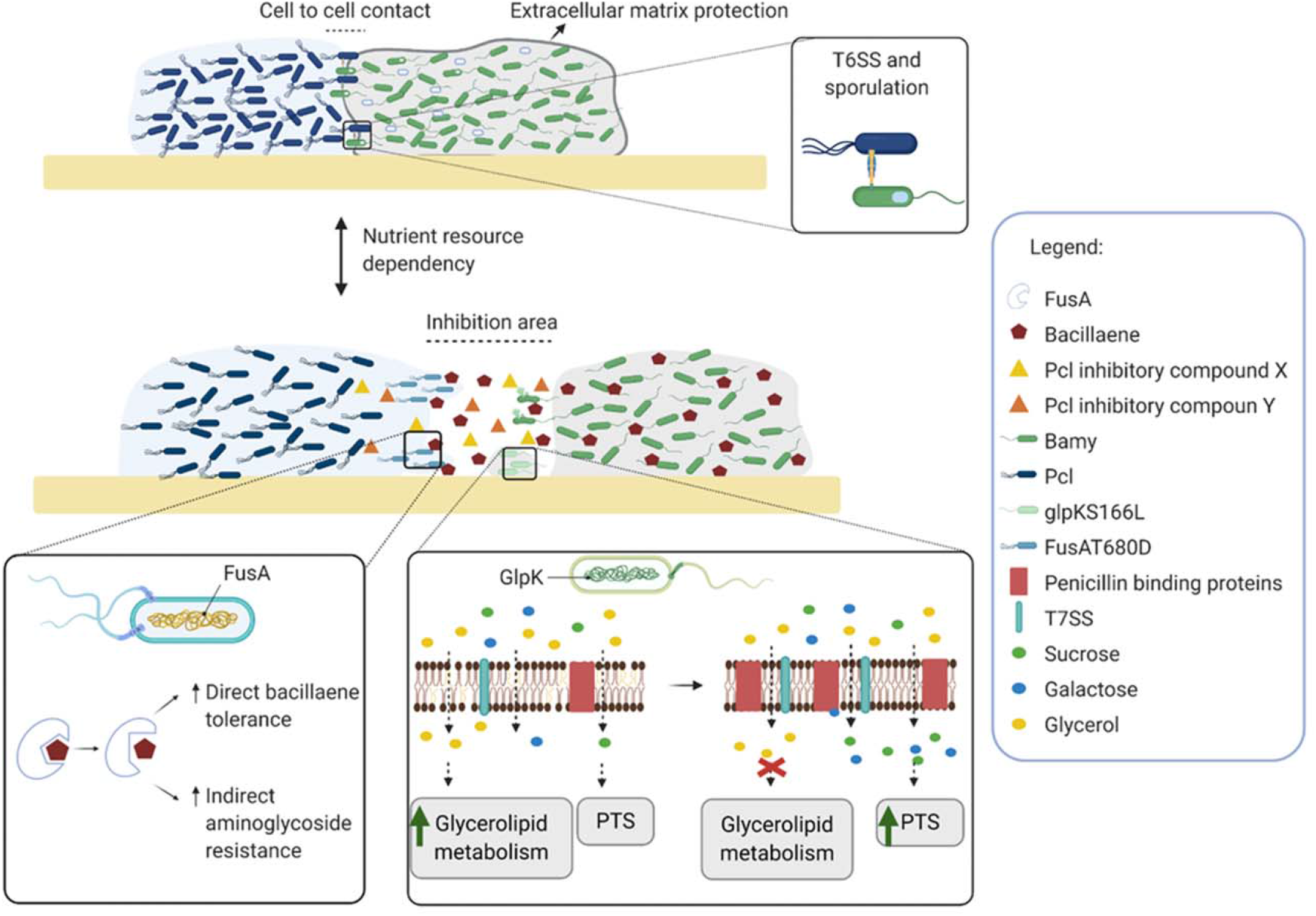
Antagonism or co-existence is timely and spatially determined by multiple bacterial factors and nutrient availability. Schematic representations of the interaction between *Bacillus* and *Pseudomonas* depending on critical factors, e.g., nutrients and distance. (Upper part) When bacterial populations are in contact, T6SS, sporulation, and the extracellular matrix play important roles in the interaction. (Bottom part) When populations are physically separated, the production of secondary metabolites is critical for the evolution of the interaction. Mutations in *Pseudomonas fusA* permit adaptation to *Bacillus-*produced bacillaene, therby increasing resistance to bacillaene and aminoglycoside antibiotics. Mutations in *Bacillus glpK* provoke a wide array of transcriptional and metabolic changes, an increase in bacterial membrane rigidity, and a reduction membrane potential, accompanied by overexpression of penicillin-binding proteins and T7SS as well as changes in lipid composition, resulting in increased antibiotic tolerance.

In a previous study we demonstrated the relevance of the *Bacillus subtilis* extracellular matrix in the avoidance of colonization by *Pseudomonas chlororaphis* cells. The same experiment in King’s B medium, however, indicated that diffusible molecules were more relevant than the extracellular matrix in modulating the interaction dynamics (Figure 1a and Suppl. Figure 1). Time-lapse microscopy allowed us to better characterize the sequence of events that progressively shift the interaction from antagonism to co-existence over a period of 4−5 days. Unexpectedly, we could not restrict the antagonism of Pcl towards Bamy to a single compound. Our current results show that the antagonistic effect of Pcl in the short term is most likely multifactorial and controlled by the GacA-GacS two-component system. This environmental sensing system regulates the expression of genes required for the production of secondary metabolites and extracellular enzymes that influence motility, iron acquisition, biofilm formation, and other aspects of primary metabolism in closely related *Pseudomonas* strains^46–49^. Contrary to the combinatorial inhibition displayed by Pcl, we identified bacillaene as the most important molecule produced by Bamy that contributes to the initial antagonistic interaction. Bacillaene has been proposed to be a bacteriostatic molecule that protects *Bacillus* from predation by *Myxococcus xanthus*^50^ and kills *Serratia plymuthica*, facilitating its engulfment by *B. subtilis*^51^. Although the arrest of protein synthesis has been suggested as its mode of action, this possibility has not been clarified^33^. In this study, we have demonstrated that the bacteriostatic effect of bacillaene confers protection to the Bamy colony by impeding the expansion of the Pcl colony without massive cell death. Based on our mutational and *in silico* analyses, we concluded that bacillaene’s mechanism of action (similar to that of fusidic acid^34^) depends on an interaction with elongation factor G (FusA), which leads to translation inhibition. Consistently, a typical evolutionary resistance strategy mediated by modification of the antibiotic’s target, e.g, via point mutations in Pcl *fusA*, reduces the interaction with bacillaene, causing resistance to this molecule and indirectly to aminoglycosides that inhibit protein synthesis by targeting ribosomal subunits (Figure 7).

The antagonism of Pcl based on multiple compounds might explain the fact that predominant spontaneous mutants of Bamy do not rely on a specific antibiotic target but in the gene *glpK* encoding the GlpK. The GlpK enzyme catalyzes the first step in the glycerolipid pathway converting glycerol into glycerol-3-phosphate, a modification that ensures the entrance of this carbon source in the catabolic pathways and promotes the formation of glycerolipids. Mutations of *glpK* have been also found in *Mycobacterium tuberculosis* in response to long antibiotic treatments, however, these are transient and the mechanism behind the antibiotic tolerance remains unexplained^52,53^. Others have speculated that GlpK may have an alternative role as a transcriptional regulator^52^. The notorious transcriptional changes observed upon deletion of *glpK* affect complementary metabolic pathways oriented to the obtaining of energy but also metabolic pathways related with lipid metabolism, motility, sporulation and signaling. These changes are evidence of the complementary role of GlpK. Based on this global deregulation we propose a diversified strategy led by GlpK to overcome antagonism in nutritional competitive scenarios (Figure 7). First, GlpK causes the induction of sporulation and the adaptation to utilizing alternative carbon sources that transiently overcome the presence of toxic molecules produced by competitors. Second, the changes in the lipid composition and properties of the cell membrane leads to reduced permeability and thus entrance of antimicrobials. Third, the induction of multidrug ATP-binding cassette transporters, type VII secretion system (T7SS), and penicillin binding proteins (Figure 7) complementarily create a non-specific physical barrier to the entry of antimicrobial and toxic compounds. Our findings, together with those obtained in human pathogens, highlight an important role of GlpK not only in antibiotic resistance but in bacterial adaptation to interactions in specific niches, from humans to plants.

Taken together we propose a model where the initial antagonism for nutrients and space evolves towards a situation where WT and mutant genotypes seem to coexist in the same niche in an stable association of mixed sub-populations with mutual benefits for species survival^54^. The participation of other bacterial structures (T6SS or ECM) most likely are affected by the spatial effect imposed by the topology or chemical nature of the host, and would complementarily determine the final organization of the bacterial community. We conclude that the appearance of mutant subpopulations with different adaptability strategies is based on the metabolic specialization, and the involvement of chemical interactions finally determines the population size and the evolution towards mixed and stable communities rather than the complete extinction of the competitor.

## Methods

### Strains, media and culture conditions

A complete list of the bacterial strains used in this study is shown in Suppl. Table 6. Routinely, bacterial cells were grown in liquid lysogeny borth (LB) medium at 30 °C (Pcl and Bamy) or 37 °C (*E. coli*) with shaking on an orbital platform. When necessary, antibiotics were added to the media at appropriate concentrations. Strains and plasmids were constructed using standard methods^55^.

### Pseudomonas chlororaphis mutants

Chromosomal deletions of Pcl mutants were performed using the I-SceI method^56,57^ in which upstream and downstream segments of homologous DNA are separately amplified and then joined to a previously digested pEMG vector using Gibson Assembly Master Mix^58^. The oligonucleotide sequences are shown in Suppl. Table 7. The resulting plasmid was then electroporated into Pcl^1606^. After selection for positive clones, the pSEVA628S I-SceI expression plasmid was also electroporated, and kanamycin (Km)-sensitive clones were analyzed via PCR to verify the deletions. The pSEVA628S plasmid was cured via growth without selective pressure, and its loss confirmed by sensitivity to 60 µg ml^−1^ gentamicin and colony PCR screening as described by Martinez-Garcia and de Lorenzo^56^.

### Bacillus amyloliquefaciens mutants

Bamy strains with *glpK* mutations were obtained as follows. Approximately 1500-base pair genomic regions upstream and downstream of the genes of interest were amplified from Bamy FZB42 chromosomal DNA. The two gel-purified double-stranded DNA fragments were linked by a Km resistance cassette and then ligated into pMAD. The linearized plasmids were integrated into the genome of Bamy via double-crossover recombination, yielding the Bamy knockout mutants.

### Construction of fluorescence labeling strains

The fluorescence labeling plasmid pKM008V was constructed for Bamy strains. Briefly, the P_veg_ promoter fragment (300 bp) was extracted from pBS1C3 via EcoRI and HindIII digestion, purified, and cloned into the plasmid pKM008, which was previously digested with the same restriction enzymes. We used P_veg_ as it is considered a constitutive promoter in Bamy. pKM008V was then linearized and transformed into Bamy via natural competence, and transformants were selected by plating on LB plates supplemented with spectinomycin (100 μg ml^−1^).

### Pairwise interactions

Bamy and Pcl strains were routinely spotted 0.7 cm apart on King B agar plates using 2 µl of cell suspension at an OD_600_ of 0.5. Plates were incubated at 30 °C, and were images taken at different timepoints. For confocal microscopy time-course experiments, 0.7 µl of cell suspension was spotted at distance of 0.5 cm onto 1.3-mm thick LB agar supplemented with propidium iodide in 35-mm glass-bottomed dishes suitable for confocal microscopy (Ibidi). Temperature was maintained at 28 °C during the time-course using the integrated microscope incubator. Acquisitions were performed using an inverted Leica SP5 confocal microscope with a 25× NA 0.95 NA IR APO long working distance water immersion objective. Image processing and three-dimensional (3D) visualization were performed using ImageJ/FIJI^59,60^ and Imaris version 7.6 (Bitplane).

Colony time-lapse videos were acquired using a Nikon Ti inverted microscope equipped with DIC brightfield illumination and a 4x Plan NA 0.1 dry objective. A stage-top incubation system with an incorporated digital temperature sensor (Okolab) was used to maintain the temperature at 28 °C. Time-lapse images were acquired using open-source Micromanager Software version 2.0 beta^61^ and a Hamamatsu Orca R2 monochrome camera set to 2 x 2 binning and 4 ms exposure. Mosaic acquisition was performed with a 10% field overlap using the multi-dimensional acquisition module and two-step autofocus (JAF H&P; 1st step size 2.0, 1st step number 1.0; 2nd step size 0.2, 2nd step number 5; threshold 0.02 and crop ratio 0.2). Time-lapse images were typically captured over 2−4 days at 20-minute intervals. Mosaic merging was performed using FIJI and the Grid/Collection stitching plugin^62^. Flatfield correction was performed using the Stack Normalizer plugin (Joachim Walter; https://imagej.nih.gov/ij/plugins/normalizer.html) using a Gaussian-filtered time zero image as reference.

### Generation and screening of mini-Tn5 mutants of Pcl that do not inhibit Bamy

Pcl mini-Tn5 transposon mutants were constructed as described by Matas et al.^63^ with minor modifications. The pool of pUTminiTn5Km1 vectors was transferred from *E. coli* S17 λpir to Pcl via plate conjugation mating as previously described^64^. The constructed random transposition collection consisted of 28 96-well microtiter trays with a total of 2688 Pcl mutants. For screening, a lawn of Bamy was inoculated onto King B solid medium and dried for 15 minutes at room temperature. Next, 1 µl of each Pcl mutants was individually spotted on the plates followed by incubation at 30 °C for 24 hours. Pcl mutants that did not cause an inhibition halo around them were selected. Genomic DNA was extracted from the Pcl mutant strains using the Jet Flex Extraction Kit (Genomed, Löhne, Germany) according to the manufacturer’s instructions. To determine transposon insertion sites, genomic DNA was digested with PstI or XbaI and ligated into pBluescript II SK digested with the same restriction enzyme. Ligation reactions were used to transform DH5α by heat shock^65^, and single Km-resistant colonies were selected. Plasmids were purified and DNA regions flanking transposons were sequenced by Macrogen (Madrid, Spain) using primer P7 and the sequences were analyzed using BLASTn.

### Whole-genome transcriptomic analysis

Single colonies of Pcl, Bamy, and the GlpK^S166L^ mutant were grown overnight on solid LB medium at 30 °C and spotted on King B medium as single colonies or as interactions as previously described for 24 hours. Next, cells were collected and stored at −80 °C. All assays were performed in duplicate. For disruption of single colonies and interactions, collected cells were resuspended in BirnBoim A75 and lysozyme was added followed by incubation at 37°C for 15 minutes. Next, the suspensions were centrifuged, the pellets were resuspended in Trizol, and total RNA was extracted as indicated by the manufacturer. DNA removal was carried out via treatment with Nucleo-Spin RNA Plant (Macherey-Nagel). Integrity and quality of total RNA was assessed with an Agilent 2100 Bioanalyzer (Agilent Technologies). Removal of rRNA was performed using RiboZero rRNA removal (bacteria) kit from Illumina, and 100-bp single-end reads libraries were prepared using a TruSeq Stranded Total RNA Kit (Illumina). Libraries were sequenced using a NextSeq550 sequencer (Illumina).

The raw reads were pre-processed with SeqTrimNext^66^ using the specific NGS technology configuration parameters. This pre-processing removes low-quality, ambiguous and low-complexity stretches, linkers, adapters, vector fragments, and contaminated sequences while keeping the longest informative parts of the reads. SeqTrimNext also discarded sequences below 25 bp. Subsequently, clean reads were aligned and annotated using the Pcl and Bamy reference genomes with Bowtie2^67^ in BAM files, which were then sorted and indexed using SAMtools v1.484^68^. Uniquely localized reads were used to calculate the read number value for each gene via Sam2counts (https://github.com/vsbuffalo/sam2counts). Differentially expressed genes (DEGs) were analyzed via DEgenes Hunter, which provides a combined p value calculated (based on Fisher’s method) using the nominal p values provided by from edgeR^69^ and DEseq2^70^. This combined p value was adjusted using the Benjamini-Hochberg (BH) procedure (false discovery rate approach) and used to rank all the obtained DEGs. For each gene, combined p value < 0.05 and log2-fold change >1 or <−1 were considered as the significance threshold (for Bamy vs GlpK^S166L^ the log2-fold change was fixed in >2 or <-2). The annotated DEGs were used to identify the Gene Ontology functional categories and KEGG pathways. Gephi software (https://gephi.org) was used to generate the DEG networks. RNAseq data was deposited to the GEO database as GSE161161.

### Isolation of spontaneous mutants and whole-genome sequencing

Bamy and Pcl strains were spotted 0.7 cm apart onto King B agar plates using 2 µl of cell suspension at an OD_600_ of 0.5. Plates were incubated at 30°C for 5−6 days. Next, small Bamy colonies and Pcl overgrowing subzones that were observed in the inhibition area were isolated and tested in co-culture assays (described below). Seven Bamy isolates and three Pcl isolates were used for whole-genome sequencing. Sequencing libraries were prepared using the PCRfree TrueSeq Kit from Illumina. 250-bp paired-end reads were sequenced using an Illumina MiSeq platform. Illumina raw reads, in fastq format, were trimmed using SeqtrimNext v2.1.3 to remove low-quality and low-complexity sequences, adapters, polyA tails, and several contaminant sequences. Useful reads were then mapped against their respective reference genomes (Bamy Genbank: NC_009725; Pcl Genbank: CP011110) to prepare data for subsequent variant calling using BWA configured with options –B 20 –A 30 –O 30 –E 3 to ensure high-quality alignments and accurate mapping scoring to prevent false variant calling. In the next step, a pileup file from each mapped sample was created using Samtools mpileup with options –BQ 26 –q 30 –d 10000000, as recommened by the Samtools official documentation for obtaining accurate results. Finally, mutations and InDels were independently called with VarScan2 using the pileup of bam files. Genome sequences were deposited at NCBI under Bioproject PRJNA680227.

### Compound purification

Pcl was grown on King B plates for 24 hours. Next, both bacteria and solid media were obtained, and a methanol (MeOH) (LC-MS grade, Fisher) extraction was performed with 15 minutes of sonication before centrifugation. The extracted solution was filtered and diluted to 10% MeOH with water (LC-MS grade, Fisher). Extracts were then pre-fractionated in solid phase C18 resin. After stepwise elution with 20, 40, 60, 80, and 100% MeOH (LC-MS grade Fisher), fractions were assayed to identify inhibitory compounds. Fractions with higher inhibitory activity were subjected to semi-preparative HPLC purification using a 10 x 150 mm C18 column (XBridge Waters). For the mobile phase, we used a flow rate of 5 mL/min (solvent A: H2O + 0.1% formic acid (FA), solvent B: acetonitrile (ACN) + 0.1 % FA). During the chromatographic separation, we applied a linear gradient from 0−1 min, 5% B, 1−10 min 5−50% B, 10−15 min 50−99% B, followed by a 5-minute washout phase at 99% B and a 5 minute re-equilibration phase at 5% B. 1-mL fractions were collected in deep well plates, and fractions of interest were then dried in a vacuum centrifuge (Centrivap, Labconco). Weight was recorded for all isolated compounds.

### Minimum inhibitory concentration (MIC) assays

MIC assays were performed in liquid LB medium using the two-fold serial dilution test according to the guidelines of the Clinical and Laboratory Standards Institute (2003). The highest concentrations of the compounds were: pyochelin (500 μg ml^−1^), HPR (1000 μg ml^−1^), kanamycin (1000 μg ml^−1^), gentamicin (1500 μg ml^−1^), and ampicillin (10000 μg ml^−1^). Experiments were carried out in triplicate and the MIC was determined as the lowest antibiotic concentration that inhibited growth by >90%.

### Matrix-assisted laser desorption ionization imaging mass spectrometry (MALDI-MSI)

To perform MALDI-MSI, a small section of King B agar containing the cultured microorganisms (both in single colonies and in interactions) were cut and transferred to a MALDI MSP 96 anchor plate. Deposition of matrix (1:1 mixture of 2,5-dihydroxybenzoic acid and α-cyano-4-hydroxycinnamic acid) over the agar was performed using a 53-µm molecular sieve. Next, plates were dried at 37 °C for 4 hours. Images were collected before and after matrix deposition. Samples were analyzed using a Bruker Microflex MALDI-TOF mass spectrometer (Bruker Daltonics, Billerica, MA, USA) in positive reflectron mode, with 300 µm–400 µm laser intervals in X and Y directions, and a mass range of 100–3200 Da. Data were analyzed using FlexImaging 3.0 software (Bruker Daltonics, Billerica, MA, USA). The acquired spectra were normalized by dividing all the spectra by the mean of all data points (TIC normalization method). The resulting mass spectrum was then filtered manually in 0.25% (3.0 Da) increments assigning colors to the selected ions associated with the metabolites of interest.

### Liquid chromatography-tandem mass spectrometry (LC-MS/MS)

Non-targeted LC-MS/MS analysis was performed on a Q-Exactive Quadrupole-Orbitrap mass spectrometer coupled to Vanquish ultra-high performance liquid chromatography (UHPLC) system (Thermo Fisher Scientific, Bremen, Germany) according to^71^. 5 µL of the samples were injected for UHPLC separation on a C18 core-shell column (Kinetex, 50 x 2 mm, 1.8-um particle size, 100 A-pore size, Phenomenex, Torrance, CA, USA). For the mobile phase, we used a flow rate of 0.5 mL/min (solvent A: H_2_O + 0.1% formic acid (FA), solvent B: acetonitrile (ACN) + 0.1% FA). During the chromatographic separation, we applied a linear gradient from 0–0.5 min, 5% B, 0.5–4 min 5–50% B, 4–5 min 50–99% B, followed by a 2-minute washout phase at 99% B and a 2-minute re-equilibration phase at 5% B. For positive mode MS/MS acquisition, the electrospray ionization (ESI) was set to a 35 L/min sheath gas flow, 10 L/min auxiliary gas flow, 2 L/min sweep gas flow, with a 400°C auxiliary gas temperature. The spray voltage was set to 3.5 kV with an inlet capillary of 250 °C. The S-lens voltage was set to 50 V. MS/MS product ion spectra were acquired in data-dependent acquisition (DDA) mode. MS1 survey scans (150–1500 m/z) and up to 5 MS/MS scans per DDA duty cycle were measured with a resolution (R) of 17,500. The C-trap fill time was set to a maximum of 100 ms or until the AGC target of 5E5 ions was reached. The quadrupole precursor selection width was set to 1 m/z. Normalized collision energy was applied stepwise at 20, 30, and 40% with z = 1 as the default charge state. MS/MS scans were triggered with apex mode within 2–15 seconds from their first occurrence in a survey scan. Dynamic precursor exclusion was set to 5 seconds. Precursor ions with unassigned charge states and isotope peaks were excluded from MS/MS acquisition.

### Data analysis and MS/MS network analysis

After LC-MS/MS acquisition, raw spectra were converted to .mzXML files using MSconvert (ProteoWizard). MS1 and MS/MS feature extraction was performed with Mzmine2.30^72^. For MS1 spectra, an intensity threshold of 1E5 was used, and for MS/MS spectra, an intensity threshold of 1E3 was used. For MS1 chromatogram building, a 10-ppm mass accuracy and a minimum peak intensity of 5E5 was set. Extracted ion chromatograms (XICs) were deconvolved using the baseline cut-off algorithm at an intensity of 1E5. After chromatographic deconvolution, XICs were matched to MS/MS spectra within 0.02 m/z and 0.2-minute retention time windows. Isotope peaks were grouped and features from different samples were aligned with 10 ppm mass tolerance and 0.1-minute retention time tolerance. MS1 features without MS2 features assigned were filtered out the resulting matrix as well as features that did not contain isotope peaks and that did not occur in at least three samples. After filtering, gaps in the feature matrix were filled with relaxed retention time tolerance at 0.2 minute but also 10 ppm mass tolerance. Finally, the feature table was exported as a .csv file, and corresponding MS/MS spectra exported as .mgf files. Contaminate features observed in blank samples were filtered, and only those with a relative abundance ratio blank to average lower than 50% were considered for further analysis.

For feature-based molecular networking and spectrum library matching, the .mgf file was uploaded to GNPS (gnps.ucsd.edu)^27^. For molecular networking, the minimum cosine score was set to 0.7. The precursor ion mass tolerance was set to 0.01 Da, and the fragment ion mass tolerance was set to 0.01 Da. Minimum matched fragment peaks were set to 4, minimum cluster size was set to 1 (MS Cluster off), and library search minimum matched fragment peaks were set to 4. When analog searches were performed, the cosine score threshold was 0.7 and the maximum analog search mass difference was 100 *m/z*. Molecular networks were visualized with Cytoscape version 3.484. Feature-based molecular networking analysis can be accessed through the following link: https://gnps.ucsd.edu/ProteoSAFe/status.jsp?task=574c0acbea58405e80e1c0dc526bc903.

To enhance the chemical structural information in the molecular network, information from *in silico* structure annotations from GNPS Library Search, Network Annotation Propagation were incorporated into the network using the GNPS MolNetEnhancer workflow (https://ccms-ucsd.github.io/GNPSDocumentation/molnetenhancer/) on the the GNPS website (http://gnps.ucsd.edu). Chemical class annotations were performed using the ClassyFire chemical ontology^27–30^. MolNetEnhancer analysis can be accessed through the following link: https://gnps.ucsd.edu/ProteoSAFe/status.jsp?task=15e9aa6189e04c859b685eb0eb0f6089

### MS/MS data availability

All LC-MS/MS data were deposited to the Mass Spectrometry Interactive Virtual Environment (MassIVE) at https://massive.ucsd.edu/ with the identifier MSV000085326: https://gnps.ucsd.edu/ProteoSAFe/result.jsp?task=d7df96995b1a443ba826a7c5b96888f7=view=advanced_view

### Docking and *in silico* analysis of proteins

I-Tasser workspace (https://zhanglab.ccmb.med.umich.edu/I-TASSER/) was used for automated protein tertiary structure homology modeling of FusA^73^. To identify potential binding sites of bacillaene (PubChem ID: 25144999) and fusidic acid (PubChem ID: 3000226) to the Pcl *<u>fusA</u>*-encoded protein, automated molecular docking and thermodynamic analysis were performed using the web-based SwissDock program (www.swissdock.ch/docking)^74^. SwissDock predicts possible molecular interactions between a target protein and a small molecule based on the EADock DSS docking algorithm ^74^. Docking was performed using the “Accurate” parameter with otherwise default parameters, with no region of interest defined (blind docking). Binding energies were estimated via CHARMM (Chemistry at HARvard Macromolecular Mechanics), a molecular simulation program implemented within SwissDock software. The most favorable energies are evaluated via FACTS (Fast Analytical Continuum Treatment of Solvation). Finally, the resulting energies were scored and ranked based on full fitness (kcal mol^−1^), and the spontaneous binding was reported as the estimated Gibbs free energy ΔG (kcal mol^−1^). Negative ΔG values support the assertion that the binding process is highly spontaneous. Modeling and docking results were visualized using UCSF Chimera v1.8 software.

### Membrane staining with DilC12

To detect regions of increased membrane fluidity, staining with DiIC12 (Thermo Fisher) staining was performed. Bamy strains were grown overnight on King B plates and then resuspended in 1 ml dH_2_O followed by addition of 1 μg ml^−1^ DiIC12. 0.5% Benzyl-alcohol was added to positive control samples. Cells were incubated at 28 °C for 3 hours and then washed four times and resuspended in dH_2_O. Images were obtained by visualizing samples using a Leica SP5 confocal microscope with a 63x NA 1.3 Plan APO oil immersion objective and acquisitions with excitation at 651 nm. Image processing was performed using FIJI/ImageJ software. For each experiment, laser settings, scan speed, PMT detector gain, and pinhole aperture were kept constant for all acquired image stacks.

### Membrane potential and respiration assays

Membrane potential was evaluated using the image-iT TMRM reagent (Invitrogen) following the manufacturer’s instructions. Colonies grown at 28 °C on King B solid medium were isolated at 24 hours and resuspended in 1X phosphate-buffered saline. As a control, samples were treated with 20 µM carbonyl cyanide m-chlorophenyl hydrazine (CCCP), a known protonophore and uncoupler of bacterial oxidative phosphorylation, prior to staining. TMRM was added to the bacterial suspensions to a final concentration of 100 nM, and the mixtures were incubated at 37 °C for 30 minutes. After incubation, the cells were immediately visualized by confocal laser scanning microscopy (CLSM) with excitation at 561 nm and emission detection between 576 and 683 nm.

Respiratory activity was measured using 5 mM CTC staining following the manufacturer’s instructions. Colonies were grown as described above, and samples were incubated with CTC at 37 °C for 30 minutes. After incubation, the cells were immediately visualized by CLSM with excitation at 561 nm and emission detection between 576 and 683 nm.

### Statistical analysis

Results are expressed as mean ± SEM. Statistical significance was assessed using ANOVA or Student’s t-test. All analyses were performed using GraphPad Prism® version 6. P-values <0.05 were considered significant.

## Acknowledgments

We thank Saray Morales Rojas for technical support, the Ultrasequencing Unit of the SCBI-UMA for RNA sequencing and the Bioinformatic unit of GENYO (Granada, Spain) for the analytical treatment of the data. We are grateful to Francisco M Cazorla for kindly providing the wild type strain PCL1606, suggestions and comments. This work was supported by grants from ERC Starting Grant (BacBio 637971) and Plan Nacional de I+D+i of Ministerio de Economía y Competitividad and Ministerio de Ciencia e Innovación (AGL2016-78662-R and PID2019-107724GB-I00). C.M.S is funded by the program Juan de la Cierva Incorporación (IJC2018-036923-I). A.I.P.L is funded by the program FPU (FPU19/00289), JAE-Intro 2019 (JAEINT 19 00269) and the program Plan Propio de Investigación y Transferencia from Universidad de Málaga. D.P. was supported by the German Research Foundation (DFG) with Grant PE 2600/1. A.M.C.R. and P.C.D. were supported by the National Science Foundation grant IOS-1656481 and National Institutes of Health (NIH) grant number 1DP2GM137413-01. P.C.D. was supported by the Gordon and Betty Moore Foundation through Grant GBMF7622, and the NIH (P41 GM103484, R03 CA211211, R01 GM107550).

## Author contributions

D.R. conceived the study; D.R. and C.M.S. designed the experiments; C.M.S. performed the main experimental work; C.M.S. and J.P. performed and designed the confocal microscopy work and data analysis; D.V.C. performed the docking analyses; L.D.M. and C.M.S. performed the RNAseq and genomic data analyses; C.M.S., S.S.T., and A.I.P.L. constructed strains; C.M.S., D.P., P.C.D., and A.M.C.R. performed MALDI-MSI and LC-MS/MS analysis; D.R. and C.M.S. wrote the manuscript; and A.V., D.P., A.M.C.R., and P.C.D. contributed critically to writing and editing the final version of the manuscript.

## Competing interests

The authors declare no competing interests.

## Notes

### Competing Interest Statement

The authors have declared no competing interest.

