## Supplementary Data for "Chemical interplay and complementary adaptative strategies toggle bacterial antagonism and co-existence"

### Supplementary Figures

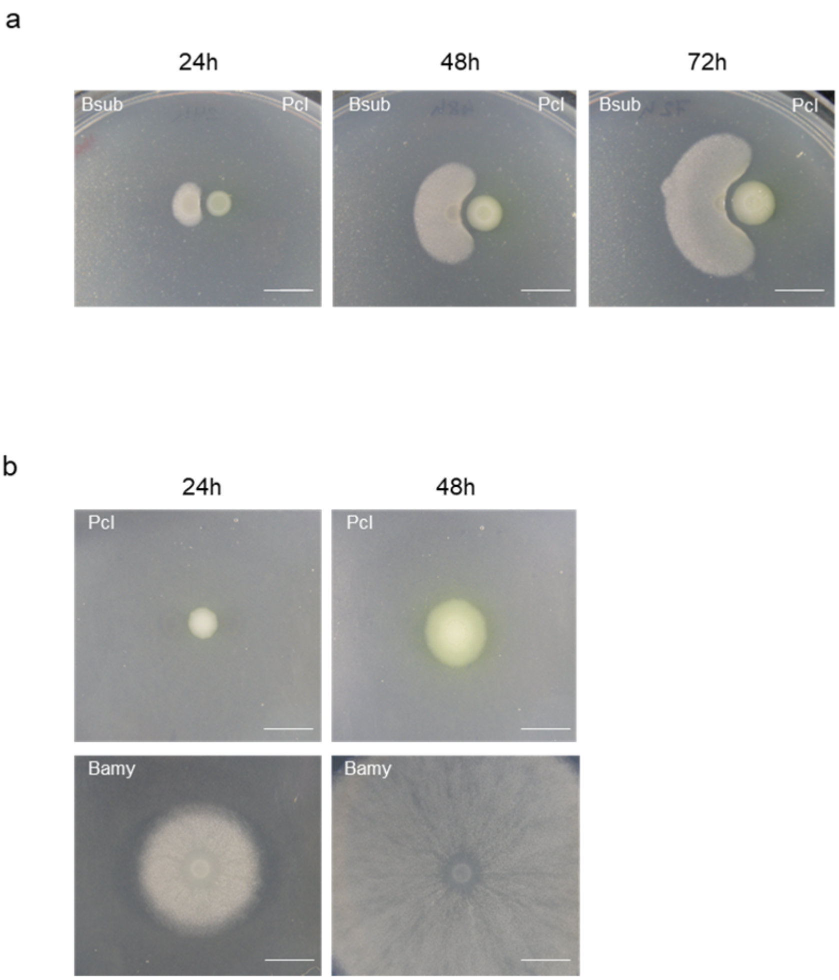


Suppl. Figure 1. Pcl inhibits *B. subtilis* 3610 on King B medium. a) Pairwise interaction time-lapse between *B. subtilis* 3610 (left) and Pcl (right) in King B medium at 24, 48, and 72 h. Scale = 10 mm. b) Single colony growth of Pcl (top) and Bamy (bottom) on King B medium at 24 and 48 h.

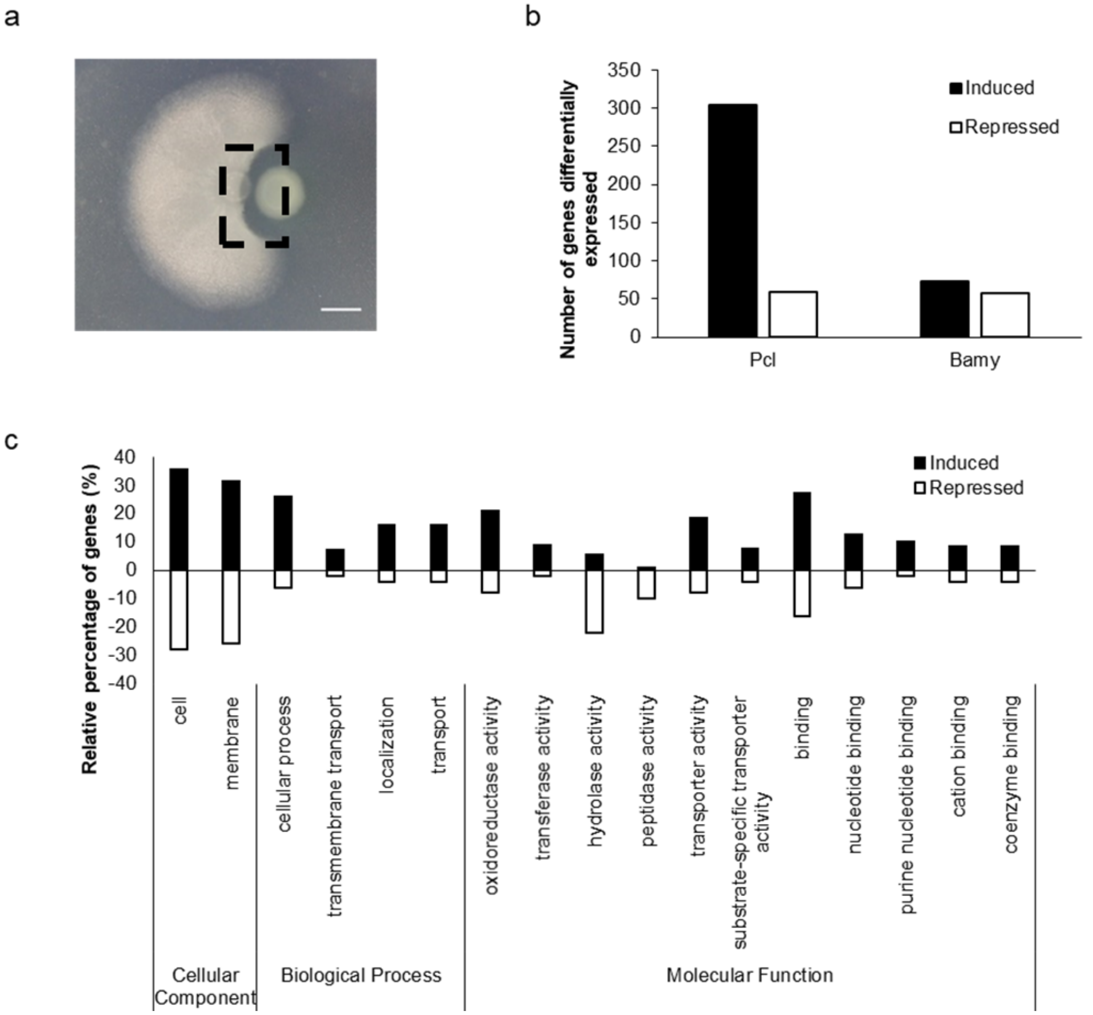


Suppl. Figure 2. Transcriptomic changes during the interaction of Pcl with Bamy at 24 h. a) Representation of the interaction area analyzed by RNAseq. Scale = 5 mm. b) Number of differentially expressed genes in Pcl and Bamy strains. Black bars indicate induced genes and empty bars indicate repressed genes. c) GO terms of differentially expressed genes in Pcl. Black bars indicate induced GO terms and empty bars indicate repressed GO terms.

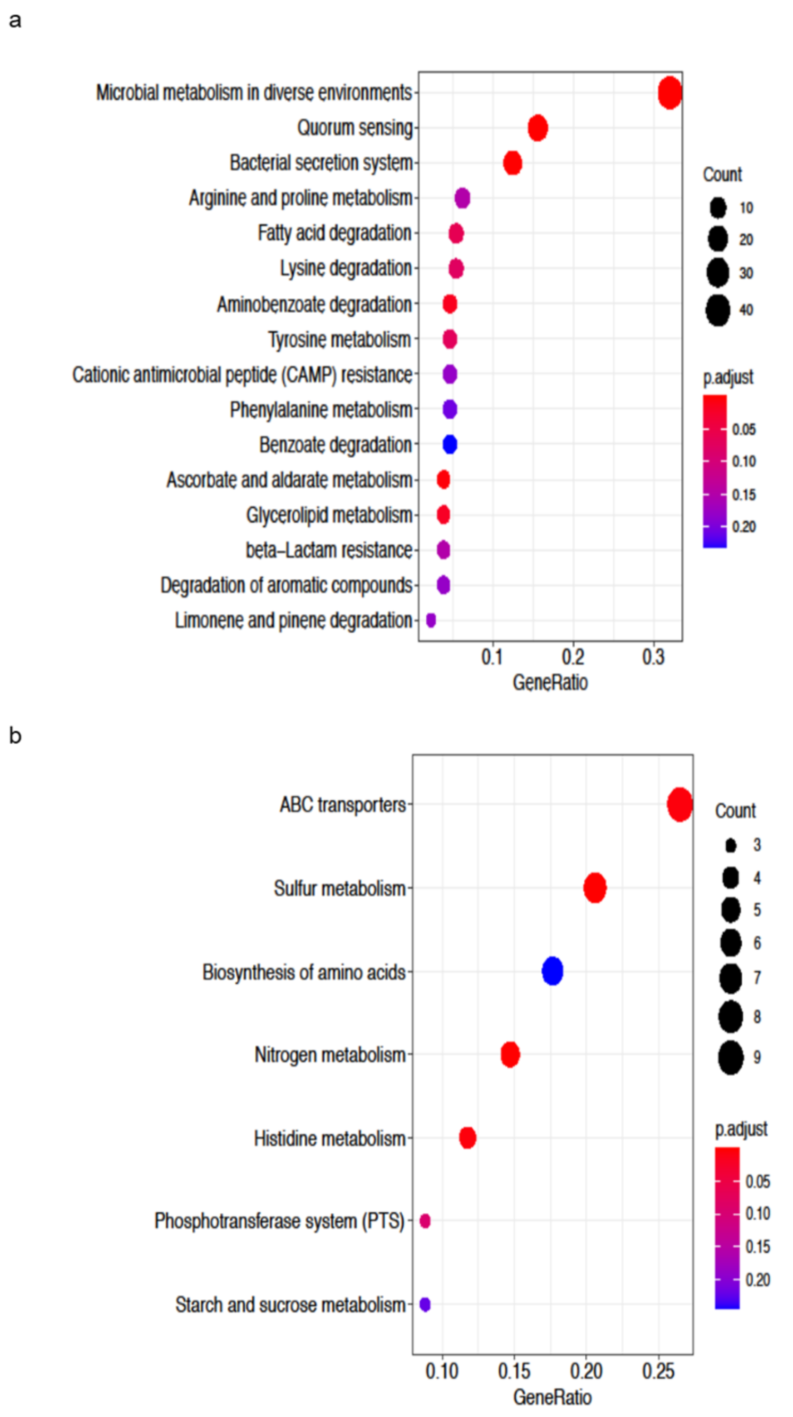


Suppl. Figure 3. KEGG pathways differentially expressed during the interaction in a) Pcl and b) Bamy compared with control samples.

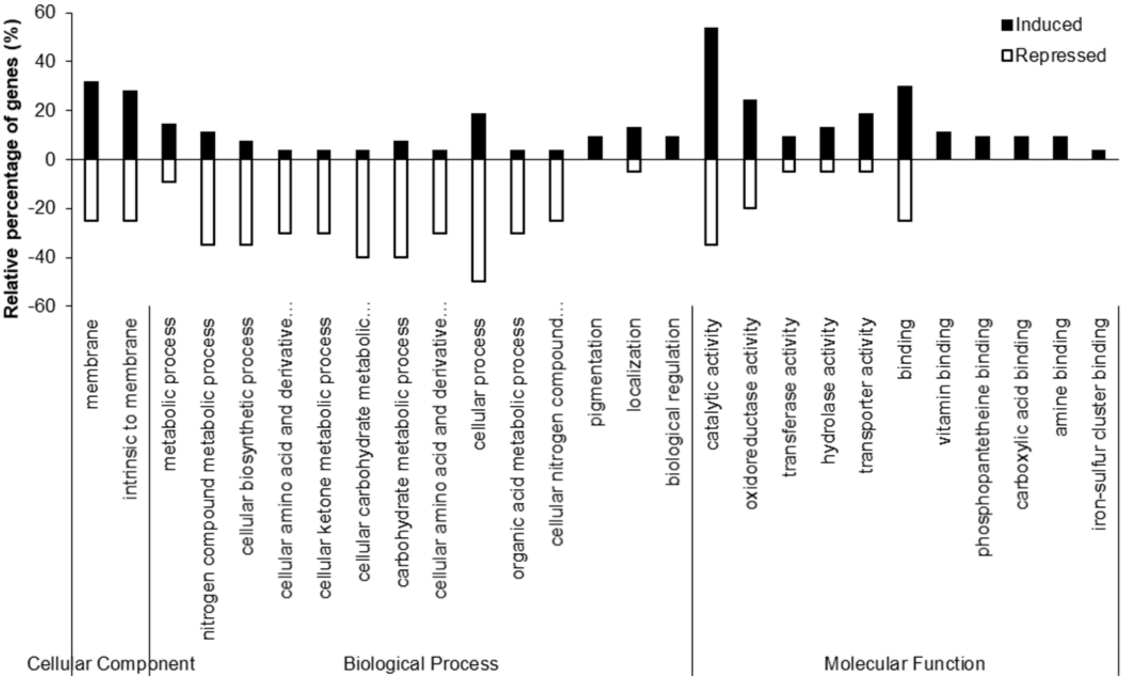


Suppl. Figure 4. Transcriptomic changes during the interaction of Pcl with Bamy at 24 h. GO terms of differentially expressed genes in Bamy. Black bars indicate induced GO terms and empty bars indicate repressed GO terms.

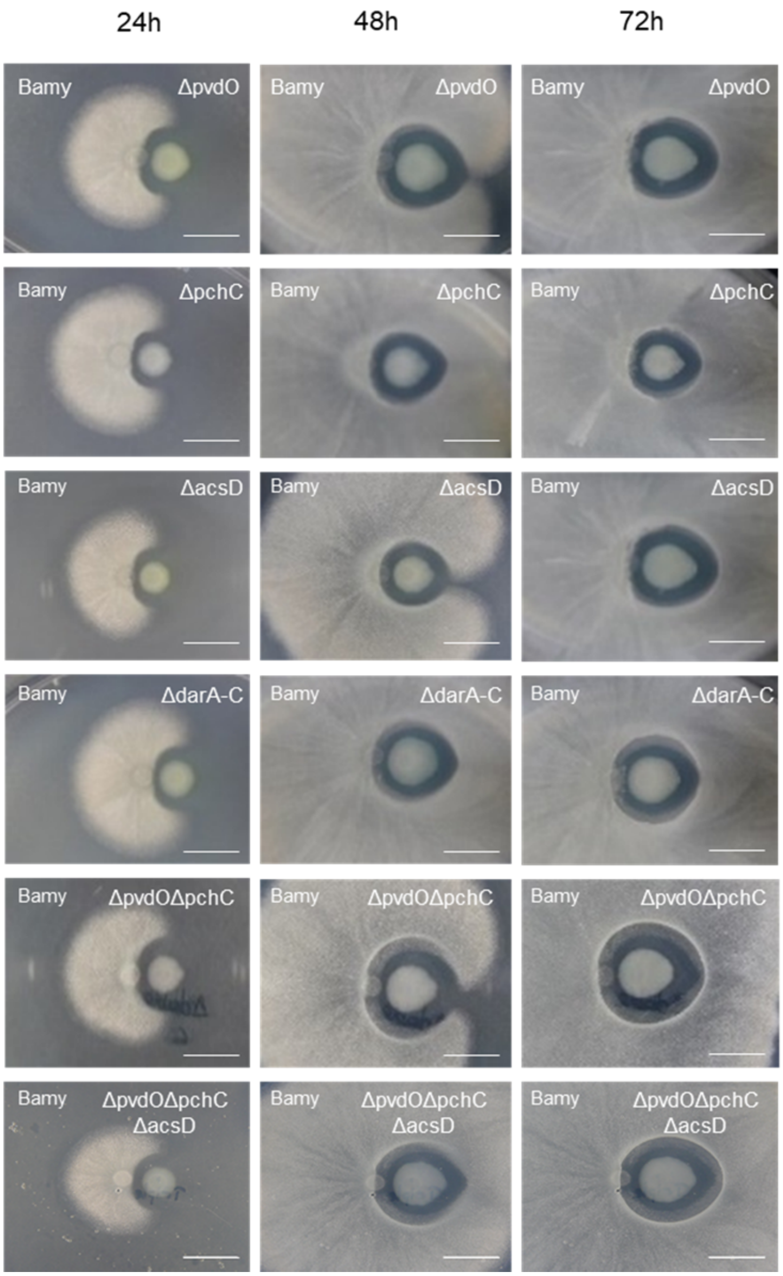


Suppl. Figure 5. Time-course pairwise interactions between Bamy and Pcl mutants involving secondary metabolites on King B medium at 24, 48, and 72 h. Scale = 10 mm. ΔpvdO = pyoverdine mutant; ΔpchC = pyochelin mutant; ΔacsD = achromobactin mutant; ΔdarA-C = HPR mutant; ΔpvdOΔpchC = double mutant in pyoverdine and pyochelin; ΔpvdOΔpchCΔacsD = triple mutant in pyoverdine, pyochelin, and achromobactin.

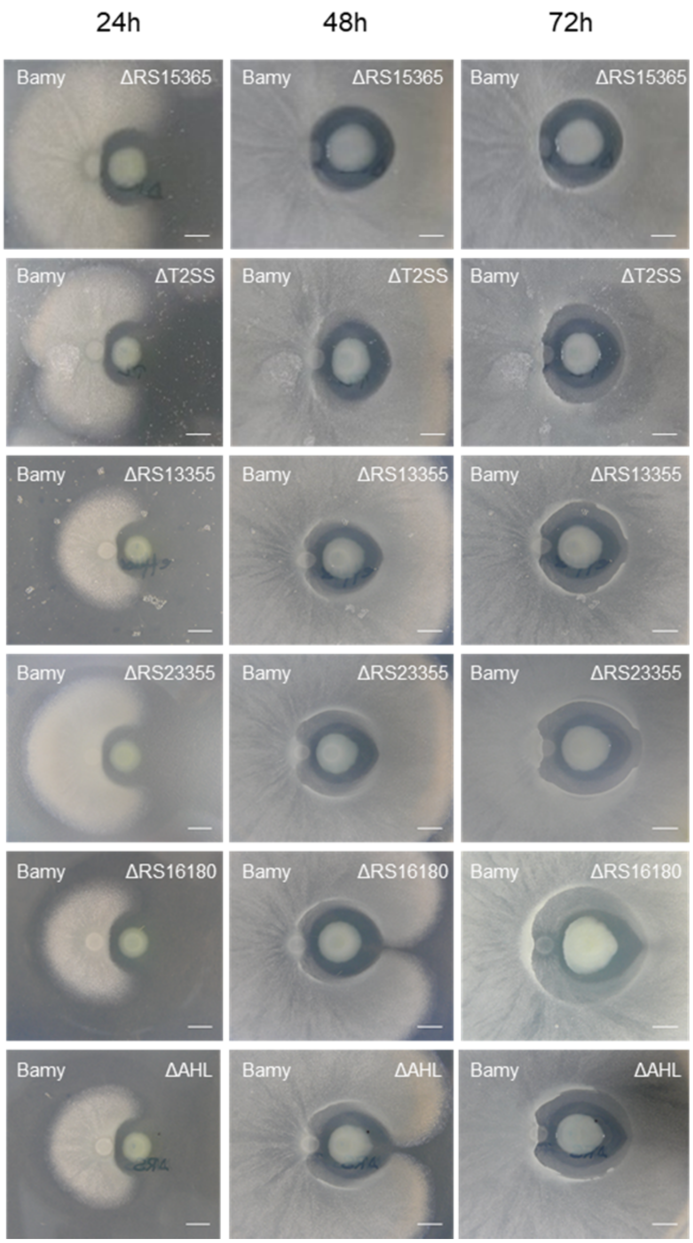


Suppl. Figure 6. Time-course pairwise interactions between Bamy and Pcl mutants involving secondary metabolites, T2SS, and efflux pumps on King B medium at 24, 48, and 72 h. Scale = 5 mm. ΔAHL = acyl-homoserine lactone mutant; ΔT2SS = type II secretion system mutant.

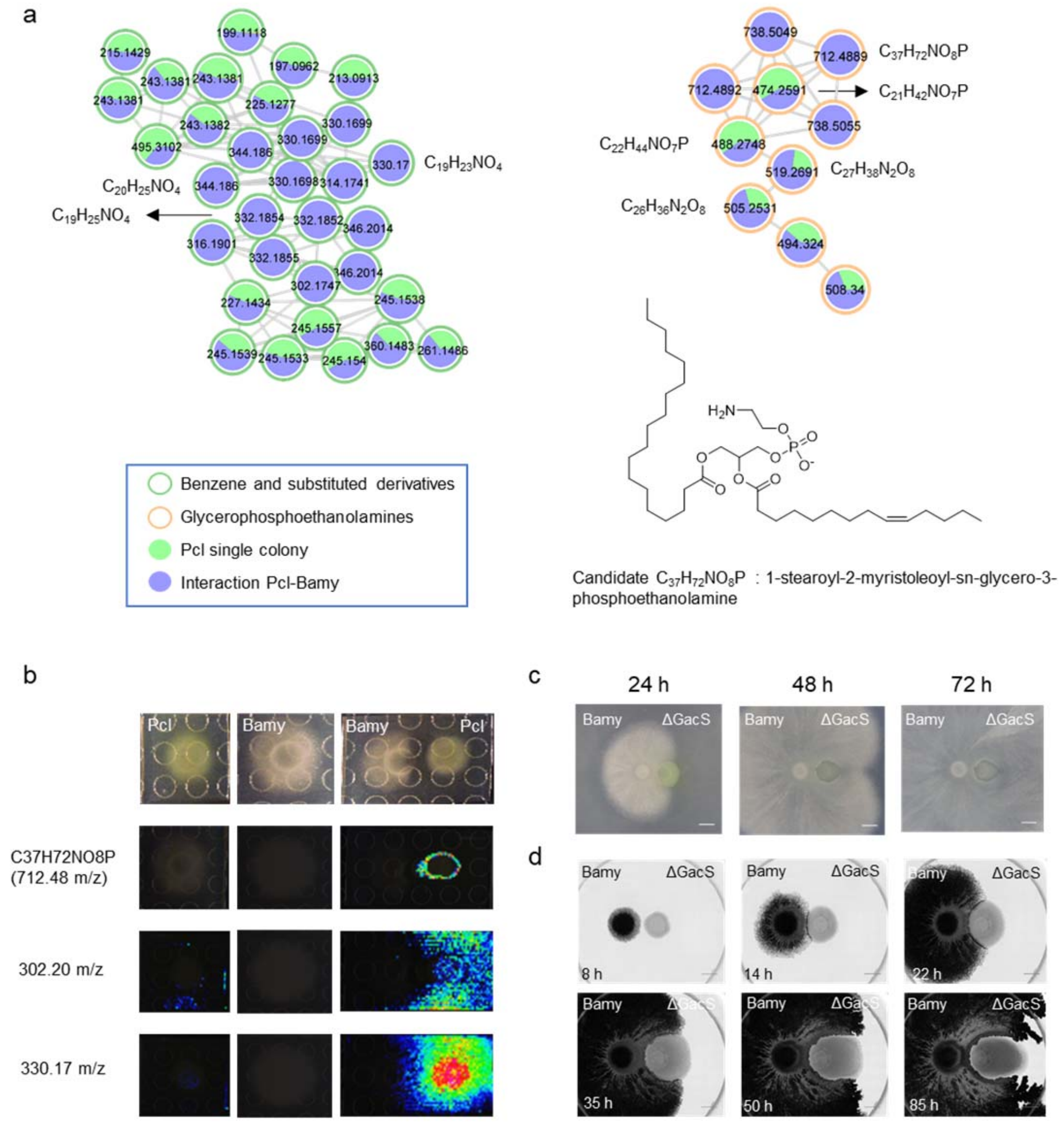


Suppl. Figure 7. Candidate Pcl metabolites responsible for Bamy inhibition. a) Molecular families of Pcl secondary metabolites mostly detected during the interaction with Bamy. Chemical structure of annotated features based on SIRIUS analyses, as representative of these molecular families. Border color indicates ClassyFire classification. The sizes of the compounds are directly related to their abundance in the metabolome. Squares indicate a library hit level 2 in GNPS, while circles indicate unknown compounds based on GNPS. b) MALDI-TOF-MSI heatmaps showing the spatial distribution of representative metabolites indicated in panel a with m/z 712.48, 302.20 and 330.17. c) Pairwise interaction time lapse between Bamy (left) and ΔGacS (right) on King B media at 24, 48, and 72 h. Scale = 5 mm. d) Time-lapse microscopy of the pairwise interaction between Bamy (left) and ΔGacS (right) during 85 h. Scale= 2 mm.

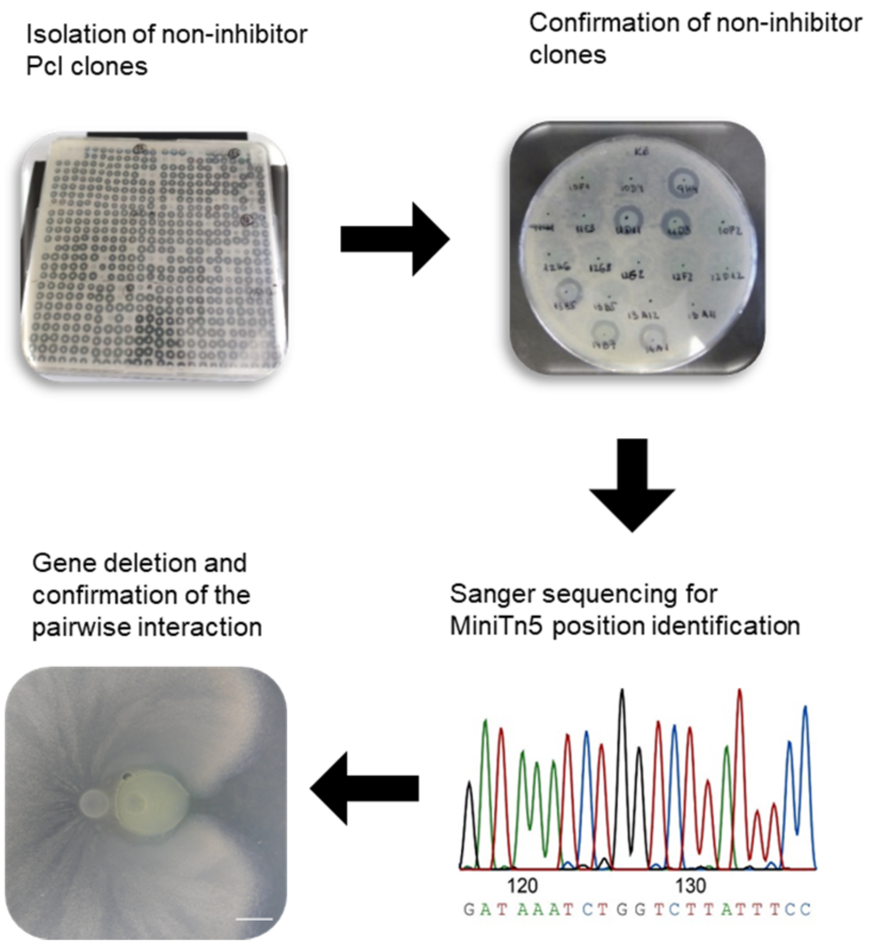


Suppl. Figure 8. Schematic representation of the workflow from miniTn5 transposon mutagenesis to isolation of non-inhibitor clones to sequence confirmation and pairwise interaction analysis.

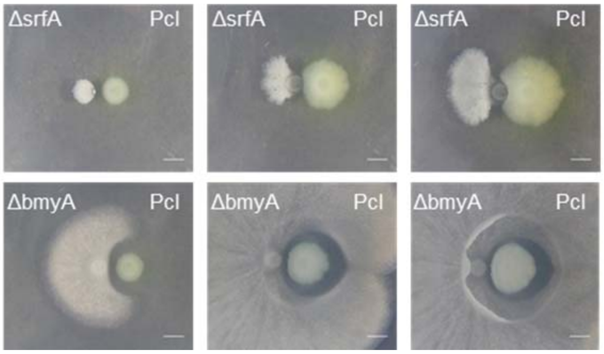


Suppl. Figure 9. Pairwise interactions between Pcl and Bamy mutants related to the secondary metabolites surfactin (ΔsrfA) (top) and bacillomycin (ΔbmyA) (bottom). Scale = 5 mm.

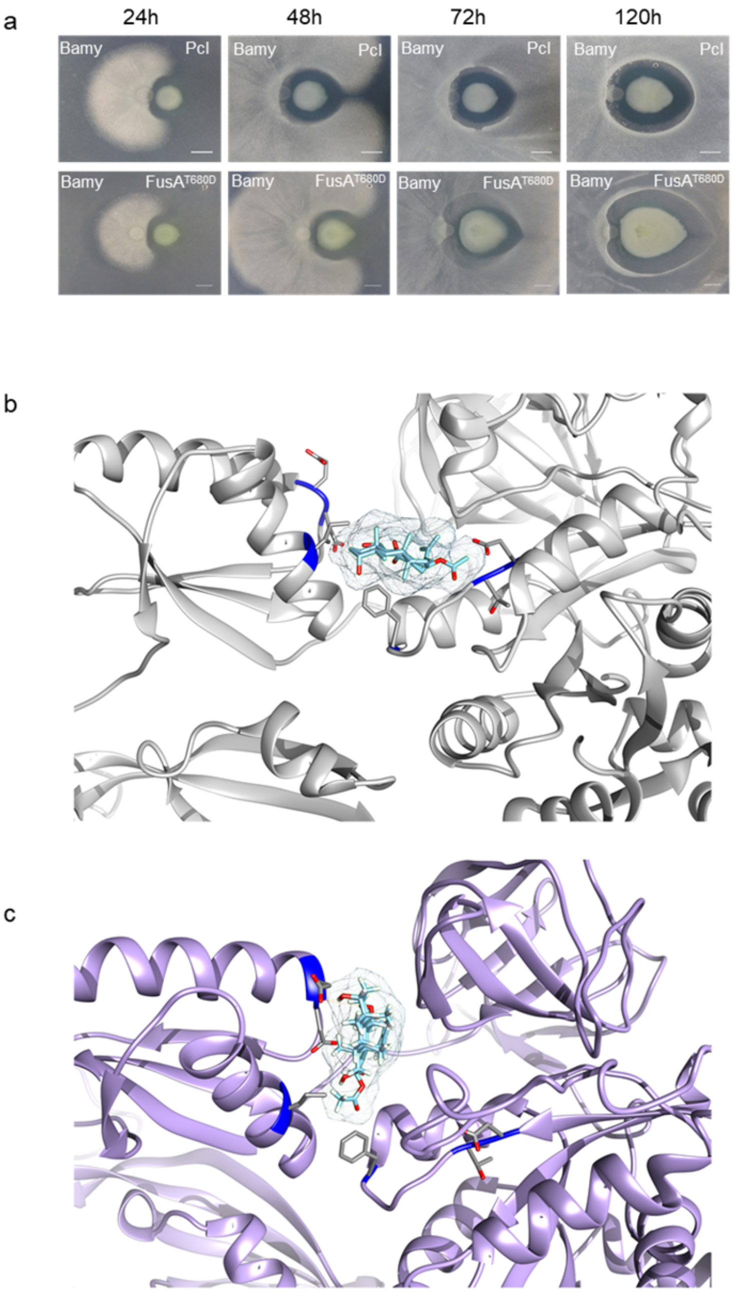


Suppl. Figure 10. a) Pairwise interactions between Bamy and Pcl WT (top) or FusA$^{T680D}$ (bottom) on King B medium at 24, 48, 72 and 120 h. Scale = 5 mm. b and c) Molecular docking between fusidic acid and (b) FusA or (c) FusA$^{T680D}$ showing differences in the binding pocket.

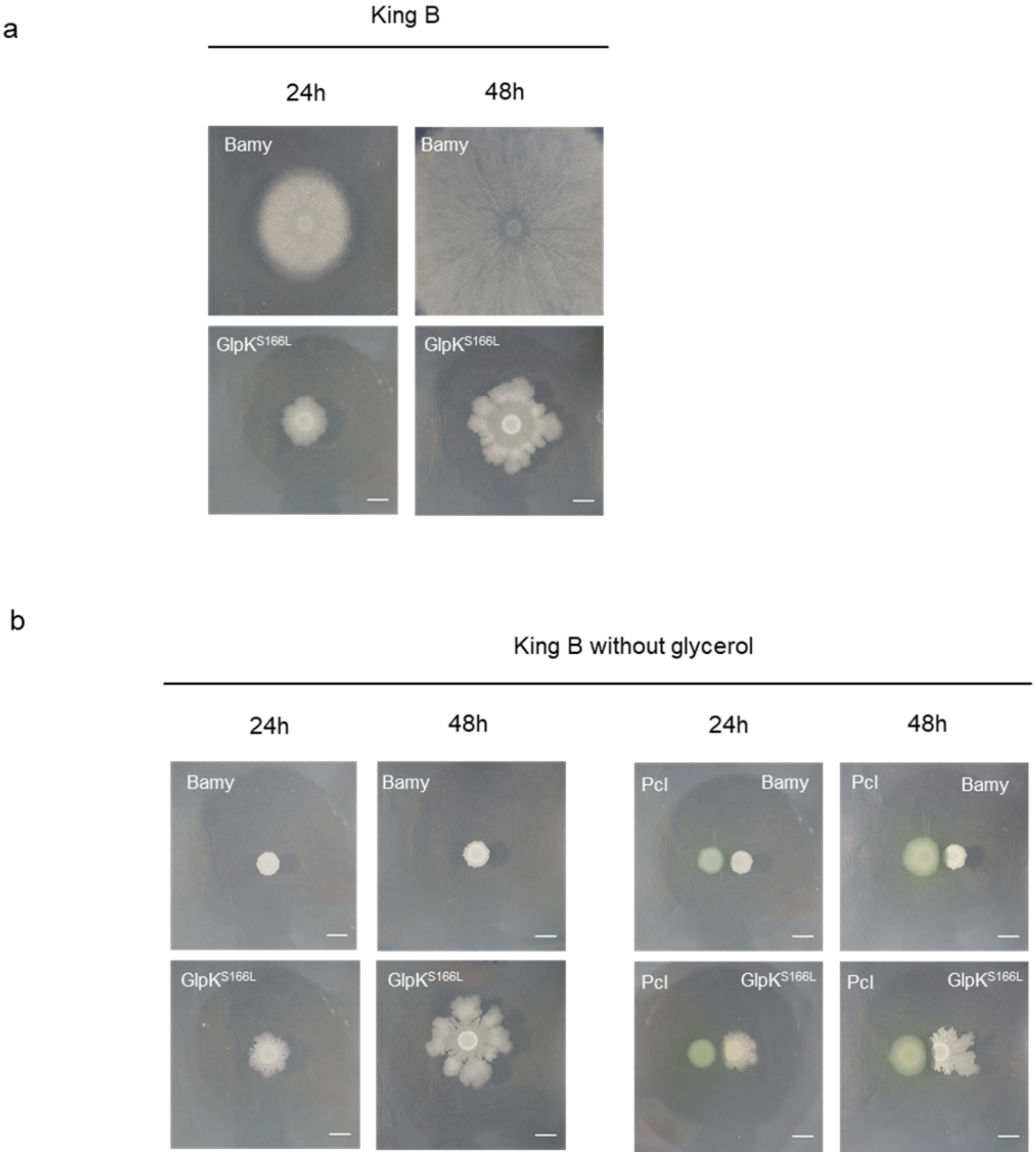


Suppl. Figure 11. Growth differences in Bamy strains on King B medium with and without glycerol. a) Single Bamy and GlpK$^{S166L}$ colonies growing on King B medium at 24 and 48 h. b) Single Bamy and GlpK$^{S166L}$ colonies and their interactions with Pcl growing on King B medium without glycerol at 24 and 48 h.

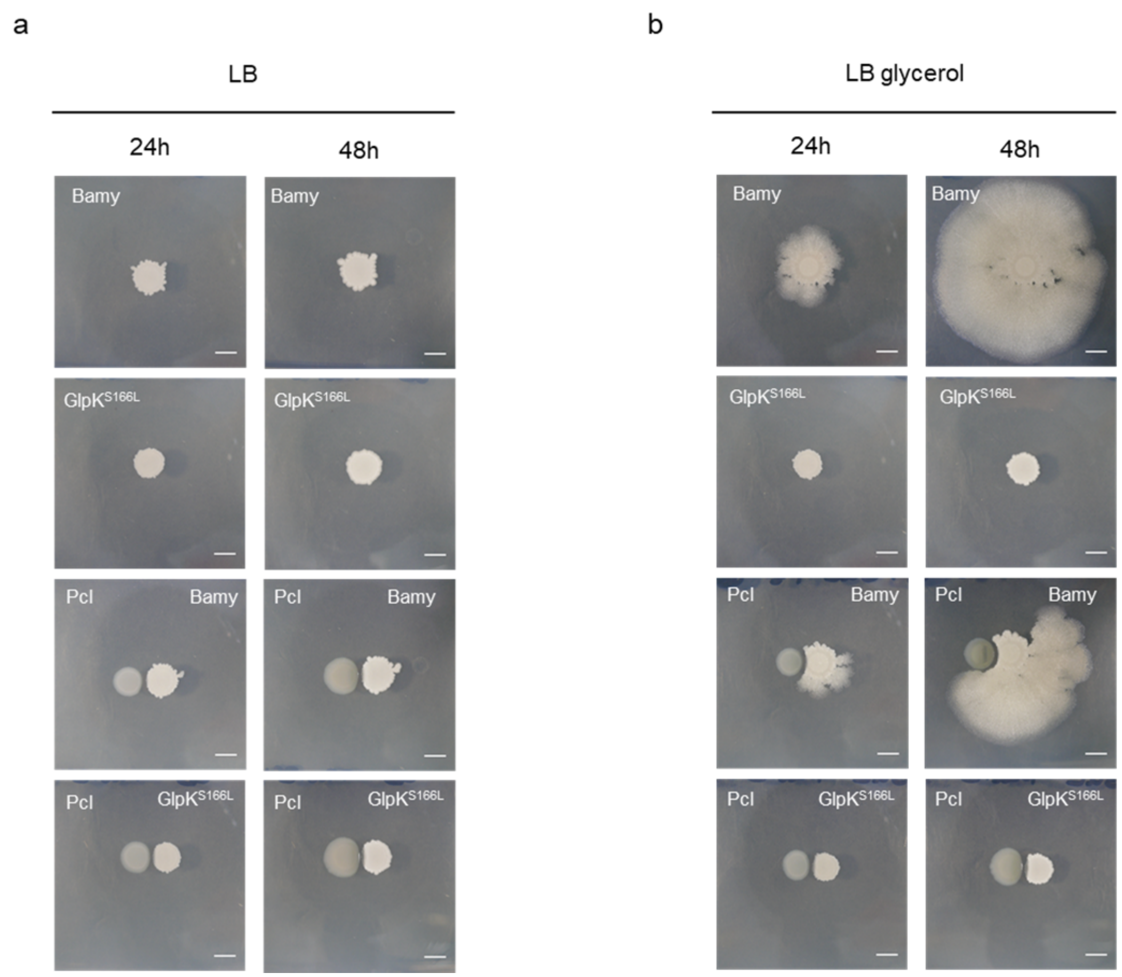


Suppl. Figure 12. Growth differences in Bamy strains on LB medium with and without glycerol. a) Single Bamy and GlpK$^{S166L}$ colonies and their interactions with Pcl growing on LB B medium at 24 and 48 h. b) Single Bamy and GlpK$^{S166L}$ colonies and their interactions with Pcl growing on LB medium supplemented with glycerol at 24 and 48 h.

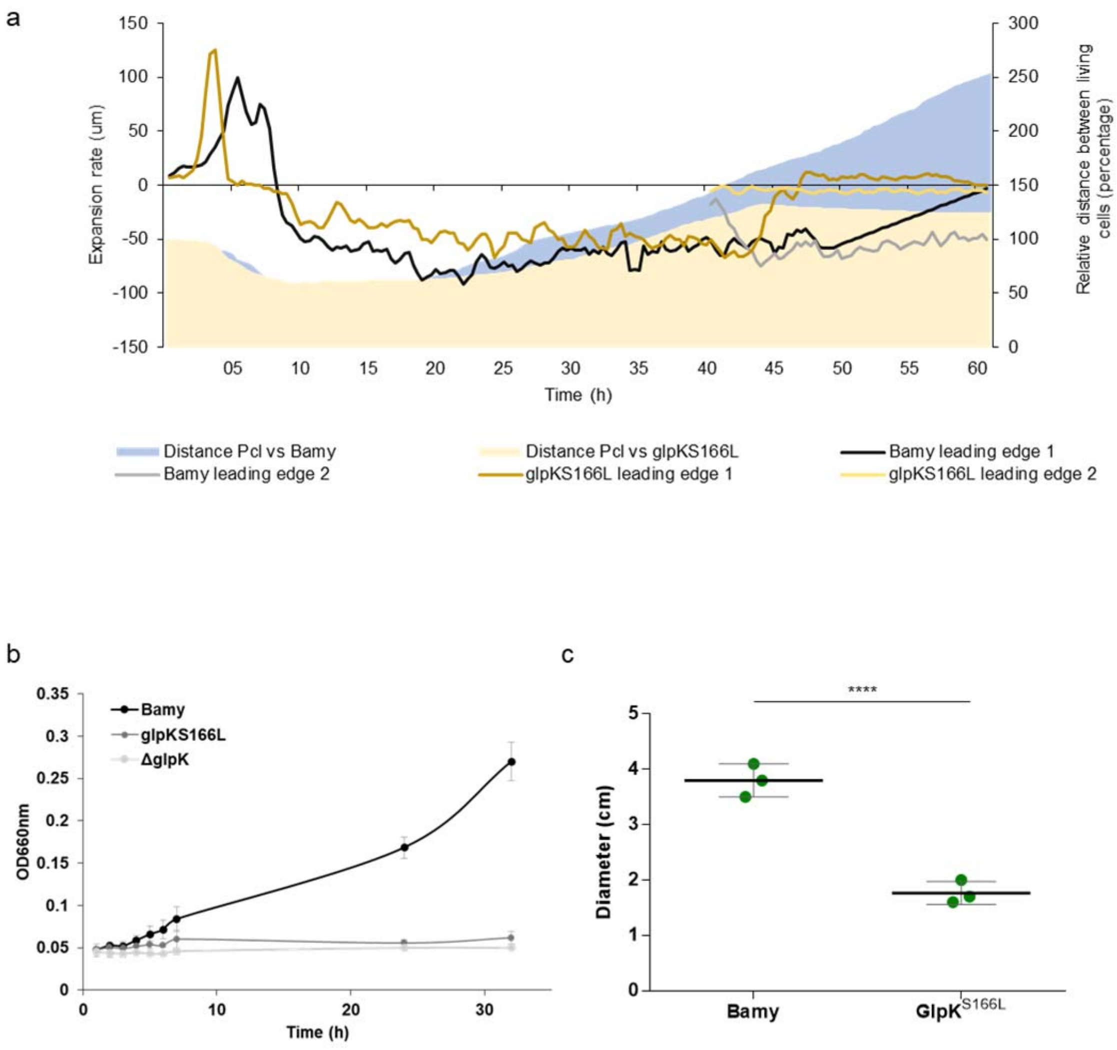


Suppl. Figure 13. Phenotypic changes in a GlpK$^{S166L}$ mutant. a) Expansion rates and distance of the GlpK$^{S166L}$ and Bamy leading edges during the interaction with Pcl. Brown and yellow lines represent GlpK$^{S166L}$ leading edges 1 and 2. Black and grey lines represent Bamy leading edges 1 and 2. Blue area represents the distance between the Bamy and Pcl populations during the entire interaction while the light orange area represents the distance between the GlpK$^{S166L}$ and Pcl populations. b) Growth curves of Bamy (black dots), GlpK$^{S166L}$ (dark grey dots) and ΔglpK (grey dots) in M9 supplemented with 5 mM glycerol. c) Swarming motility reduction in a GlpK$^{S166L}$ mutant. The plot represents the diameter of the colony. ****P-value < 0.0001.

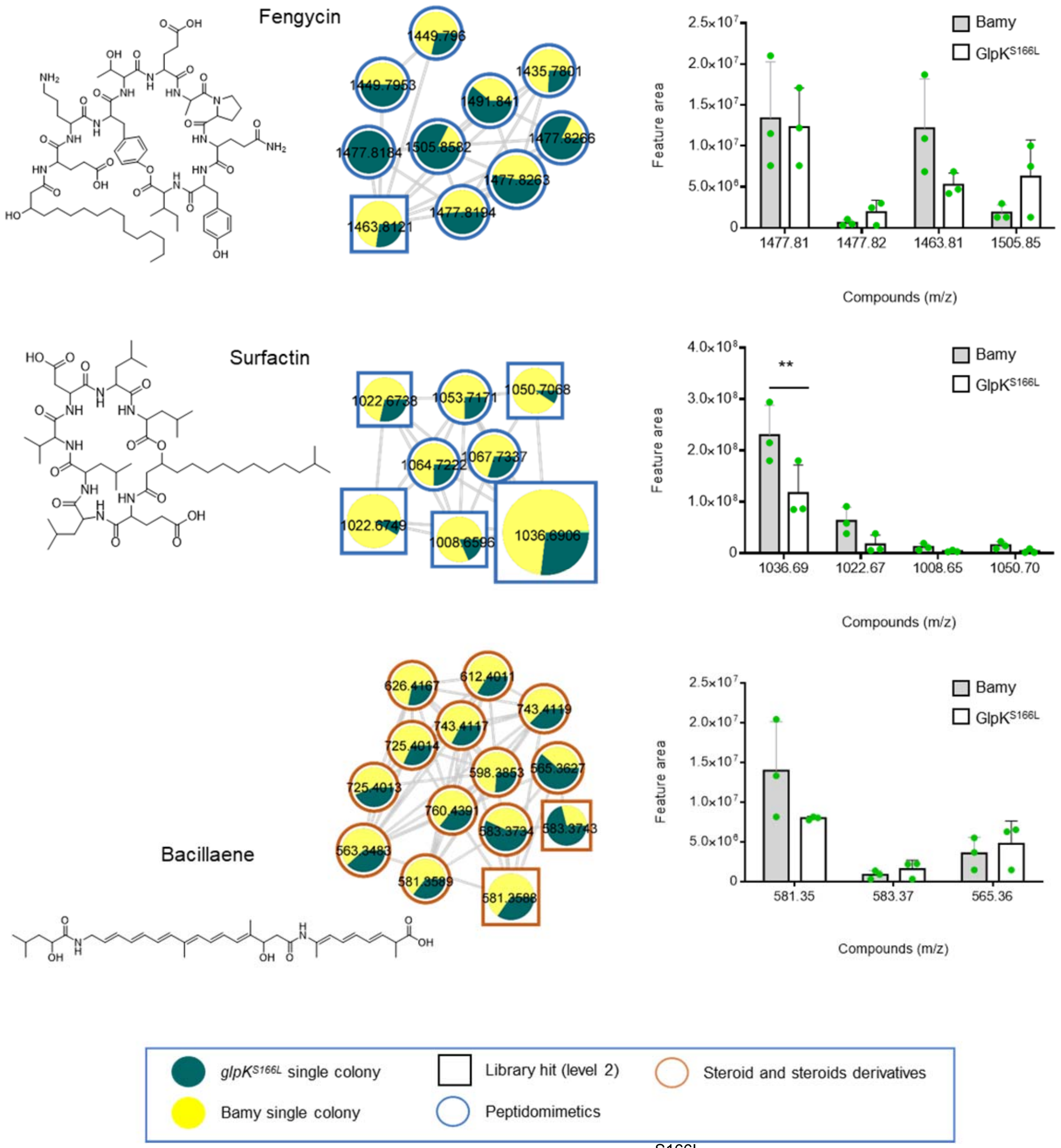


Suppl. Figure 14. Metabolomic changes found in GlpK$^{S166L}$ in comparison with Bamy. Molecular families of the secondary metabolites fengycin, surfactin, and bacillaene. The chemical structures of annotated features are based on spectral matches to GNPS libraries as representative of these molecular families. Border color indicates ClassyFire classification. Yellow color represents relative abundance in Bamy, while dark green color represents relative abundance in GlpK$^{S166L}$. Bar plots show the quantification of the relative abundance of the selected molecules in the interactions in Bamy and GlpK$^{S166L}$ strains. Grey bars represent Bamy feature area of metabolites, while empty bars represent GlpK$^{S166L}$ feature area of metabolites.

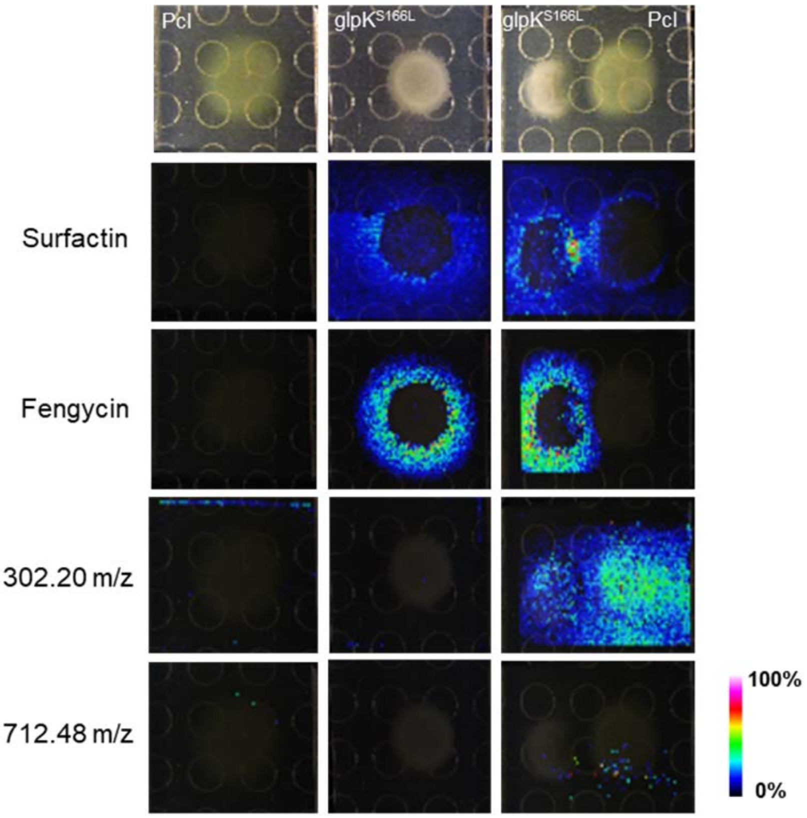


Suppl. Figure 15. Changes in spatial distribution of metabolites found in GlpK$^{S166L}$ and Bamy strains. MALDI-TOF-MSI heatmaps showing the spatial distribution of the secondary metabolites surfactin and fengycin produced by GlpK$^{S166L}$ growing alone and during the interaction with Pcl and the candidate inhibitory molecules (i.e., those with m/z 302.20 and 712.48) produced by Pcl.

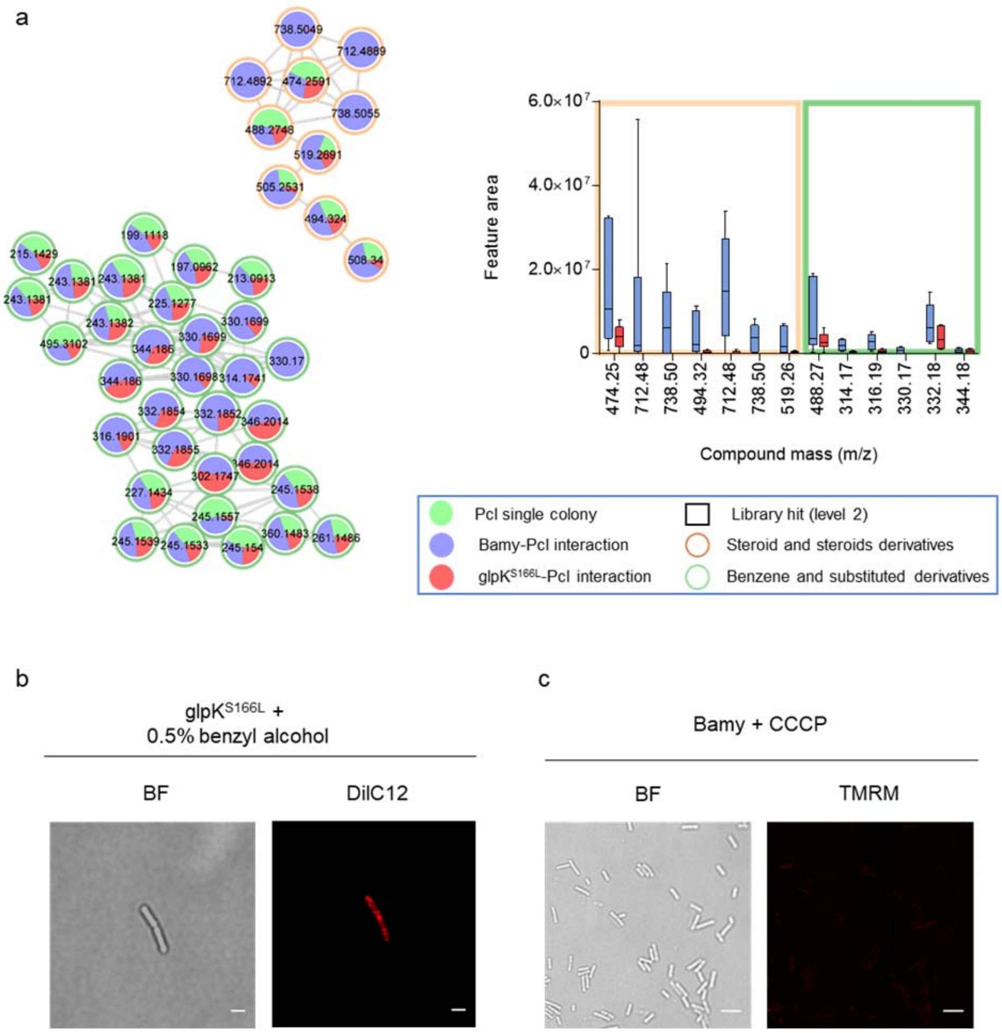


Suppl. Figure 16. Metabolomic changes found in GlpK$^{S166L}$ during the interaction with Pcl in comparison with the interaction of Bamy with Pcl. a) Changes in the abundance of candidate molecular families of secondary metabolites produced by Pcl during the interaction with Bamy and GlpK$^{S166L}$. The chemical structures of annotated features are based on spectral matches to GNPS libraries, as representative of these molecular families. Border color indicates ClassyFire classification. Right panel represents the quantification of the relative abundance of the selected molecules during Pcl-Bamy and Pcl-GlpK$^{S166L}$ interactions. b) Membrane staining of GlpK$^{S166L}$ with (left panel) DiIC12 dye using 0.5% benzyl alcohol as positive control (scale = 2 μm). c) TMRM positive control experiment of Bamy supplemented with 20 μM CCCP (scale 5 μm).

**Supplementary Tables**

Suppl. Table 1. Pcl genes induced and repressed during the Pcl-Bamy interaction.

Excel file.

Suppl. Table 2. Bamy genes induced and repressed during the Pcl-Bamy interaction.

Excel file.

Suppl. Table 3. Metabolites purified from a Pcl culture with MIC activity against Bamy.

| Mass (m/z) | Compound | MIC Bamy (µg/ml) |
|---|---|---|
| 237.18 | 2-Hexyl-5-propyl-rescorcinol (HPR) | 40 |
| 239.2 | unknown | 62.5 |
| 339.08 | Pyochelin methyl ester | 250 |
| 444.22 | unknown | 250 |

Suppl. Table 4. MIC of PcI and *fusA* mutants to aminoglycoside antibiotics kanamycin (Km) and gentamicin (Gm).

| | MIC (µg/ml) | |
|---|---|---|
| | **Km** | **Gm** |
| PcI | 4.5 | 2 |
| *fusA*$^{T680D}$ | 9 | 20 |
| *fusA*$^{K366N}$ | 9 | 20 |

Suppl. Table 5. GlpK$^{S166L}$ genes induced and repressed in comparison with Bamy.

Excel file.

Suppl. Table 6. Strains used in this study.

| ***Strain*** | ***Genotype*** | ***Reference*** |
|---|---|---|
| ***B. amyloliquefaciens FZB42*** | Wild type | *Bacillus* Genetic Stock Center (BGSC) |
| ***B. amyloliquefaciens FZB42*** | glpK::km | This study |
| ***B. amyloliquefaciens FZB42*** | Spontaneous mutant. *glpK*$^{F38S, W52stop, G254A}$ | This study |
| ***B. amyloliquefaciens FZB42*** | *Spontaneous mutant. glpK*$^{G147R, Q421R}$ | This study |
| ***B. amyloliquefaciens FZB42*** | *Spontaneous mutant. glpK*$^{W355R}$ | This study |
| ***B. amyloliquefaciens FZB42*** | *Spontaneous mutant. glpK*$^{S166L}$ | This study |
| ***B. amyloliquefaciens FZB42*** | *Spontaneous mutant. glpK*$^{+}$ | This study |
| ***B. amyloliquefaciens FZB42*** | *Spontaneous mutant. glpK*$^{G162R, W485stop}$ | This study |
| ***B. amyloliquefaciens FZB42*** | *Spontaneous mutant. glpK*$^{Q246-, G265S}$ | This study |
| ***B. amyloliquefaciens FZB42*** | Wild type, amyE:: Pveg-yfp | This study |
| ***B. amyloliquefaciens FZB42*** | ΔfenA | *Bacillus* Genetic Stock Center (BGSC) |
| ***B. amyloliquefaciens FZB42*** | ΔbaeJ | BGSC |

| | | |
|---|---|---|
| ***B. amyloliquefaciens FZB42*** | ΔbmyA | BGSC |
| ***B. amyloliquefaciens FZB42*** | Δdfn | BGSC |
| ***B. amyloliquefaciens FZB42*** | ΔsrfA | BGSC |
| ***P. chlororaphis PCL1606*** | Wild type | [1] |
| ***P. chlororaphis PCL1606*** | ΔPCL1606_RS10280 (PvdO) | This study |
| ***P. chlororaphis PCL1606*** | ΔPCL1606_RS14085 (AcsD) | This study |
| ***P. chlororaphis PCL1606*** | ΔPCL1606_RS13600 (PchC) | This study |
| ***P. chlororaphis PCL1606*** | ΔPCL1606_RS05710 | This study |
| ***P. chlororaphis PCL1606*** | ΔPCL1606_RS13355 | This study |
| ***P. chlororaphis PCL1606*** | ΔPCL1606_RS16180 | This study |
| ***P. chlororaphis PCL1606*** | ΔPCL1606_RS15365 | This study |
| ***P. chlororaphis PCL1606*** | ΔPCL1606_RS16050 | This study |
| ***P. chlororaphis PCL1606*** | ΔPCL1606_RS23355 | This study |
| ***P. chlororaphis PCL1606*** | ΔPCL1606_RS08425 | This study |
| ***P. chlororaphis PCL1606*** | ΔPCL1606_RS13600 (PchC), ΔPCL1606_RS14085 (AcsD), ΔPCL1606_RS10280 (PvdO) | This study |
| ***P. chlororaphis PCL1606*** | ΔPCL1606_RS13600 (PchC), ΔPCL1606_RS10280 (PvdO) | This study |
| ***P. chlororaphis PCL1606*** | ΔPCL1606_RS10170-RS10180 (DarA-DarB-DarC) | This study |
| ***P. chlororaphis PCL1606*** | ΔPCL1606_RS10170-RS10180 (DarA-DarB-DarC), ΔPCL1606_RS10280 (PvdO) | This study |
| ***P. chlororaphis PCL1606*** | Spontaneous mutant. *fusA*$^{T680D}$ | This study |
| ***P. chlororaphis PCL1606*** | Spontaneous mutant. *fusA*$^{T680D}$ | This study |

| | | |
|---|---|---|
| ***P. chlororaphis PCL1606*** | Spontaneous mutant. *fusA*$^{K366N}$ | This study |

Suppl. Table 7. Oligonucleotides used in this study.

| Name | Sequence |
|---|---|
| T2SS_up_Fwd | agtatagggataacagggtaatctgcaggacccggacatcatcatgatcggc |
| T2SS_up_Rev | cgataacggttcacaggcgtcccgggtcacacggagga |
| T2SS_down_Fwd | cccgggacgcctgtgaaccgttatcgctacgaagccgc |
| T2SS_down_Rev | agaggatccccgggtaccgagctcgcgtgaggatcttgccacgcagggtg |
| RS05710_up_Fwd | tctgaattcgagctcggtacccgggaccaccggcgggcataagttgtcgtaatt |
| RS05710_up_Rev | tgcagctgcatgagtcccggaaggggccttcgaccata |
| RS05710_down_Fwd | ccccttccgggactcatgcagctgcaattcggagccga |
| RS05710_down_Rev | gcatgcctgcaggtcgactctagagcagctgggtgcggatgaagtgggaaaag |
| UP_RS14085_fwd | agtatagggataacagggtaatctgcgcgggctgggcctgatgctgggca |
| UP_RS14085_rev | ggtcgccgcgccaaaaggctccacagtagccatgcgatactggccacg |
| Down_RS14085_fwd | tgtggagccttttggcgcggcgacctgccgctgctcga |
| Down_RS14085_rev | agaggatccccgggtaccgagctcgcaactgagtgcgcagacgcaccacccgc |
| RS13600-up_fwd | agtatagggataacagggtaatctgcgatgaggtgaactcgccgtcattg |
| RS13600-up_rev | aagggtttttccgggatctcctatttgtagaaataattggctacaaatg |
| RS13600-DOWN_fwd | aataggagatcccggaaaaacccttccatgaaaacccctgac |
| RS13600-DOWN_rev | agaggatccccgggtaccgagctcgagaacaggctttcggcgtggggctg |
| UP_RS10280_Fwd | agtatagggataacagggtaatctgtctgcgcccgcagcacggagctcaa |
| UP_RS10280_Rev | ggcggtgcggggggggtcgtctcgaaggttggaaaaggtggatcgc |
| DOWN_RS10280_Fwd | ttcgagacgaccccccgcaccgccccacgccccgga |
| DOWN_RS10280_Rev | Agaggatccccgggtaccgagctcgggcagcggctacagatgcgagg cgttaacacgacatctgtaggagc |
| UP_RS10170_Fwd | tctgaattcgagctcggtacccggggtcgaaagagcgggcgccacgatgc |
| UP_RS10170_Rev | acaggaccggcgaaatacggggaccttttgctttacaaccacaaag |
| DOWN_RS10170_Fwd | ggtccccgtatttcgccggtcctgtgaagatgcggccg |
| DOWN_RS10170_Rev | gcatgcctgcaggtcgactctagagctatgtcacctcggtgtgcaccggc |
| RS15365_UP_Fwd | tctgaattcgagctcggtacccggggagagcgcaatagagcagccaggac |
| RS15365_UP_Rev | tggcctcccttaatgacgctccataaattaatcctacagtgag |
| RS15365_Down_Fwd | tatggagcgtcattaagggaggccaggcagcggccgtc |
| RS15365_Down_Rev | gcatgcctgcaggtcgactctagagtcagcagcctatccatgggagtgaagaccag |
| bact_RS23355_UP_fwd | tctgaattcgagctcggtacccgggcaggccggtgtgcgtcgtgctgttg |
| bact_RS23355_UP_rev | cggcttgggtatcgacactcaggcggacggtggattac |
| bact_RS23355_DOWN_fwd | cgcctgagtgtcgatacccaagccgccttcgccagcgc |
| bact_RS23355_DOWN_rev | gcatgcctgcaggtcgactctagagggcccatagcgcgaccactggatgag |
| GlpK_up_Fwd | tctgcagacgcgtcgacgtcatatgtttagatatccacctcggtcaattaaaag |
| GlpK_up_Rev | aaatggttcgctggatgccgctctcctttttaaatatattc |

| | |
|---|---|
| Km_GlpK_Fwd | gagagcggcatccagcgaaccatttgaggtgataggtaag |
| Km_GlpK_Rev | cacatttttatccgatacaaattcctcgtaggcgctc |
| GlpK_down_Fwd | ggaatttgtatcggataaaaatgtggtatactgaaaacaagttaatag |
| GlpK_down_Rev | tccagcctcgcgtcgggcgatatcgtagagccgtatctgatggctaactg |
| RS16180_UP_fwd | attcgagctcggtacccgggaagtgccgaggccgaaggaaggaat |
| RS16180_UP_rev | cgacaacttctgaatcattccaagttctctagttgcgaaaaac |
| RS16180_DOWN_fwd | gaatgattcagaagttgtcgttggcgtgcagcgcc |
| RS16180_DOWN_rev | cctgcaggtcgactctagaggttgattccctcttataaccaccgttttctactgg |
| RS08425_UP_fwd | tctagagtcgacctgcaggcatgcactgatgcgatcggcctcggccccca |
| RS08425_UP_rev | tgttgccctgaggcaactctcctgctaccacgctgttttgaatgaac |
| RS08425_DOWN_fwd | gcaggagagttgcctcagggcaacaccccggctgcccc |
| RS08425_DOWN_rev | aaaaaagaatatataaggcttttaattccccaccaggtcgaaaccggtcagg |

### Supplementary Movies

Suppl. Movie 1. Movie of the interaction between Pcl and Bamy acquired via inverted microscopy over 120 h.

Suppl. Movie 2. Movie of the interaction between ΔGacS and Bamy acquired via inverted microscopy over 72 h.

Suppl. Movie 3. Movie of the interaction between Pcl and Δbae acquired via inverted microscopy over 72 h.

Suppl. Movie 4. Movie of the interaction between FusA$^{T680D}$ and Bamy acquired via inverted microscopy over 72 h.

Suppl. Movie 5. Comparison of WT FusA and FusA$^{T680D}$.

Suppl. Movie 6. Movie of the interaction between Pcl and GlpKS166L acquired via inverted microscopy over 72 h.
