## Supplementary Table 2 for "Chemical interplay and complementary adaptative strategies toggle bacterial antagonism and co-existence"

| Gene | Product | logFC | pvalue |
|---|---|---|---|
| RBAM_RS171 | nitrate reductase subunit alpha | 4.15 | 1.7E-103 |
| RBAM_RS094 | MBL fold metallo-hydrolase | 3.79 | 1.3E-68 |
| RBAM_RS094 | transcriptional regulator | 3.72 | 1.6E-56 |
| RBAM_RS171 | nitrate reductase subunit beta | 3.69 | 3.4E-61 |
| RBAM_RS094 | AraC family transcriptional regulator | 3.46 | 4.0E-79 |
| RBAM_RS171 | nitrate reductase molybdenum cofactor assembly chaperc | 3.46 | 2.5E-56 |
| RBAM_RS045 | sulfonate ABC transporter substrate-binding protein | 3.38 | 2.6E-28 |
| RBAM_RS171 | nitrate reductase subunit gamma | 3.37 | 5.3E-51 |
| RBAM_RS028 | transcriptional regulator | 3.28 | 3.4E-45 |
| RBAM_RS045 | aliphatic sulfonates ABC transporter ATP-binding protein S | 3.24 | 1.0E-39 |
| RBAM_RS045 | sulfonate ABC transporter permease | 3.02 | 6.3E-30 |
| RBAM_RS171 | transcriptional regulator | 2.93 | 2.1E-30 |
| RBAM_RS094 | tRNA-binding protein | 2.75 | 3.9E-70 |
| RBAM_RS172 | MFS transporter | 2.6 | 1.3E-33 |
| RBAM_RS045 | alkanesulfonate monooxygenase | 2.5 | 8.8E-33 |
| RBAM_RS121 | hypothetical protein | 2.45 | 9.4E-37 |
| RBAM_RS121 | formate dehydrogenase subunit alpha | 2.4 | 3.6E-49 |
| RBAM_RS153 | N-acetyltransferase | 2.2 | 1.8E-14 |
| RBAM_RS153 | amino acid ABC transporter substrate-binding protein | 2.13 | 7.4E-18 |
| RBAM_RS153 | cysteine ABC transporter permease | 2.1 | 1.1E-14 |
| RBAM_RS153 | arginine ABC transporter ATP-binding protein | 2.05 | 4.9E-14 |
| RBAM_RS028 | dihydrodipicolinate synthase family protein | 2.01 | 4.9E-18 |
| RBAM_RS153 | LLM class flavin-dependent oxidoreductase | 1.95 | 2.0E-18 |
| RBAM_RS192 | GapA-binding peptide SR1P | 1.89 | 6.8E-20 |
| RBAM_RS072 | polysaccharide deacetylase | 1.87 | 3.0E-30 |
| RBAM_RS067 | cell wall hydrolase | 1.85 | 4.6E-13 |
| RBAM_RS153 | NrdH-redoxin | 1.82 | 5.1E-10 |
| RBAM_RS022 | D-lyxose isomerase | 1.72 | 3.1E-30 |
| RBAM_RS153 | amino acid ABC transporter substrate-binding protein | 1.65 | 3.5E-11 |
| RBAM_RS045 | 2-amino-3-carboxymuconate-6-semialdehyde decarboxyla | 1.64 | 2.2E-17 |
| RBAM_RS140 | nitric oxide dioxygenase | 1.61 | 3.9E-15 |
| RBAM_RS053 | hypothetical protein | 1.55 | 5.5E-06 |
| RBAM_RS145 | NADH dehydrogenase | 1.54 | 2.3E-22 |
| RBAM_RS145 | hypothetical protein | 1.49 | 7.1E-07 |
| RBAM_RS142 | DUF378 domain-containing protein | 1.47 | 7.1E-19 |
| RBAM_RS153 | cysteine ABC transporter permease | 1.42 | 4.9E-07 |
| RBAM_RS153 | LLM class flavin-dependent oxidoreductase | 1.41 | 5.2E-09 |
| RBAM_RS029 | EamA family transporter | 1.39 | 2.6E-21 |
| RBAM_RS044 | tRNA-Leu | 1.39 | 8.2E-05 |
| RBAM_RS057 | GTP-binding protein | 1.39 | 8.0E-19 |
| RBAM_RS051 | hypothetical protein | 1.36 | 5.4E-22 |
| RBAM_RS071 | hypothetical protein | 1.34 | 1.0E-21 |
| RBAM_RS184 | fatty acid desaturase | 1.34 | 3.8E-13 |
| RBAM_RS173 | membrane protein | 1.32 | 3.4E-20 |
| RBAM_RS178 | PTS sugar transporter subunit IIB | 1.32 | 1.9E-05 |
| RBAM_RS155 | choline ABC transporter permease | 1.28 | 2.8E-09 |
| RBAM_RS100 | acetyltransferase | 1.25 | 7.7E-17 |

| | | | |
|---|---|---|---|
| RBAM_RS178 | PTS cellobiose transporter subunit IIA | 1.25 | 1.4E-06 |
| RBAM_RS022 | NAD(P)-dependent oxidoreductase | 1.24 | 1.0E-18 |
| RBAM_RS121 | cystathionine gamma-synthase | 1.23 | 1.3E-14 |
| RBAM_RS122 | AI-2E family transporter | 1.22 | 3.2E-05 |
| RBAM_RS179 | cytochrome ubiquinol oxidase subunit I | 1.22 | 2.4E-04 |
| RBAM_RS085 | sporulation protein | 1.15 | 2.2E-04 |
| RBAM_RS190 | hypothetical protein | 1.15 | 3.8E-04 |
| RBAM_RS036 | hypothetical protein | 1.14 | 2.0E-09 |
| RBAM_RS073 | hypothetical protein | 1.14 | 8.6E-08 |
| RBAM_RS023 | manganese catalase | 1.12 | 1.5E-13 |
| RBAM_RS138 | tRNA-Glu | 1.11 | 7.3E-05 |
| RBAM_RS176 | GlsB/YeaQ/YmgE family stress response membrane protei | 1.11 | 1.5E-16 |
| RBAM_RS185 | hypothetical protein | 1.11 | 7.1E-06 |
| RBAM_RS121 | formate/nitrite transporter | 1.1 | 1.1E-10 |
| RBAM_RS084 | polyketide synthase | 1.08 | 2.2E-11 |
| RBAM_RS155 | amino acid ABC transporter permease | 1.08 | 8.5E-08 |
| RBAM_RS094 | peptidoglycan-binding protein | 1.07 | 6.5E-14 |
| RBAM_RS155 | osmoprotectant ABC transporter substrate-binding protei | 1.07 | 2.5E-08 |
| RBAM_RS109 | polyketide synthase | 1.05 | 5.6E-15 |
| RBAM_RS031 | tRNA-Asp | 1.04 | 8.1E-05 |
| RBAM_RS178 | PTS system, cellobiose-specific IIC component | 1.03 | 6.3E-07 |
| RBAM_RS038 | NAD(P)-dependent oxidoreductase | 1.02 | 6.2E-08 |
| RBAM_RS109 | polyketide synthase | 1.02 | 9.0E-13 |
| RBAM_RS152 | sulfite reductase [NADPH] flavoprotein, alpha-component | 1.02 | 6.5E-13 |
| RBAM_RS084 | non-ribosomal peptide synthetase | 1.01 | 2.1E-11 |
| RBAM_RS094 | hypothetical protein | 1 | 3.3E-07 |
| RBAM_RS160 | imidazole glycerol phosphate synthase subunit HisH | -1.01 | 6.5E-08 |
| RBAM_RS173 | hypothetical protein | -1.01 | 3.0E-07 |
| RBAM_RS173 | hypothetical protein | -1.01 | 2.8E-07 |
| RBAM_RS034 | phosphoribosylformylglycinamidine synthase, purS protei | -1.02 | 7.3E-06 |
| RBAM_RS129 | antiholin | -1.02 | 1.6E-09 |
| RBAM_RS173 | hypothetical protein | -1.02 | 1.3E-07 |
| RBAM_RS034 | phosphoribosylaminoimidazolesuccinocarboxamide synth | -1.05 | 2.8E-08 |
| RBAM_RS034 | phosphoribosylformylglycinamidine synthase subunit PurL | -1.05 | 1.9E-07 |
| RBAM_RS067 | hypothetical protein | -1.05 | 1.3E-08 |
| RBAM_RS013 | glycerol-3-phosphate transporter | -1.06 | 1.2E-11 |
| RBAM_RS017 | proline dehydrogenase | -1.07 | 8.8E-10 |
| RBAM_RS021 | hypothetical protein | -1.08 | 8.2E-14 |
| RBAM_RS172 | LuxR family transcriptional regulator | -1.09 | 6.6E-11 |
| RBAM_RS021 | hypothetical protein | -1.1 | 9.4E-13 |
| RBAM_RS104 | stage IV sporulation protein A | -1.1 | 3.8E-07 |
| RBAM_RS158 | transcriptional regulator | -1.1 | 2.5E-09 |
| RBAM_RS172 | hypothetical protein | -1.1 | 2.5E-11 |
| RBAM_RS181 | pectate lyase | -1.1 | 3.4E-14 |
| RBAM_RS021 | copper transporter | -1.11 | 5.6E-15 |
| RBAM_RS034 | phosphoribosylformylglycinamidine synthase subunit Pur[illegible] | -1.11 | 8.4E-08 |
| RBAM_RS160 | bifunctional phosphoribosyl-AMP cyclohydrolase/phospho | -1.11 | 2.3E-08 |

| | | | |
|---|---|---|---|
| RBAM_RS076 | dihydroorotate dehydrogenase | -1.13 | 5.4E-05 |
| RBAM_RS155 | UDP-glucose 6-dehydrogenase | -1.13 | 1.5E-12 |
| RBAM_RS160 | imidazole glycerol phosphate synthase cyclase subunit | -1.13 | 4.1E-10 |
| RBAM_RS046 | sporulation protein YhbH | -1.14 | 1.8E-05 |
| RBAM_RS137 | MFS transporter | -1.14 | 1.9E-16 |
| RBAM_RS160 | imidazoleglycerol-phosphate dehydratase | -1.14 | 1.3E-09 |
| RBAM_RS031 | QacE family quaternary ammonium compound efflux SMR | -1.16 | 2.5E-08 |
| RBAM_RS015 | stress protein | -1.17 | 2.7E-20 |
| RBAM_RS140 | flotillin family protein | -1.17 | 1.1E-11 |
| RBAM_RS160 | ATP phosphoribosyltransferase regulatory subunit | -1.17 | 1.7E-10 |
| RBAM_RS160 | histidinol dehydrogenase | -1.18 | 3.0E-12 |
| RBAM_RS172 | hypothetical protein | -1.18 | 1.8E-13 |
| RBAM_RS160 | ATP phosphoribosyltransferase | -1.19 | 1.2E-09 |
| RBAM_RS094 | hypothetical protein | -1.21 | 7.3E-05 |
| RBAM_RS186 | hypothetical protein | -1.22 | 5.6E-08 |
| RBAM_RS076 | orotidine-5'-phosphate decarboxylase | -1.25 | 4.9E-06 |
| RBAM_RS160 | 1-(5-phosphoribosyl)-5-((5- phosphoribosylamino)methyli | -1.25 | 2.4E-11 |
| RBAM_RS010 | alcohol dehydrogenase | -1.27 | 3.5E-14 |
| RBAM_RS131 | sporulation protein YtfJ | -1.31 | 4.2E-05 |
| RBAM_RS158 | hypothetical protein | -1.35 | 2.2E-21 |
| RBAM_RS030 | penicillin-binding protein 4 | -1.36 | 5.6E-20 |
| RBAM_RS076 | orotate phosphoribosyltransferase | -1.36 | 9.2E-08 |
| RBAM_RS151 | DNA-directed RNA polymerase sigma-70 factor | -1.36 | 1.4E-08 |
| RBAM_RS172 | transcriptional regulator | -1.36 | 2.8E-20 |
| RBAM_RS010 | hypothetical protein | -1.44 | 1.7E-12 |
| RBAM_RS010 | aspartate aminotransferase family protein | -1.48 | 4.3E-19 |
| RBAM_RS190 | hypothetical protein | -1.52 | 1.2E-11 |
| RBAM_RS176 | hypothetical protein | -1.62 | 1.3E-09 |
| RBAM_RS036 | hypothetical protein | -1.71 | 1.1E-05 |
| RBAM_RS094 | hypothetical protein | -1.74 | 3.0E-07 |
| RBAM_RS172 | hypothetical protein | -1.79 | 3.8E-33 |
| RBAM_RS088 | hypothetical protein | -1.81 | 9.1E-35 |
| RBAM_RS140 | membrane protein | -1.94 | 3.1E-25 |
| RBAM_RS172 | hypothetical protein | -1.97 | 4.4E-24 |
| RBAM_RS174 | L-glutamate gamma-semialdehyde dehydrogenase | -2.54 | 2.4E-34 |
| RBAM_RS174 | glutamate dehydrogenase | -3.05 | 5.0E-22 |
| RBAM_RS017 | cobalamin synthesis protein CobW | -3.7 | 2.1E-21 |
