## Supplementary Table 5 for "Chemical interplay and complementary adaptative strategies toggle bacterial antagonism and co-existence"

| Gene | Product | log2FoldChange |
|---|---|---|
| RBAM_RS04200 | acetoin:2,6-dichlorophenolindophenol oxidoreductas | 11.28266821 |
| RBAM_RS04195 | acetoin:2,6-dichlorophenolindophenol oxidoreductas | 10.78487146 |
| RBAM_RS04205 | 2-oxo acid dehydrogenase subunit E2 | 10.54284658 |
| RBAM_RS04210 | dihydrolipoyl dehydrogenase | 9.458596041 |
| RBAM_RS04235 | PTS transporter subunit EIIC | 8.508908274 |
| RBAM_RS18535 | MFS transporter | 8.333327335 |
| RBAM_RS08485 | L-threonine 3-dehydrogenase | 7.64702645 |
| RBAM_RS18540 | myo-inosose-2 dehydratase | 7.201325394 |
| RBAM_RS18530 | Gfo/Idh/MocA family oxidoreductase | 7.193457299 |
| RBAM_RS18525 | sugar phosphate isomerase/epimerase | 6.951031532 |
| RBAM_RS04225 | 6-phospho-alpha-glucosidase | 6.902915531 |
| RBAM_RS13960 | phosphoenolpyruvate carboxykinase (ATP) | 6.780836385 |
| RBAM_RS13185 | glyceraldehyde-3-phosphate dehydrogenase | 6.707171656 |
| RBAM_RS13075 | L-ribulose-5-phosphate 4-epimerase | 6.490863971 |
| RBAM_RS18545 | 3D-(3,5/4)-trihydroxycyclohexane-1,2-dione acylhydr | 6.436260027 |
| RBAM_RS18520 | 2-keto-myo-inositol isomerase | 6.330894914 |
| RBAM_RS08490 | glycine C-acetyltransferase | 6.28131839 |
| RBAM_RS13050 | carbohydrate ABC transporter permease | 6.017396777 |
| RBAM_RS13760 | carbohydrate ABC transporter substrate-binding prot | 5.930713602 |
| RBAM_RS14420 | DUF378 domain-containing protein | 5.858032429 |
| RBAM_RS04215 | sigma-54-dependent Fis family transcriptional regula | 5.833035913 |
| RBAM_RS18550 | 5-dehydro-2-deoxygluconokinase | 5.793023982 |
| RBAM_RS18075 | PTS cellobiose transporter subunit IIC | 5.727208029 |
| RBAM_RS18515 | class II fructose-1,6-bisphosphate aldolase | 5.678726025 |
| RBAM_RS09255 | AMP-binding protein | 5.654327745 |
| RBAM_RS09235 | enoyl-CoA hydratase | 5.598682398 |
| RBAM_RS18385 | urocanate hydratase | 5.569161095 |
| RBAM_RS13755 | LacI family DNA-binding transcriptional regulator | 5.456259052 |
| RBAM_RS09850 | aspartate aminotransferase family protein | 5.437264486 |
| RBAM_RS13080 | ribulokinase | 5.433823737 |
| RBAM_RS15845 | sugar porter family MFS transporter | 5.402513779 |
| RBAM_RS09250 | acetyl-CoA carboxylase biotin carboxylase subunit | 5.395387877 |
| RBAM_RS04230 | MurR/RpiR family transcriptional regulator | 5.186606078 |
| RBAM_RS18380 | histidine ammonia-lyase | 5.143203447 |
| RBAM_RS18070 | PTS lactose/cellobiose transporter subunit IIA | 5.142151129 |
| RBAM_RS09175 | CoA transferase subunit A | 5.128170518 |
| RBAM_RS13765 | sugar ABC transporter permease | 5.098723989 |
| RBAM_RS18390 | imidazolonepropionase | 5.062203649 |
| RBAM_RS13070 | HAD-IIA family hydrolase | 5.012515102 |
| RBAM_RS13055 | sugar ABC transporter permease | 4.936423666 |
| RBAM_RS03315 | MFS transporter | 4.927784379 |
| RBAM_RS18065 | 6-phospho-beta-glucosidase | 4.924612943 |
| RBAM_RS04880 | mechanosensitive ion channel | 4.911092781 |
| RBAM_RS18395 | formimidoylglutamase | 4.895193556 |
| RBAM_RS18080 | PTS sugar transporter subunit IIB | 4.874130057 |
| RBAM_RS13575 | CBS domain-containing protein | 4.855155268 |
| RBAM_RS09840 | 3-oxoacid CoA-transferase subunit B | 4.854683639 |

| | | |
|---|---|---|
| RBAM_RS02220 | D-lyxose/D-mannose family sugar isomerase | 4.799814211 |
| RBAM_RS10465 | ubiquinol-cytochrome c reductase iron-sulfur subunit | 4.750903905 |
| RBAM_RS09835 | peptidase | 4.67191888 |
| RBAM_RS09595 | aldehyde dehydrogenase family protein | 4.652499038 |
| RBAM_RS09180 | GntP family permease | 4.645001611 |
| RBAM_RS11340 | 3-hydroxybutyryl-CoA dehydrogenase | 4.58137628 |
| RBAM_RS09165 | 3-hydroxybutyrate dehydrogenase | 4.567805974 |
| RBAM_RS15940 | L-lactate permease | 4.562626166 |
| RBAM_RS01790 | glutathione-dependent formaldehyde dehydrogenase | 4.489854221 |
| RBAM_RS02215 | SDR family oxidoreductase | 4.467657933 |
| RBAM_RS17115 | hypothetical protein | 4.456860783 |
| RBAM_RS11335 | acyl-CoA dehydrogenase | 4.446375039 |
| RBAM_RS10460 | cytochrome b6 | 4.427602601 |
| RBAM_RS09845 | CoA transferase subunit A | 4.373957795 |
| RBAM_RS03175 | sugar porter family MFS transporter | 4.366337208 |
| RBAM_RS13090 | glycoside hydrolase family 43 protein | 4.334781076 |
| RBAM_RS16745 | sugar ABC transporter ATP-binding protein | 4.276531682 |
| RBAM_RS08800 | alcohol dehydrogenase AdhP | 4.271470691 |
| RBAM_RS18010 | PTS cellobiose transporter subunit IIC | 4.269480779 |
| RBAM_RS15470 | hypothetical protein | 4.232617661 |
| RBAM_RS13085 | L-arabinose isomerase | 4.216433221 |
| RBAM_RS09830 | putative beta-lysine N-acetyltransferase | 4.21215083 |
| RBAM_RS13060 | carbohydrate ABC transporter substrate-binding prot | 4.208933777 |
| RBAM_RS11330 | citrate synthase | 4.203809208 |
| RBAM_RS08760 | xylose isomerase | 4.181170356 |
| RBAM_RS16105 | MFS transporter | 4.180968898 |
| RBAM_RS03955 | citrate transporter | 4.140136133 |
| RBAM_RS02390 | manganese catalase family protein | 4.137658138 |
| RBAM_RS09265 | family 10 glycosylhydrolase | 4.133772612 |
| RBAM_RS11500 | stage III sporulation protein AA | 4.126705656 |
| RBAM_RS03960 | MBL fold metallo-hydrolase | 4.117577643 |
| RBAM_RS06150 | PTS transporter subunit EIIC | 4.104025796 |
| RBAM_RS14765 | NAD(P)/FAD-dependent oxidoreductase | 4.102297961 |
| RBAM_RS13580 | acetoin utilization protein AcuC | 4.095216819 |
| RBAM_RS18355 | PTS glucose transporter subunit IIA | 4.076640326 |
| RBAM_RS11345 | acetyl-CoA C-acetyltransferase | 4.052326809 |
| RBAM_RS09745 | hypothetical protein | 4.026368512 |
| RBAM_RS09240 | hydroxymethylglutaryl-CoA lyase | 4.019074881 |
| RBAM_RS17760 | DegT/DnrJ/EryC1/StrS family aminotransferase | 4.015651066 |
| RBAM_RS17845 | hypothetical protein | 3.984155385 |
| RBAM_RS13065 | sn-glycerol-1-phosphate dehydrogenase | 3.974721208 |
| RBAM_RS01560 | alpha-glucosidase | 3.913191808 |
| RBAM_RS00105 | glycoside hydrolase family 18 protein | 3.870093007 |
| RBAM_RS12900 | acyl-CoA thioesterase | 3.844366269 |
| RBAM_RS13770 | carbohydrate ABC transporter permease | 3.838160957 |
| RBAM_RS16750 | ribose ABC transporter permease | 3.836989488 |
| RBAM_RS11485 | stage III sporulation protein AD | 3.814640638 |

| | | |
|---|---|---|
| RBAM_RS13570 | GNAT family N-acetyltransferase | 3.7778736 |
| RBAM_RS04035 | PTS system trehalose-specific EIIBC component | 3.764016072 |
| RBAM_RS16740 | D-ribose pyranase | 3.747108918 |
| RBAM_RS07315 | GapA-binding peptide SR1P | 3.714143555 |
| RBAM_RS07310 | polysaccharide deacetylase family protein | 3.68625132 |
| RBAM_RS10595 | DUF2768 domain-containing protein | 3.674998634 |
| RBAM_RS05515 | germination protein GerPB | 3.670134598 |
| RBAM_RS13045 | alpha-N-arabinofuranosidase | 3.649231446 |
| RBAM_RS15745 | NAAT family transporter | 3.642177881 |
| RBAM_RS09230 | acyl-CoA carboxylase subunit beta | 3.597417092 |
| RBAM_RS01650 | starch-binding protein | 3.592294821 |
| RBAM_RS07545 | sporulation-specific transcriptional regulator GerR | 3.588484982 |
| RBAM_RS06155 | PTS lactose/cellobiose transporter subunit IIA | 3.571957851 |
| RBAM_RS06145 | galactokinase | 3.56536441 |
| RBAM_RS06210 | formate dehydrogenase subunit alpha | 3.536687287 |
| RBAM_RS16655 | GerAB/ArcD/ProY family transporter | 3.519726657 |
| RBAM_RS17750 | N-acetylneuraminate synthase family protein | 3.512780509 |
| RBAM_RS05055 | YheC/YheD family protein | 3.512358937 |
| RBAM_RS06885 | redoxin domain-containing protein | 3.497434029 |
| RBAM_RS02410 | dicarboxylate/amino acid:cation symporter | 3.46881222 |
| RBAM_RS01795 | RDD family protein | 3.46760682 |
| RBAM_RS18400 | amino acid permease | 3.460239602 |
| RBAM_RS11495 | stage III sporulation protein SpoAB | 3.444862276 |
| RBAM_RS06205 | YjgB family protein | 3.440415747 |
| RBAM_RS13200 | aldo/keto reductase | 3.371147228 |
| RBAM_RS04125 | calcium/proton exchanger | 3.354897124 |
| RBAM_RS18020 | glycosyltransferase family 8 protein | 3.347368551 |
| RBAM_RS17915 | galactokinase | 3.339137543 |
| RBAM_RS09305 | aldose 1-epimerase | 3.338583333 |
| RBAM_RS06135 | UDP-glucose--hexose-1-phosphate uridylyltransferas | 3.326751897 |
| RBAM_RS17835 | PTS sucrose transporter subunit IIBC | 3.324232457 |
| RBAM_RS06140 | UDP-glucose 4-epimerase GalE | 3.291581799 |
| RBAM_RS15130 | acetyl-CoA C-acetyltransferase | 3.288287798 |
| RBAM_RS18275 | family 16 glycosylhydrolase | 3.287392293 |
| RBAM_RS07450 | cytochrome c oxidase subunit I | 3.283604426 |
| RBAM_RS18205 | sn-glycerol-3-phosphate ABC transporter ATP-bindin | 3.254193042 |
| RBAM_RS07460 | cytochrome c oxidase subunit IVB | 3.202756735 |
| RBAM_RS18140 | ROK family protein | 3.187184232 |
| RBAM_RS06200 | DUF2809 domain-containing protein | 3.162945568 |
| RBAM_RS04040 | alpha,alpha-phosphotrehalase | 3.161482381 |
| RBAM_RS03670 | spore coat protein CotJB | 3.156674902 |
| RBAM_RS14525 | Na+/H+ antiporter subunit A | 3.156163272 |
| RBAM_RS16650 | spore germination protein | 3.15319604 |
| RBAM_RS04685 | sporulation protein YhbH | 3.151080665 |
| RBAM_RS05050 | YheC/YheD family protein | 3.104319944 |
| RBAM_RS06470 | ABC transporter permease | 3.097892027 |
| RBAM_RS13375 | DUF2953 domain-containing protein | 3.094278482 |

| | | |
|---|---|---|
| RBAM_RS01900 | LysR family transcriptional regulator | 3.094243425 |
| RBAM_RS08310 | DUF503 domain-containing protein | 3.070849997 |
| RBAM_RS18170 | NAD(P)H-hydrate dehydratase | 3.06301798 |
| RBAM_RS04680 | serine/threonine protein kinase PrkA | 3.031495957 |
| RBAM_RS03990 | DUF3243 domain-containing protein | 3.028771723 |
| RBAM_RS10455 | c-type cytochrome | 3.027057202 |
| RBAM_RS10585 | stage IV sporulation protein A | 3.022788921 |
| RBAM_RS02205 | transcription antiterminator | 3.018880464 |
| RBAM_RS09825 | lysine 2,3-aminomutase | 2.998918629 |
| RBAM_RS17675 | amino acid permease | 2.989779361 |
| RBAM_RS18375 | hut operon transcriptional regulator HutP | 2.989242548 |
| RBAM_RS14540 | Na+/H+ antiporter subunit D | 2.987673919 |
| RBAM_RS06475 | ABC transporter ATP-binding protein | 2.98170997 |
| RBAM_RS15900 | MurR/RpiR family transcriptional regulator | 2.976550717 |
| RBAM_RS17745 | spore coat polysaccharide biosynthesis protein SpsF | 2.952608119 |
| RBAM_RS00385 | serine/threonine protein kinase | 2.947203126 |
| RBAM_RS04885 | SpoVR family protein | 2.943600075 |
| RBAM_RS04820 | glycerol-3-phosphate responsive antiterminator | 2.920364638 |
| RBAM_RS16755 | ribose ABC transporter substrate-binding protein Rbs | 2.907325304 |
| RBAM_RS07465 | cytochrome c oxidase assembly factor CtaG | 2.887579176 |
| RBAM_RS03665 | spore coat associated protein CotJA | 2.886325077 |
| RBAM_RS01450 | ring-cleaving dioxygenase | 2.885916453 |
| RBAM_RS17920 | GtrA family protein | 2.878150807 |
| RBAM_RS06130 | glycosyl hydrolase 53 family protein | 2.87407857 |
| RBAM_RS11480 | stage III sporulation protein AE | 2.865867537 |
| RBAM_RS16670 | sugar porter family MFS transporter | 2.864771643 |
| RBAM_RS09170 | CoA transferase subunit B | 2.857709111 |
| RBAM_RS09430 | hypothetical protein | 2.847387045 |
| RBAM_RS06565 | hypothetical protein | 2.841354028 |
| RBAM_RS07810 | calcium-translocating P-type ATPase, SERCA-type | 2.830490293 |
| RBAM_RS03740 | cytochrome P450 | 2.826896677 |
| RBAM_RS05815 | putative glycoside hydrolase | 2.821953642 |
| RBAM_RS14545 | Na+/H+ antiporter subunit E | 2.821439367 |
| RBAM_RS07065 | mechanosensitive ion channel family protein | 2.810795487 |
| RBAM_RS03310 | sorbitol dehydrogenase | 2.805112064 |
| RBAM_RS16730 | LacI family DNA-binding transcriptional regulator | 2.79888806 |
| RBAM_RS12945 | electron transfer flavoprotein subunit alpha/FixB fam | 2.797377712 |
| RBAM_RS06815 | membrane protein | 2.795807633 |
| RBAM_RS10760 | spore maturation protein SpmB | 2.768453234 |
| RBAM_RS06215 | DUF1641 domain-containing protein | 2.751077997 |
| RBAM_RS10590 | hypothetical protein | 2.737689452 |
| RBAM_RS17910 | UDP-glucose--hexose-1-phosphate uridylyltransferase | 2.736458184 |
| RBAM_RS18555 | 5-deoxy-glucuronate isomerase | 2.730765571 |
| RBAM_RS13040 | carbon starvation protein A | 2.719550261 |
| RBAM_RS04130 | YfkD famly protein | 2.718151845 |
| RBAM_RS05970 | DUF1360 domain-containing protein | 2.713687229 |
| RBAM_RS05505 | hypothetical protein | 2.711451501 |

| | | |
|---|---|---|
| RBAM_RS01730 | proline dehydrogenase | 2.710122325 |
| RBAM_RS13320 | YtpI family protein | 2.709084649 |
| RBAM_RS00380 | VWA domain-containing protein | 2.705001676 |
| RBAM_RS12410 | transporter substrate-binding domain-containing pro | 2.702078134 |
| RBAM_RS17360 | cardiolipin synthase | 2.691668587 |
| RBAM_RS04070 | type 1 glutamine amidotransferase | 2.689765168 |
| RBAM_RS07550 | N-acetyltransferase | 2.682624048 |
| RBAM_RS12865 | metallophosphoesterase | 2.67997393 |
| RBAM_RS07635 | S8 family serine peptidase | 2.671252968 |
| RBAM_RS02290 | pyruvate oxidase | 2.666709009 |
| RBAM_RS09055 | zinc-binding alcohol dehydrogenase family protein | 2.666062524 |
| RBAM_RS18690 | NTP transferase domain-containing protein | 2.664929492 |
| RBAM_RS04700 | DHA2 family efflux MFS transporter permease subun | 2.653352429 |
| RBAM_RS01085 | PTS transporter subunit EIIC | 2.642066505 |
| RBAM_RS06880 | polysaccharide biosynthesis protein | 2.584361211 |
| RBAM_RS07455 | cytochrome (ubi)quinol oxidase subunit III | 2.571681544 |
| RBAM_RS14535 | Na(+)/H(+) antiporter subunit C | 2.56889965 |
| RBAM_RS10765 | spore maturation protein | 2.568601537 |
| RBAM_RS18265 | catalase | 2.567412969 |
| RBAM_RS09065 | mannonate dehydratase | 2.563011359 |
| RBAM_RS14355 | alpha-glucosidase | 2.560330032 |
| RBAM_RS15490 | ATP-binding domain-containing protein | 2.558568103 |
| RBAM_RS12710 | SPOR domain-containing protein | 2.55724292 |
| RBAM_RS18755 | hypothetical protein | 2.553942682 |
| RBAM_RS14550 | Na(+)/H(+) antiporter subunit F1 | 2.546170303 |
| RBAM_RS06480 | peptide ABC transporter substrate-binding protein | 2.534694607 |
| RBAM_RS13035 | (Fe-S)-binding protein | 2.526309907 |
| RBAM_RS18750 | hypothetical protein | 2.526288758 |
| RBAM_RS04760 | hypothetical protein | 2.524374768 |
| RBAM_RS08745 | MFS transporter | 2.521152755 |
| RBAM_RS19130 | VOC family protein | 2.519588799 |
| RBAM_RS05610 | cytochrome P450 | 2.514241473 |
| RBAM_RS09425 | hypothetical protein | 2.509787435 |
| RBAM_RS05695 | carbamoyl phosphate synthase large subunit | 2.490054474 |
| RBAM_RS06465 | ABC transporter permease | 2.489616008 |
| RBAM_RS18150 | mannan endo-1,4-beta-mannosidase | 2.488108265 |
| RBAM_RS06160 | 6-phospho-beta-galactosidase | 2.486201061 |
| RBAM_RS10895 | anti-sigma F factor | 2.483415245 |
| RBAM_RS08350 | YlmC/YmxH family sporulation protein | 2.474648061 |
| RBAM_RS04025 | spore germination protein | 2.473165413 |
| RBAM_RS05260 | YhgE/Pip domain-containing protein | 2.45987614 |
| RBAM_RS11865 | cytochrome c | 2.457042186 |
| RBAM_RS02275 | cellulose biosynthesis cyclic di-GMP-binding regulato | 2.454933906 |
| RBAM_RS18155 | catalase | 2.443912433 |
| RBAM_RS01465 | DUF2512 family protein | 2.421301311 |
| RBAM_RS01290 | hypothetical protein | 2.420953739 |
| RBAM_RS16290 | N-acetylglucosamine-6-phosphate deacetylase | 2.420231092 |

| | | |
|---|---|---|
| RBAM_RS15395 | iron ABC transporter permease | 2.415801585 |
| RBAM_RS05945 | spore coat protein | 2.409631372 |
| RBAM_RS11290 | dihydrolipoyl dehydrogenase | 2.405793174 |
| RBAM_RS11655 | HlyC/CorC family transporter | 2.402560374 |
| RBAM_RS04910 | SDR family oxidoreductase | 2.398564229 |
| RBAM_RS09050 | sugar kinase | 2.396102143 |
| RBAM_RS00925 | polysaccharide deacetylase family sporulation protei | 2.393083455 |
| RBAM_RS16145 | SulP family inorganic anion transporter | 2.392500429 |
| RBAM_RS17560 | RsfA family transcriptional regulator | 2.390650926 |
| RBAM_RS06575 | glycosyltransferase | 2.38849582 |
| RBAM_RS06660 | MarR family transcriptional regulator | 2.385512526 |
| RBAM_RS00400 | ATP-dependent zinc metalloprotease FtsH | 2.380097095 |
| RBAM_RS12415 | amino acid ABC transporter permease | 2.378249157 |
| RBAM_RS18810 | ABC transporter substrate-binding protein | 2.373825508 |
| RBAM_RS13370 | sporulation protein YtfJ | 2.364225595 |
| RBAM_RS16365 | MFS transporter | 2.3622329 |
| RBAM_RS06560 | organic hydroperoxide resistance protein | 2.361299095 |
| RBAM_RS16560 | glycosyltransferase family 4 protein | 2.361016566 |
| RBAM_RS05305 | fatty acid--CoA ligase family protein | 2.345840993 |
| RBAM_RS05025 | ATP-binding cassette domain-containing protein | 2.345739201 |
| RBAM_RS12940 | alpha-N-arabinofuranosidase | 2.344994194 |
| RBAM_RS16735 | ribokinase | 2.340987027 |
| RBAM_RS14530 | Na(+)/H(+) antiporter subunit B | 2.330674752 |
| RBAM_RS05520 | spore germination protein | 2.319456337 |
| RBAM_RS15345 | SMP-30/gluconolactonase/LRE family protein | 2.317723236 |
| RBAM_RS06875 | cell wall hydrolase | 2.30970335 |
| RBAM_RS17770 | glycosyltransferase family 2 protein | 2.304006702 |
| RBAM_RS18270 | citrate/sodium symporter CitN | 2.297894501 |
| RBAM_RS17830 | sucrose-6-phosphate hydrolase | 2.295116279 |
| RBAM_RS14555 | Na+/H+ antiporter subunit G | 2.288897517 |
| RBAM_RS18350 | 6-phospho-beta-glucosidase | 2.282878062 |
| RBAM_RS15125 | acyl-CoA dehydrogenase family protein | 2.271300609 |
| RBAM_RS16355 | phosphotransferase | 2.263566911 |
| RBAM_RS07855 | 16S rRNA (cytosine(967)-C(5))-methyltransferase Rsn | 2.260160539 |
| RBAM_RS00110 | cysteine hydrolase | 2.259389576 |
| RBAM_RS12880 | GerMN domain-containing protein | 2.246913904 |
| RBAM_RS00375 | stage II sporulation protein E | 2.246448294 |
| RBAM_RS18560 | CoA-acylating methylmalonate-semialdehyde dehydr | 2.244044644 |
| RBAM_RS10890 | RNA polymerase sporulation sigma factor SigF | 2.243362499 |
| RBAM_RS01845 | surfactin non-ribosomal peptide synthetase SrfAC | 2.237319239 |
| RBAM_RS10905 | D-alanyl-D-alanine carboxypeptidase | 2.23199235 |
| RBAM_RS12420 | amino acid ABC transporter permease | 2.229467113 |
| RBAM_RS16150 | carbonic anhydrase | 2.229020429 |
| RBAM_RS02550 | SpoIIE family protein phosphatase | 2.22848156 |
| RBAM_RS03035 | glycerol dehydrogenase | 2.222075893 |
| RBAM_RS06000 | subclass B1 metallo-beta-lactamase | 2.219561519 |
| RBAM_RS09600 | squalene--hopene cyclase | 2.217686003 |

| | | |
|---|---|---|
| RBAM_RS05550 | asparagine synthase (glutamine-hydrolyzing) | 2.217267936 |
| RBAM_RS12960 | TetR/AcrR family transcriptional regulator | 2.21622222 |
| RBAM_RS11370 | helix-turn-helix domain-containing protein | 2.216053686 |
| RBAM_RS07470 | hypothetical protein | 2.212768717 |
| RBAM_RS02260 | diguanylate cyclase | 2.212337301 |
| RBAM_RS02265 | DUF2334 domain-containing protein | 2.211006554 |
| RBAM_RS13650 | M42 family metallopeptidase | 2.209875803 |
| RBAM_RS11925 | diacylglycerol kinase family protein | 2.208541194 |
| RBAM_RS03535 | FMN-binding glutamate synthase family protein | 2.200116948 |
| RBAM_RS05495 | spore germination protein | 2.198194426 |
| RBAM_RS18145 | mannose-6-phosphate isomerase, class I | 2.197387808 |
| RBAM_RS05030 | ABC transporter ATP-binding protein | 2.196875899 |
| RBAM_RS07415 | glutaminase A | 2.18985244 |
| RBAM_RS03060 | GNAT family N-acetyltransferase | 2.186915734 |
| RBAM_RS08585 | polyketide synthase dehydratase domain-containing | 2.185485206 |
| RBAM_RS04705 | NAD(P)H-dependent oxidoreductase | 2.174402346 |
| RBAM_RS13565 | acetate--CoA ligase | 2.171464209 |
| RBAM_RS01455 | DoxX family protein | 2.169987104 |
| RBAM_RS03225 | tRNA (adenosine(37)-N6)-threonylcarbamoyltransfer | 2.167795879 |
| RBAM_RS16665 | glycosyltransferase | 2.162196488 |
| RBAM_RS01215 | aldo/keto reductase | 2.152346913 |
| RBAM_RS16405 | YitT family protein | 2.151275135 |
| RBAM_RS10025 | SCO family protein | 2.148259329 |
| RBAM_RS12405 | amino acid ABC transporter ATP-binding protein | 2.143411667 |
| RBAM_RS08580 | non-ribosomal peptide synthetase | 2.142911641 |
| RBAM_RS17775 | spore coat protein GerQ | 2.137831526 |
| RBAM_RS03855 | SDR family oxidoreductase | 2.135830592 |
| RBAM_RS03925 | NTTRR-F1 domain | 2.135636335 |
| RBAM_RS11110 | SDR family NAD(P)-dependent oxidoreductase | 2.131735966 |
| RBAM_RS00920 | KinB-signaling pathway activation protein | 2.127016215 |
| RBAM_RS18370 | glycoside hydrolase family 43 protein | 2.124489747 |
| RBAM_RS01555 | glucose 1-dehydrogenase | 2.122259725 |
| RBAM_RS08570 | SDR family NAD(P)-dependent oxidoreductase | 2.118982604 |
| RBAM_RS17380 | cardiolipin synthase | 2.118789322 |
| RBAM_RS06485 | LD-carboxypeptidase | 2.118715756 |
| RBAM_RS05545 | DUF2777 family protein | 2.117392659 |
| RBAM_RS09805 | hypothetical protein | 2.11720424 |
| RBAM_RS11620 | ComGF family competence protein | 2.117121022 |
| RBAM_RS06185 | MgtC/SapB family protein | 2.116100591 |
| RBAM_RS03820 | hypothetical protein | 2.111452638 |
| RBAM_RS09795 | phosphatase PAP2 family protein | 2.106814273 |
| RBAM_RS06490 | dipeptide epimerase | 2.102265127 |
| RBAM_RS07585 | stage V sporulation protein D | 2.101490624 |
| RBAM_RS17765 | CDP-glycerol glycerophosphotransferase family prote | 2.101107372 |
| RBAM_RS04815 | aspartate aminotransferase family protein | 2.099600258 |
| RBAM_RS11475 | stage III sporulation protein AF | 2.099449508 |
| RBAM_RS09145 | non-ribosomal peptide synthetase | 2.098135841 |

| | | |
|---|---|---|
| RBAM_RS04350 | MFS transporter | 2.096535662 |
| RBAM_RS02270 | glycosyltransferase family 2 protein | 2.086763603 |
| RBAM_RS15235 | serine protease | 2.084264529 |
| RBAM_RS19335 | hypothetical protein | 2.081396256 |
| RBAM_RS12300 | formate dehydrogenase subunit alpha | 2.080633604 |
| RBAM_RS15135 | enoyl-CoA hydratase/isomerase family protein | 2.078285273 |
| RBAM_RS09070 | SDR family oxidoreductase | 2.074366958 |
| RBAM_RS05510 | germination protein GerPC | 2.071118397 |
| RBAM_RS12955 | enoyl-CoA hydratase | 2.065135773 |
| RBAM_RS13030 | glycolate oxidase subunit GlcD | 2.064140409 |
| RBAM_RS16565 | MOP flippase family protein | 2.061932732 |
| RBAM_RS07085 | molybdenum cofactor guanylyltransferase | 2.05749746 |
| RBAM_RS16660 | Ger(x)C family spore germination protein | 2.04826776 |
| RBAM_RS07270 | SDR family NAD(P)-dependent oxidoreductase | 2.048176967 |
| RBAM_RS12985 | DNA polymerase/3'-5' exonuclease PolX | 2.043040686 |
| RBAM_RS05185 | hypothetical protein | 2.041735063 |
| RBAM_RS15730 | ABC transporter permease | 2.029645053 |
| RBAM_RS18695 | XdhC family protein | 2.019841283 |
| RBAM_RS03155 | APC family permease | 2.000185408 |
| RBAM_RS07785 | adenylyl-sulfate kinase | -2.000424869 |
| RBAM_RS06935 | methyl-accepting chemotaxis protein | -2.006884965 |
| RBAM_RS03845 | aldehyde dehydrogenase family protein | -2.007678777 |
| RBAM_RS03370 | hypothetical protein | -2.00804293 |
| RBAM_RS09775 | M15 family metallopeptidase | -2.013854576 |
| RBAM_RS16690 | C40 family peptidase | -2.014272907 |
| RBAM_RS03470 | phosphoribosylaminoimidazolesuccinocarboxamide s | -2.014913097 |
| RBAM_RS17435 | DUF1934 domain-containing protein | -2.016879247 |
| RBAM_RS15400 | iron-hydroxamate ABC transporter substrate-binding | -2.016975452 |
| RBAM_RS03460 | 5-(carboxyamino)imidazole ribonucleotide synthase | -2.018133899 |
| RBAM_RS14870 | homoserine dehydrogenase | -2.020833259 |
| RBAM_RS06180 | hypothetical protein | -2.023680376 |
| RBAM_RS02225 | GNAT family N-acetyltransferase | -2.026357023 |
| RBAM_RS06650 | hypothetical protein | -2.034398399 |
| RBAM_RS03290 | hypothetical protein | -2.036491541 |
| RBAM_RS12820 | 3-isopropylmalate dehydrogenase | -2.037799346 |
| RBAM_RS17590 | hypothetical protein | -2.048784918 |
| RBAM_RS09330 | excisionase family DNA-binding protein | -2.061591444 |
| RBAM_RS18025 | GTP pyrophosphokinase family protein | -2.062190258 |
| RBAM_RS08915 | recombinase family protein | -2.063813992 |
| RBAM_RS03905 | general stress protein | -2.064079725 |
| RBAM_RS17035 | P-II family nitrogen regulator | -2.073306898 |
| RBAM_RS16260 | ATP phosphoribosyltransferase regulatory subunit | -2.07557015 |
| RBAM_RS07770 | phosphoadenylyl-sulfate reductase | -2.076728829 |
| RBAM_RS16585 | hypothetical protein | -2.079269336 |
| RBAM_RS17350 | DNA-directed RNA polymerase subunit delta | -2.081049115 |
| RBAM_RS14775 | GMP reductase | -2.088270637 |
| RBAM_RS03255 | twin-arginine translocase TatA/TatE family subunit | -2.089160419 |

| | | |
|---|---|---|
| RBAM_RS10670 | genetic competence negative regulator | -2.093417561 |
| RBAM_RS19110 | serine/threonine protein kinase | -2.093529336 |
| RBAM_RS10675 | metallophosphoesterase | -2.096682567 |
| RBAM_RS16420 | ribosome-associated translation inhibitor RaiA | -2.101403364 |
| RBAM_RS18365 | (deoxy)nucleoside triphosphate pyrophosphohydrola | -2.10150573 |
| RBAM_RS06970 | chemotaxis protein CheV | -2.102515156 |
| RBAM_RS13525 | type 1 glutamine amidotransferase | -2.103247518 |
| RBAM_RS17595 | hypothetical protein | -2.111116784 |
| RBAM_RS13925 | glycerophosphoryl diester phosphodiesterase | -2.112362445 |
| RBAM_RS12310 | hypothetical protein | -2.113139143 |
| RBAM_RS16385 | right-handed parallel beta-helix repeat-containing pr | -2.11821803 |
| RBAM_RS11810 | DUF308 domain-containing protein | -2.119589012 |
| RBAM_RS10160 | hypothetical protein | -2.123324772 |
| RBAM_RS15775 | PIG-L family deacetylase | -2.125131292 |
| RBAM_RS16820 | gamma-glutamyltransferase | -2.133149886 |
| RBAM_RS05335 | histidine phosphatase family protein | -2.133997942 |
| RBAM_RS03440 | Fur-regulated basic protein FbpA | -2.139054102 |
| RBAM_RS08115 | MotE family protein | -2.146401305 |
| RBAM_RS02085 | NAD-dependent succinate-semialdehyde dehydroger | -2.146514218 |
| RBAM_RS11175 | NADPH dehydrogenase NamA | -2.156079911 |
| RBAM_RS17875 | GNAT family N-acetyltransferase | -2.15908081 |
| RBAM_RS04690 | MarR family transcriptional regulator | -2.162319646 |
| RBAM_RS04455 | peroxide-responsive transcriptional repressor PerR | -2.162470107 |
| RBAM_RS14940 | PucR family transcriptional regulator | -2.17243832 |
| RBAM_RS05180 | YtxH domain-containing protein | -2.178410336 |
| RBAM_RS00310 | aminoacyl-tRNA hydrolase | -2.180269948 |
| RBAM_RS18060 | branched-chain amino acid aminotransferase | -2.185025294 |
| RBAM_RS16305 | TetR/AcrR family transcriptional regulator | -2.190652431 |
| RBAM_RS05615 | hypothetical protein | -2.191667117 |
| RBAM_RS13475 | hypothetical protein | -2.199126351 |
| RBAM_RS17420 | Crp/Fnr family transcriptional regulator | -2.199491155 |
| RBAM_RS01980 | M20 peptidase aminoacylase family protein | -2.205301201 |
| RBAM_RS13105 | DUF1294 domain-containing protein | -2.207585656 |
| RBAM_RS12825 | 2-isopropylmalate synthase | -2.208959498 |
| RBAM_RS13250 | YjdF family protein | -2.209429076 |
| RBAM_RS02910 | MerR family transcriptional regulator | -2.212838087 |
| RBAM_RS05585 | bifunctional homocysteine S-methyltransferase/metl | -2.213573913 |
| RBAM_RS07045 | flavodoxin | -2.218654306 |
| RBAM_RS09470 | hypothetical protein | -2.221703625 |
| RBAM_RS07395 | YlaI family protein | -2.223131166 |
| RBAM_RS03450 | NETI motif-containing protein | -2.223643414 |
| RBAM_RS06595 | 5-methyltetrahydropteroyltriglutamate--homocystein | -2.230588702 |
| RBAM_RS02690 | SDR family NAD(P)-dependent oxidoreductase | -2.231726119 |
| RBAM_RS10570 | trp RNA-binding attenuation protein MtrB | -2.236764332 |
| RBAM_RS13140 | sensor histidine kinase | -2.245346777 |
| RBAM_RS09395 | replication termination protein | -2.249511231 |
| RBAM_RS16225 | bifunctional phosphoribosyl-AMP cyclohydrolase/pho | -2.2568887 |

| | | |
|---|---|---|
| RBAM_RS18900 | DUF2188 domain-containing protein | -2.26380344 |
| RBAM_RS19325 | hypothetical protein | -2.263838148 |
| RBAM_RS05405 | YbjQ family protein | -2.267902773 |
| RBAM_RS15435 | molybdate ABC transporter permease subunit | -2.268082309 |
| RBAM_RS06550 | organic hydroperoxide resistance protein | -2.271855395 |
| RBAM_RS12815 | 3-isopropylmalate dehydratase large subunit | -2.272693739 |
| RBAM_RS18950 | 50S ribosomal protein L9 | -2.273872025 |
| RBAM_RS05840 | competence protein CoiA | -2.276727392 |
| RBAM_RS16230 | imidazole glycerol phosphate synthase subunit HisF | -2.280720429 |
| RBAM_RS14830 | NupC/NupG family nucleoside CNT transporter | -2.282702406 |
| RBAM_RS09445 | 2-hydroxyacid dehydrogenase | -2.282917712 |
| RBAM_RS14325 | methyl-accepting chemotaxis protein | -2.28503574 |
| RBAM_RS01955 | LLM class flavin-dependent oxidoreductase | -2.285381111 |
| RBAM_RS15800 | DegT/DnrJ/EryC1/StrS family aminotransferase | -2.289762923 |
| RBAM_RS02720 | cold shock protein CspC | -2.293404527 |
| RBAM_RS15760 | GNAT family N-acetyltransferase | -2.294806595 |
| RBAM_RS17660 | bacilysin biosynthesis protein BacA | -2.309338351 |
| RBAM_RS02960 | TetR/AcrR family transcriptional regulator | -2.314911464 |
| RBAM_RS00115 | nucleoside deaminase | -2.31684429 |
| RBAM_RS15715 | GbsR/MarR family transcriptional regulator | -2.318316693 |
| RBAM_RS08720 | type I glutamate--ammonia ligase | -2.319312532 |
| RBAM_RS15145 | proline dehydrogenase family protein | -2.32099308 |
| RBAM_RS11740 | endolytic transglycosylase MltG | -2.323146179 |
| RBAM_RS14320 | methyl-accepting chemotaxis protein | -2.326314167 |
| RBAM_RS13725 | DeoR/GlpR transcriptional regulator | -2.329853929 |
| RBAM_RS08785 | hypothetical protein | -2.332432001 |
| RBAM_RS01280 | hypothetical protein | -2.337946038 |
| RBAM_RS17825 | hypothetical protein | -2.339819653 |
| RBAM_RS18005 | VOC family protein | -2.346812474 |
| RBAM_RS08025 | FlhB-like flagellar biosynthesis protein | -2.349017425 |
| RBAM_RS16235 | 1-(5-phosphoribosyl)-5-(5-phosphoribosylamino)met | -2.352952303 |
| RBAM_RS14760 | YuiA family protein | -2.361104225 |
| RBAM_RS15770 | endonuclease | -2.364205797 |
| RBAM_RS05455 | TetR/AcrR family transcriptional regulator | -2.366917531 |
| RBAM_RS10005 | peptide-methionine (S)-S-oxide reductase MsrA | -2.371686846 |
| RBAM_RS16465 | flagellar hook-associated protein FlgL | -2.377768785 |
| RBAM_RS01645 | hypothetical protein | -2.380756409 |
| RBAM_RS16990 | flagellar hook-basal body protein | -2.386212529 |
| RBAM_RS15360 | helix-turn-helix domain-containing protein | -2.392034888 |
| RBAM_RS16245 | imidazoleglycerol-phosphate dehydratase HisB | -2.404457485 |
| RBAM_RS16435 | flagellar export chaperone FliS | -2.407983367 |
| RBAM_RS03710 | YezD family protein | -2.408965046 |
| RBAM_RS06655 | Fur-regulated basic protein FbpA | -2.426240958 |
| RBAM_RS00130 | YbaB/EbfC family nucleoid-associated protein | -2.432270578 |
| RBAM_RS09325 | site-specific integrase | -2.442202679 |
| RBAM_RS01965 | amino acid ABC transporter substrate-binding protei | -2.453215455 |
| RBAM_RS03875 | MFS transporter | -2.45482497 |

| | | |
|---|---|---|
| RBAM_RS16090 | hypothetical protein | -2.455245297 |
| RBAM_RS08790 | hypothetical protein | -2.460201974 |
| RBAM_RS02385 | DUF4937 domain-containing protein | -2.469808856 |
| RBAM_RS17095 | urease subunit alpha | -2.478077336 |
| RBAM_RS10495 | histidinol-phosphate transaminase | -2.48241405 |
| RBAM_RS01855 | aminotransferase class I/II-fold pyridoxal phosphate-( | -2.482473307 |
| RBAM_RS05145 | peptidylprolyl isomerase | -2.48362256 |
| RBAM_RS02145 | HAD family phosphatase | -2.490086776 |
| RBAM_RS11180 | alpha/beta fold hydrolase | -2.499667056 |
| RBAM_RS14820 | iron-sulfur cluster assembly accessory protein | -2.501316343 |
| RBAM_RS08015 | ribonuclease HII | -2.504730235 |
| RBAM_RS09930 | hypothetical protein | -2.506640111 |
| RBAM_RS01815 | DNA-entry nuclease | -2.507431881 |
| RBAM_RS06810 | ATP-dependent Clp protease ATP-binding subunit | -2.510270068 |
| RBAM_RS04450 | thioredoxin-dependent thiol peroxidase | -2.516552091 |
| RBAM_RS16075 | lipase pseudo=true partial=3' | -2.526741593 |
| RBAM_RS02625 | Lrp/AsnC family transcriptional regulator | -2.528468123 |
| RBAM_RS10010 | MarR family transcriptional regulator | -2.533054038 |
| RBAM_RS15375 | oxalate decarboxylase family bicupin | -2.542220834 |
| RBAM_RS15795 | NAD(P)-dependent oxidoreductase | -2.543047444 |
| RBAM_RS10500 | tryptophan synthase subunit alpha | -2.543418192 |
| RBAM_RS06800 | flagellar motor protein MotB | -2.547745464 |
| RBAM_RS08620 | poly-gamma-glutamate hydrolase family protein | -2.551221732 |
| RBAM_RS10240 | hypothetical protein | -2.554843011 |
| RBAM_RS03655 | YebC/PmpR family DNA-binding transcriptional regul: | -2.55951771 |
| RBAM_RS16870 | hypothetical protein | -2.582012477 |
| RBAM_RS16070 | DUF1433 domain-containing protein | -2.588275236 |
| RBAM_RS07380 | YlaF family protein | -2.596618581 |
| RBAM_RS15780 | WbqC family protein | -2.613832893 |
| RBAM_RS14365 | zinc metallopeptidase | -2.615522609 |
| RBAM_RS08625 | OsmC family protein | -2.618371683 |
| RBAM_RS12810 | 3-isopropylmalate dehydratase small subunit | -2.624777361 |
| RBAM_RS17480 | hypothetical protein | -2.627024395 |
| RBAM_RS16470 | flagellar hook-associated protein FlgK | -2.63178225 |
| RBAM_RS05370 | IDEAL domain-containing protein | -2.633383082 |
| RBAM_RS01515 | triacylglycerol lipase | -2.640829382 |
| RBAM_RS04625 | 30S ribosomal protein S14 | -2.64309885 |
| RBAM_RS18335 | pectate lyase | -2.648844025 |
| RBAM_RS17370 | hypothetical protein | -2.658904516 |
| RBAM_RS02135 | beta-hydroxyacyl-ACP dehydratase | -2.660101535 |
| RBAM_RS02350 | hypothetical protein | -2.660306667 |
| RBAM_RS08940 | aspartyl-phosphate phosphatase Spo0E family protei | -2.666578496 |
| RBAM_RS18630 | helix-turn-helix transcriptional regulator | -2.674429082 |
| RBAM_RS10195 | MGMT family protein | -2.687730096 |
| RBAM_RS12830 | ketol-acid reductoisomerase | -2.695232655 |
| RBAM_RS14425 | general stress protein 13 | -2.695646485 |
| RBAM_RS10215 | hypothetical protein | -2.703827627 |

| | | |
|---|---|---|
| RBAM_RS09555 | hypothetical protein pseudo=true partial=3' | -2.705260234 |
| RBAM_RS08080 | flagellar basal body rod protein FlgC | -2.711529755 |
| RBAM_RS16815 | hypothetical protein pseudo=true partial=3' | -2.713401156 |
| RBAM_RS03300 | hypothetical protein | -2.720100688 |
| RBAM_RS03390 | GABA permease | -2.736982446 |
| RBAM_RS12600 | SafA/ExsA family spore coat assembly protein | -2.739066143 |
| RBAM_RS03365 | 2,3-butanediol dehydrogenase | -2.740822449 |
| RBAM_RS09725 | alpha/beta hydrolase | -2.743643608 |
| RBAM_RS02610 | NAD(P)H-dependent oxidoreductase | -2.75026459 |
| RBAM_RS15805 | nucleotide sugar dehydrogenase | -2.751197571 |
| RBAM_RS14275 | NO-inducible flavohemoprotein | -2.751720246 |
| RBAM_RS09025 | CoA-binding protein | -2.762753034 |
| RBAM_RS18035 | teichoic acid D-Ala incorporation-associated protein I | -2.791725474 |
| RBAM_RS13450 | hypothetical protein | -2.799011279 |
| RBAM_RS13865 | GntR family transcriptional regulator | -2.800662984 |
| RBAM_RS18340 | DMT family transporter | -2.806696772 |
| RBAM_RS08665 | hypothetical protein | -2.841885334 |
| RBAM_RS15550 | MFS transporter | -2.844251724 |
| RBAM_RS11200 | NADP-dependent phosphogluconate dehydrogenase | -2.860451545 |
| RBAM_RS11235 | stressosome-associated protein Prli42 | -2.877771292 |
| RBAM_RS02140 | hypothetical protein | -2.905196194 |
| RBAM_RS03660 | hypothetical protein | -2.919548863 |
| RBAM_RS06805 | flagellar motor stator protein MotA | -2.920901525 |
| RBAM_RS09375 | glutamate synthase large subunit | -2.921547724 |
| RBAM_RS14690 | MbtH family NRPS accessory protein | -2.934973853 |
| RBAM_RS18910 | YycC family protein | -2.936903677 |
| RBAM_RS05785 | tryptophan--tRNA ligase | -2.952539019 |
| RBAM_RS02525 | anti-sigma regulatory factor | -2.968064969 |
| RBAM_RS16640 | glucosaminidase domain-containing protein | -2.97136354 |
| RBAM_RS00420 | cysteine synthase A | -2.972125071 |
| RBAM_RS01775 | FAD-dependent oxidoreductase | -2.978351447 |
| RBAM_RS04595 | catalase | -2.987522452 |
| RBAM_RS01765 | NADPH-nitrite reductase | -2.988332866 |
| RBAM_RS16065 | DUF1433 domain-containing protein | -3.037564611 |
| RBAM_RS02680 | NAD(P)-dependent oxidoreductase | -3.042550665 |
| RBAM_RS16475 | flagellar protein FlgN | -3.04490918 |
| RBAM_RS16440 | flagellar hook-associated protein 2 | -3.04532162 |
| RBAM_RS02440 | Fur-regulated basic protein FbpB | -3.054314333 |
| RBAM_RS15165 | DHA2 family efflux MFS transporter permease subun | -3.066570695 |
| RBAM_RS11360 | hypothetical protein | -3.085635298 |
| RBAM_RS01985 | MmgE/PrpD family protein | -3.087296844 |
| RBAM_RS06620 | ECF transporter S component | -3.090344116 |
| RBAM_RS06220 | cytochrome P450 | -3.091804417 |
| RBAM_RS13365 | thiol peroxidase | -3.098682809 |
| RBAM_RS07660 | acetylornithine deacetylase | -3.103449771 |
| RBAM_RS04740 | cold-shock protein | -3.104658703 |
| RBAM_RS14235 | DinB family protein | -3.105112419 |

| | | |
|---|---|---|
| RBAM_RS05445 | AbrB family transcriptional regulator | -3.120810273 |
| RBAM_RS04635 | helix-turn-helix transcriptional regulator | -3.123705302 |
| RBAM_RS11580 | DNA-binding anti-repressor SinI | -3.128748863 |
| RBAM_RS14945 | hypothetical protein | -3.13059542 |
| RBAM_RS17980 | CidB/LrgB family autolysis modulator | -3.134940185 |
| RBAM_RS03455 | 5-(carboxyamino)imidazole ribonucleotide mutase | -3.168489178 |
| RBAM_RS08900 | hypothetical protein | -3.171219013 |
| RBAM_RS15160 | MarR family transcriptional regulator | -3.19712392 |
| RBAM_RS05420 | hypothetical protein | -3.20032691 |
| RBAM_RS07665 | YlmC/YmxH family sporulation protein | -3.205991864 |
| RBAM_RS08605 | hypothetical protein | -3.21622844 |
| RBAM_RS10000 | peptide-methionine (R)-S-oxide reductase MsrB | -3.221737715 |
| RBAM_RS07400 | YhcN/YlaJ family sporulation lipoprotein | -3.227760178 |
| RBAM_RS12195 | AAA family ATPase | -3.229366921 |
| RBAM_RS18195 | 2-hydroxycarboxylate transporter family protein | -3.235483724 |
| RBAM_RS14615 | YueH family protein | -3.245884812 |
| RBAM_RS09580 | PH domain-containing protein | -3.263859211 |
| RBAM_RS06795 | MarR family transcriptional regulator | -3.271425206 |
| RBAM_RS03980 | YflJ family protein | -3.277622405 |
| RBAM_RS17855 | LLM class flavin-dependent oxidoreductase | -3.280370914 |
| RBAM_RS19245 | 50S ribosomal protein L34 | -3.281678245 |
| RBAM_RS09370 | glutamate synthase small subunit | -3.307734418 |
| RBAM_RS14030 | type B 50S ribosomal protein L31 | -3.315548808 |
| RBAM_RS17485 | stage II sporulation protein M | -3.316030202 |
| RBAM_RS05925 | thiazole biosynthesis adenylyltransferase ThiF | -3.319460974 |
| RBAM_RS12625 | transcription repressor NadR | -3.320378933 |
| RBAM_RS02455 | thioredoxin family protein | -3.329056522 |
| RBAM_RS16195 | thioredoxin-disulfide reductase | -3.329391433 |
| RBAM_RS10250 | hypothetical protein | -3.334021826 |
| RBAM_RS04575 | phosphomethylpyrimidine synthase ThiC | -3.337609725 |
| RBAM_RS15430 | molybdate ABC transporter substrate-binding proteir | -3.366989875 |
| RBAM_RS13965 | DUF2584 domain-containing protein | -3.373816038 |
| RBAM_RS19180 | DUF951 domain-containing protein | -3.38009301 |
| RBAM_RS02025 | aspartate kinase | -3.381048166 |
| RBAM_RS13905 | MFS transporter | -3.381303436 |
| RBAM_RS02685 | TetR/AcrR family transcriptional regulator | -3.383017272 |
| RBAM_RS16775 | acetolactate synthase AlsS | -3.398469881 |
| RBAM_RS05355 | globin-coupled sensor protein | -3.398716288 |
| RBAM_RS09365 | hypothetical protein | -3.409053519 |
| RBAM_RS14330 | methyl-accepting chemotaxis protein | -3.40937733 |
| RBAM_RS09210 | 6-carboxyhexanoate--CoA ligase | -3.413780414 |
| RBAM_RS18970 | spore coat protein | -3.445202664 |
| RBAM_RS05920 | thiazole synthase | -3.44661647 |
| RBAM_RS16485 | membrane protein | -3.461952411 |
| RBAM_RS08920 | DUF896 domain-containing protein | -3.463705549 |
| RBAM_RS14575 | competence pheromone ComX | -3.487811159 |
| RBAM_RS17100 | urease subunit beta | -3.508326719 |

| | | |
|---|---|---|
| RBAM_RS01510 | type II asparaginase | -3.516249261 |
| RBAM_RS02895 | HAD-IIA family hydrolase | -3.529380713 |
| RBAM_RS00315 | anti-sigma-F factor Fin family protein | -3.532027827 |
| RBAM_RS01770 | molybdopterin-dependent oxidoreductase | -3.536341803 |
| RBAM_RS17490 | hypothetical protein | -3.54802892 |
| RBAM_RS17865 | oxygen-insensitive NADPH nitroreductase | -3.573014807 |
| RBAM_RS13480 | hypothetical protein | -3.577688149 |
| RBAM_RS06645 | MerR family transcriptional regulator TnrA | -3.610829701 |
| RBAM_RS06225 | UDP-glucosyltransferase | -3.626648003 |
| RBAM_RS15080 | MetQ/NlpA family ABC transporter substrate-binding | -3.628838882 |
| RBAM_RS04145 | hypothetical protein | -3.652683156 |
| RBAM_RS08660 | hypothetical protein | -3.665098995 |
| RBAM_RS15530 | copper-sensing transcriptional repressor CsoR | -3.671464187 |
| RBAM_RS11730 | 50S ribosomal protein L33 | -3.682410775 |
| RBAM_RS12295 | DUF2294 domain-containing protein | -3.689512507 |
| RBAM_RS17975 | CidA/LrgA family holin-like protein | -3.689802033 |
| RBAM_RS10210 | cysteine hydrolase | -3.716098774 |
| RBAM_RS16770 | acetolactate decarboxylase | -3.717431575 |
| RBAM_RS09440 | sugar kinase | -3.718648061 |
| RBAM_RS05910 | glycine oxidase ThiO | -3.723087543 |
| RBAM_RS05285 | YhfH family protein | -3.737952862 |
| RBAM_RS10925 | YqzK family protein | -3.818363266 |
| RBAM_RS05400 | excalibur calcium-binding domain-containing protein | -3.827481912 |
| RBAM_RS17105 | urease subunit gamma | -3.860032597 |
| RBAM_RS12555 | preprotein translocase subunit YajC | -3.895156369 |
| RBAM_RS10520 | anthranilate phosphoribosyltransferase | -3.902027165 |
| RBAM_RS10525 | anthranilate synthase component I | -3.908965423 |
| RBAM_RS10995 | YqkE family protein | -3.935780343 |
| RBAM_RS10515 | indole-3-glycerol phosphate synthase TrpC | -3.944617738 |
| RBAM_RS17600 | hypothetical protein | -3.946432489 |
| RBAM_RS13785 | rhodanese-like domain-containing protein | -3.972059413 |
| RBAM_RS02890 | bifunctional hydroxymethylpyrimidine kinase/phosph | -3.98880651 |
| RBAM_RS04370 | YfhJ family protein | -4.018075136 |
| RBAM_RS02725 | CarD family transcriptional regulator | -4.04208481 |
| RBAM_RS03295 | hypothetical protein | -4.085681601 |
| RBAM_RS17475 | DUF4177 domain-containing protein | -4.119610098 |
| RBAM_RS07375 | hypothetical protein | -4.123298936 |
| RBAM_RS17465 | helix-turn-helix transcriptional regulator | -4.126301477 |
| RBAM_RS17140 | VWA domain-containing protein | -4.130666919 |
| RBAM_RS17615 | putative sulfate exporter family transporter | -4.13844034 |
| RBAM_RS09590 | hypothetical protein | -4.141561942 |
| RBAM_RS07185 | AbrB/MazE/SpoVT family DNA-binding domain-conta | -4.141933233 |
| RBAM_RS17495 | LuxR family transcriptional regulator | -4.165878493 |
| RBAM_RS19290 | hypothetical protein | -4.192922812 |
| RBAM_RS05930 | bifunctional hydroxymethylpyrimidine kinase/phosph | -4.212468682 |
| RBAM_RS10510 | phosphoribosylanthranilate isomerase | -4.214473018 |
| RBAM_RS13190 | winged helix-turn-helix transcriptional regulator | -4.225555514 |

| | | |
|---|---|---|
| RBAM_RS03870 | MerR family transcriptional regulator | -4.237697173 |
| RBAM_RS01615 | MFS transporter | -4.270965814 |
| RBAM_RS05905 | thiazole tautomerase TenI | -4.273158141 |
| RBAM_RS10825 | hypothetical protein | -4.299799203 |
| RBAM_RS03695 | heme-degrading oxygenase HmoA | -4.304175901 |
| RBAM_RS16445 | flagellin Hag | -4.335474551 |
| RBAM_RS03770 | plantazolicin family RiPP | -4.336520696 |
| RBAM_RS01780 | NarK/NasA family nitrate transporter | -4.400762596 |
| RBAM_RS11185 | 50S ribosomal protein L33 | -4.418055702 |
| RBAM_RS10505 | tryptophan synthase subunit beta | -4.430455073 |
| RBAM_RS05045 | YheE family protein | -4.434651507 |
| RBAM_RS08890 | prohibitin family protein | -4.462795278 |
| RBAM_RS16480 | flagellar biosynthesis anti-sigma factor FlgM | -4.654511014 |
| RBAM_RS02880 | thymidylate synthase | -4.66915063 |
| RBAM_RS14000 | DNA starvation/stationary phase protection protein | -4.677242186 |
| RBAM_RS02445 | Fur-regulated basic protein FbpA | -4.697861938 |
| RBAM_RS09315 | hypothetical protein | -4.782325712 |
| RBAM_RS02885 | class I SAM-dependent methyltransferase | -4.833233747 |
| RBAM_RS06115 | antibiotic biosynthesis monooxygenase | -4.92606984 |
| RBAM_RS12620 | IscS subfamily cysteine desulfurase | -4.975911257 |
| RBAM_RS00270 | small, acid-soluble spore protein, alpha/beta type | -5.009121061 |
| RBAM_RS15230 | DNA starvation/stationary phase protection protein | -5.082529015 |
| RBAM_RS17470 | hypothetical protein | -5.155474491 |
| RBAM_RS17030 | ammonium transporter | -5.39556093 |
| RBAM_RS12605 | quinolinate synthase NadA | -5.416909757 |
| RBAM_RS02730 | metal-sensitive transcriptional regulator | -5.453928861 |
| RBAM_RS16080 | DUF1433 domain-containing protein | -5.681322608 |
| RBAM_RS02875 | hypothetical protein | -5.819786124 |
| RBAM_RS06050 | hypothetical protein | -5.839911304 |
| RBAM_RS09585 | DUF4025 domain-containing protein | -6.042009666 |
| RBAM_RS15525 | copper chaperone CopZ | -6.126077974 |
| RBAM_RS05900 | thiaminase II | -6.179871327 |
| RBAM_RS12610 | carboxylating nicotinate-nucleotide diphosphorylase | -6.186891603 |
| RBAM_RS13495 | hypothetical protein | -6.487883343 |
| RBAM_RS13130 | antiholin-like murein hydrolase modulator LrgA | -6.523580495 |
| RBAM_RS10280 | hypothetical protein | -6.535315696 |
| RBAM_RS06060 | hypothetical protein | -6.549031197 |
| RBAM_RS09655 | glycosyl transferase family 1 | -6.5532805 |
| RBAM_RS04835 | glycerol-3-phosphate dehydrogenase/oxidase | -6.747366569 |
| RBAM_RS13490 | hypothetical protein pseudo=true | -6.757351164 |
| RBAM_RS17385 | YitT family protein | -6.761845697 |
| RBAM_RS19300 | hypothetical protein | -6.900702812 |
| RBAM_RS12615 | L-aspartate oxidase | -6.910487267 |
| RBAM_RS14585 | hypothetical protein | -7.097043125 |
| RBAM_RS13125 | antiholin-like protein LrgB | -7.152555539 |
| RBAM_RS10940 | hypothetical protein | -7.368556093 |
| RBAM_RS01430 | tryptophan RNA-binding attenuator protein inhibitor | -8.224998891 |

| | | |
|---|---|---|
| RBAM_RS10110 | hypothetical protein pseudo=true | -8.286613986 |
| RBAM_RS05915 | sulfur carrier protein ThiS | -8.618959389 |
| RBAM_RS04590 | ABC transporter substrate-binding protein pseudo=tr | -10.03884707 |
