## Supplementary Table 1 for "Chemical interplay and complementary adaptative strategies toggle bacterial antagonism and co-existence"

| Gene | Product | log2FoldCha | pvalue |
|---|---|---|---|
| PCL1606_RS0! | hypothetical p | 3.10 | 8.5E-09 |
| PCL1606_RS1( | hypothetical p | 2.96 | 3.0E-08 |
| PCL1606_RS2. | N-acetyltransf | 2.64 | 1.5E-06 |
| PCL1606_RS1: | MexE family m | 2.63 | 4.8E-07 |
| PCL1606_RS2. | AraC family tra | 2.61 | 3.3E-07 |
| PCL1606_RS1( | peptide chain | 2.31 | 2.2E-06 |
| PCL1606_RS1: | multidrug efflu | 2.31 | 5.8E-06 |
| PCL1606_RS2. | pyridine nucle | 2.12 | 8.5E-07 |
| PCL1606_RS1{ | hypothetical p | 2.02 | 7.0E-08 |
| PCL1606_RS0{ | YIP1 family pro | 1.96 | 3.1E-05 |
| PCL1606_RS1: | MFS transport | 1.92 | 5.1E-08 |
| PCL1606_RS1( | hypothetical p | 1.89 | 1.6E-04 |
| PCL1606_RS1( | RtcB family pr | 1.84 | 7.0E-05 |
| PCL1606_RS2( | MATE family e | 1.83 | 1.8E-04 |
| PCL1606_RS1( | general secret | 1.81 | 1.0E-03 |
| PCL1606_RS0! | hypothetical p | 1.77 | 7.7E-04 |
| PCL1606_RS1( | acyl-CoA synth | 1.73 | 1.7E-03 |
| PCL1606_RS1( | general secret | 1.69 | 2.2E-03 |
| PCL1606_RS1( | hypothetical p | 1.65 | 1.4E-03 |
| PCL1606_RS1. | acyl-CoA dehy | 1.65 | 2.8E-03 |
| PCL1606_RS1. | molybdate AB | 1.63 | 1.6E-04 |
| PCL1606_RS1. | ABC transport | 1.63 | 3.1E-03 |
| PCL1606_RS1. | peptide ABC tr | 1.62 | 3.3E-03 |
| PCL1606_RS1. | peptidase T4 | 1.60 | 3.6E-03 |
| PCL1606_RS1. | acyl-CoA synth | 1.59 | 4.0E-03 |
| PCL1606_RS1. | ABC transport | 1.59 | 4.2E-03 |
| PCL1606_RS1( | aspartate ami | 1.57 | 4.0E-03 |
| PCL1606_RS1. | ABC transport | 1.56 | 4.3E-03 |
| PCL1606_RS1( | type II secretio | 1.55 | 5.1E-03 |
| PCL1606_RS2! | hypothetical p | 1.55 | 4.4E-03 |
| PCL1606_RS1: | glycerate kina | 1.54 | 1.6E-05 |
| PCL1606_RS1( | decarboxylase | 1.53 | 4.7E-03 |
| PCL1606_RS1. | acyl-CoA dehy | 1.53 | 5.6E-03 |
| PCL1606_RS0. | hypothetical p | 1.52 | 5.1E-03 |
| PCL1606_RS1( | aldehyde dehy | 1.51 | 6.0E-03 |
| PCL1606_RS2. | 6-O-methylgu | 1.50 | 3.1E-03 |
| PCL1606_RS1( | pseudopilin | 1.50 | 5.9E-03 |
| PCL1606_RS1{ | hypothetical p | 1.49 | 7.4E-05 |
| PCL1606_RS1. | stationary-pha | 1.49 | 7.1E-03 |
| PCL1606_RS1. | hypothetical p | 1.49 | 6.9E-03 |
| PCL1606_RS1. | ABC transport | 1.48 | 7.2E-03 |
| PCL1606_RS1! | hemagglutinin | 1.48 | 5.8E-03 |
| PCL1606_RS1( | general secret | 1.48 | 7.2E-03 |
| PCL1606_RS1( | acyl-CoA dehy | 1.47 | 5.9E-03 |
| PCL1606_RS1( | type II secretio | 1.47 | 5.8E-03 |
| PCL1606_RS1. | hypothetical p | 1.47 | 6.8E-03 |
| PCL1606_RS1( | hypothetical p | 1.47 | 1.7E-10 |

| | | | |
|---|---|---|---|
| PCL1606_RS14 | nitrilotriacetat | 1.47 | 7.9E-03 |
| PCL1606_RS2 | D-amino-acid | 1.46 | 3.4E-03 |
| PCL1606_RS14 | ABC transport | 1.44 | 7.1E-03 |
| PCL1606_RS1 | thiol:disulfide | 1.44 | 5.9E-03 |
| PCL1606_RS1 | enoyl-CoA hyc | 1.44 | 8.1E-03 |
| PCL1606_RS1 | hypothetical p | 1.44 | 8.7E-03 |
| PCL1606_RS0 | peptidase M4 | 1.44 | 7.1E-03 |
| PCL1606_RS14 | acyl-CoA dehy | 1.43 | 9.5E-03 |
| PCL1606_RS1 | type II secretic | 1.43 | 7.5E-03 |
| PCL1606_RS2 | glycine/betain | 1.43 | 5.8E-03 |
| PCL1606_RS2 | (2Fe-2S)-bindi | 1.42 | 1.0E-02 |
| PCL1606_RS1 | AP endonucle | 1.41 | 6.6E-03 |
| PCL1606_RS1 | type II secretic | 1.41 | 9.2E-03 |
| PCL1606_RS14 | LLM class flavi | 1.41 | 1.1E-02 |
| PCL1606_RS0 | hypothetical p | 1.40 | 3.6E-03 |
| PCL1606_RS1 | butanediol de | 1.40 | 1.1E-02 |
| PCL1606_RS1 | spermidine/pu | 1.40 | 1.1E-02 |
| PCL1606_RS0 | aldehyde dehy | 1.40 | 2.4E-03 |
| PCL1606_RS0 | membrane pro | 1.39 | 1.1E-02 |
| PCL1606_RS1 | amino acid AB | 1.39 | 8.2E-03 |
| PCL1606_RS2 | TetR family tra | 1.38 | 4.4E-03 |
| PCL1606_RS1 | ketosteroid iso | 1.38 | 2.7E-03 |
| PCL1606_RS04 | ribosome alte | 1.37 | 2.9E-05 |
| PCL1606_RS14 | diaminopimela | 1.36 | 4.7E-03 |
| PCL1606_RS2 | hydroxyprolin | 1.36 | 8.1E-03 |
| PCL1606_RS0 | methyltransfe | 1.35 | 1.3E-02 |
| PCL1606_RS14 | tetratricopept | 1.34 | 1.1E-02 |
| PCL1606_RS1 | porin | 1.33 | 1.2E-02 |
| PCL1606_RS04 | TetR family tra | 1.33 | 4.6E-03 |
| PCL1606_RS14 | iron ABC trans | 1.33 | 1.1E-02 |
| PCL1606_RS0 | triphosphoribo | 1.33 | 1.6E-02 |
| PCL1606_RS1 | 2,4-dihydroxy | 1.31 | 1.1E-03 |
| PCL1606_RS14 | AcsD protein | 1.31 | 1.8E-04 |
| PCL1606_RS0 | hypothetical p | 1.31 | 1.6E-02 |
| PCL1606_RS2 | iron dicitrate t | 1.31 | 1.7E-07 |
| PCL1606_RS2 | DUF1328 dom | 1.30 | 2.0E-04 |
| PCL1606_RS1 | porin | 1.30 | 3.5E-03 |
| PCL1606_RS0 | transcriptiona | 1.30 | 1.9E-02 |
| PCL1606_RS1 | membrane pro | 1.30 | 1.9E-02 |
| PCL1606_RS14 | AcsA protein | 1.29 | 3.2E-03 |
| PCL1606_RS0 | CopG family tr | 1.29 | 2.0E-02 |
| PCL1606_RS1 | muconate cyc | 1.29 | 4.2E-03 |
| PCL1606_RS14 | peptide ABC tr | 1.28 | 1.4E-02 |
| PCL1606_RS1 | Na/Pi cotransp | 1.28 | 1.2E-02 |
| PCL1606_RS1 | membrane pro | 1.28 | 1.8E-11 |
| PCL1606_RS2 | lipoprotein | 1.28 | 1.5E-02 |
| PCL1606_RS1 | LysR family tra | 1.28 | 2.0E-02 |

| | | | |
|---|---|---|---|
| PCL1606_RS1( | oxidoreductas | 1.27 | 1.5E-02 |
| PCL1606_RS0: | phenylalanine | 1.27 | 1.9E-02 |
| PCL1606_RS1: | branched-chai | 1.27 | 1.7E-02 |
| PCL1606_RS0: | peptidase M1! | 1.27 | 1.2E-02 |
| PCL1606_RS0: | conjugal trans | 1.27 | 2.2E-02 |
| PCL1606_RS0: | P-type conjug: | 1.27 | 2.1E-02 |
| PCL1606_RS0! | general secret | 1.27 | 1.6E-02 |
| PCL1606_RS2: | MFS transport | 1.26 | 2.1E-09 |
| PCL1606_RS0: | biotin-indeper | 1.26 | 2.2E-02 |
| PCL1606_RS1: | MFS transport | 1.26 | 2.2E-02 |
| PCL1606_RS0: | hypothetical p | 1.26 | 7.4E-03 |
| PCL1606_RS0: | (2Fe-2S)-bindi | 1.25 | 1.2E-02 |
| PCL1606_RS1( | hybrid sensor | 1.25 | 8.3E-03 |
| PCL1606_RS1: | DNA-binding r | 1.25 | 1.3E-02 |
| PCL1606_RS1: | SfnB family su | 1.25 | 2.0E-02 |
| PCL1606_RS2: | EamA family t | 1.25 | 2.3E-04 |
| PCL1606_RS1: | 5,10-methyler | 1.25 | 1.9E-02 |
| PCL1606_RS1( | TonB-depende | 1.25 | 1.5E-02 |
| PCL1606_RS2: | stress-inducec | 1.24 | 2.1E-02 |
| PCL1606_RS1: | feruloyl-CoA s | 1.24 | 2.3E-02 |
| PCL1606_RS1( | RNA polymera | 1.24 | 2.4E-02 |
| PCL1606_RS1( | hypothetical p | 1.24 | 1.9E-02 |
| PCL1606_RS1: | MFS transport | 1.24 | 2.5E-02 |
| PCL1606_RS1: | FAD-depender | 1.24 | 2.0E-02 |
| PCL1606_RS1‹ | LLM class flavi | 1.24 | 1.6E-02 |
| PCL1606_RS1: | MFS transport | 1.23 | 2.3E-02 |
| PCL1606_RS0: | malonate dec: | 1.23 | 2.5E-02 |
| PCL1606_RS0{ | hypothetical p | 1.23 | 4.5E-03 |
| PCL1606_RS1: | transcriptiona | 1.23 | 8.4E-04 |
| PCL1606_RS1‹ | carboxylate-ai | 1.23 | 2.6E-02 |
| PCL1606_RS0: | P-type conjug: | 1.23 | 2.6E-02 |
| PCL1606_RS0( | N-acetylmurai | 1.23 | 2.0E-02 |
| PCL1606_RS1{ | rhodanese | 1.23 | 2.0E-02 |
| PCL1606_RS0: | 3-keto-5-amin | 1.23 | 2.1E-02 |
| PCL1606_RS2‹ | GNAT family a | 1.23 | 9.0E-03 |
| PCL1606_RS1‹ | biopolymer tr: | 1.22 | 2.2E-02 |
| PCL1606_RS1! | energy transd | 1.22 | 2.7E-02 |
| PCL1606_RS1‹ | MFS transport | 1.22 | 8.0E-03 |
| PCL1606_RS1: | maleylacetoac | 1.22 | 2.7E-02 |
| PCL1606_RS3( | hypothetical p | 1.22 | 2.0E-02 |
| PCL1606_RS0: | amine oxidase | 1.22 | 1.1E-08 |
| PCL1606_RS1‹ | siderophore b | 1.22 | 8.2E-03 |
| PCL1606_RS2! | acyl-CoA dehy | 1.21 | 3.6E-09 |
| PCL1606_RS1: | MFS transport | 1.21 | 2.8E-02 |
| PCL1606_RS1‹ | ABC transport | 1.21 | 1.7E-02 |
| PCL1606_RS1( | thiamine bios\ | 1.21 | 1.9E-02 |
| PCL1606_RS1( | MFS transport | 1.21 | 5.0E-03 |

| | | | |
|---|---|---|---|
| PCL1606_RS3( | hypothetical p | 1.21 | 1.4E-03 |
| PCL1606_RS14 | LLM class flavi | 1.20 | 3.0E-02 |
| PCL1606_RS1. | zinc-binding d | 1.20 | 2.3E-02 |
| PCL1606_RS1{ | RND transport | 1.20 | 3.0E-02 |
| PCL1606_RS0( | arginase | 1.20 | 2.9E-02 |
| PCL1606_RS1. | hypothetical p | 1.20 | 7.5E-03 |
| PCL1606_RS1( | 3-hydroxyisob | 1.20 | 1.2E-02 |
| PCL1606_RS1: | two-compone | 1.20 | 1.3E-02 |
| PCL1606_RS1: | class II histone | 1.20 | 2.3E-02 |
| PCL1606_RS1. | salicylaldehyd | 1.20 | 2.1E-02 |
| PCL1606_RS1: | isocitrate lyas | 1.19 | 5.2E-06 |
| PCL1606_RS1{ | glycolate oxid | 1.19 | 2.7E-02 |
| PCL1606_RS14 | hypothetical p | 1.19 | 2.0E-02 |
| PCL1606_RS14 | sodium:protor | 1.19 | 2.8E-02 |
| PCL1606_RS1( | carbonic anhy | 1.18 | 2.4E-02 |
| PCL1606_RS0: | RND transport | 1.18 | 3.2E-02 |
| PCL1606_RS1! | ShlB family he | 1.18 | 1.6E-02 |
| PCL1606_RS24 | amino acid AB | 1.18 | 3.7E-06 |
| PCL1606_RS14 | nitrilotriacetat | 1.18 | 2.3E-02 |
| PCL1606_RS1( | short-chain de | 1.18 | 2.3E-02 |
| PCL1606_RS14 | nitrite reducta | 1.18 | 5.4E-03 |
| PCL1606_RS2! | phosphatidic a | 1.18 | 1.8E-02 |
| PCL1606_RS1. | gentisate 1,2-( | 1.18 | 3.0E-02 |
| PCL1606_RS14 | aspartate ami | 1.18 | 4.6E-04 |
| PCL1606_RS0. | electron trans | 1.17 | 2.7E-02 |
| PCL1606_RS1: | hypothetical p | 1.17 | 2.9E-02 |
| PCL1606_RS14 | NADP(H)-depe | 1.17 | 2.4E-02 |
| PCL1606_RS2( | 2-dehydro-3-c | 1.17 | 2.0E-02 |
| PCL1606_RS2. | 2,5-dioxovaler | 1.17 | 3.4E-02 |
| PCL1606_RS1. | aldehyde deh | 1.17 | 2.3E-02 |
| PCL1606_RS1{ | hypothetical p | 1.17 | 1.9E-02 |
| PCL1606_RS1( | benzoate 1,2-( | 1.17 | 2.3E-02 |
| PCL1606_RS24 | hypothetical p | 1.17 | 3.2E-03 |
| PCL1606_RS1. | cytochrome P4 | 1.17 | 3.0E-02 |
| PCL1606_RS1. | 2-dehydropan | 1.17 | 3.3E-02 |
| PCL1606_RS1( | membrane pr | 1.17 | 1.9E-02 |
| PCL1606_RS1. | 5-carboxymet | 1.17 | 3.5E-02 |
| PCL1606_RS0. | transposase | 1.16 | 2.7E-02 |
| PCL1606_RS14 | achromobacti | 1.16 | 2.7E-02 |
| PCL1606_RS1. | porin | 1.16 | 3.4E-02 |
| PCL1606_RS2. | ketoglutarate | 1.16 | 3.6E-02 |
| PCL1606_RS24 | hypothetical p | 1.16 | 1.5E-02 |
| PCL1606_RS14 | hypothetical p | 1.16 | 3.6E-02 |
| PCL1606_RS2! | hypothetical p | 1.16 | 1.5E-02 |
| PCL1606_RS1: | 5-carboxymet | 1.16 | 6.8E-03 |
| PCL1606_RS1. | acetyl-CoA ace | 1.16 | 3.3E-02 |
| PCL1606_RS0. | conjugal trans | 1.16 | 3.6E-02 |

| | | | |
|---|---|---|---|
| PCL1606_RS1! | phenylacetate | 1.15 | 1.8E-02 |
| PCL1606_RS1( | DUF2063 dom | 1.15 | 2.8E-02 |
| PCL1606_RS0. | conjugal trans | 1.15 | 3.5E-02 |
| PCL1606_RS1. | FMN reductas | 1.15 | 3.7E-02 |
| PCL1606_RS1‹ | sulfurtransfer | 1.15 | 3.2E-02 |
| PCL1606_RS0! | type I secretio | 1.15 | 3.5E-02 |
| PCL1606_RS1. | transcriptiona | 1.15 | 1.4E-02 |
| PCL1606_RS1: | lipase | 1.15 | 3.3E-02 |
| PCL1606_RS1‹ | succinate deh | 1.14 | 3.9E-02 |
| PCL1606_RS2. | endoribonucle | 1.14 | 9.3E-03 |
| PCL1606_RS2. | MFS transport | 1.14 | 2.5E-02 |
| PCL1606_RS0. | peptidase | 1.14 | 3.4E-02 |
| PCL1606_RS0. | electron trans | 1.14 | 3.7E-02 |
| PCL1606_RS0: | hypothetical p | 1.14 | 3.6E-02 |
| PCL1606_RS1: | MFS transport | 1.14 | 2.6E-02 |
| PCL1606_RS1. | FAD-linked ox | 1.14 | 3.9E-02 |
| PCL1606_RS1. | MFS transport | 1.14 | 4.0E-02 |
| PCL1606_RS2! | Sel1 repeat-co | 1.13 | 1.5E-05 |
| PCL1606_RS1: | multidrug DM | 1.13 | 2.0E-02 |
| PCL1606_RS1. | AraC family tr | 1.13 | 3.6E-02 |
| PCL1606_RS2( | anthranilate 1 | 1.12 | 2.9E-02 |
| PCL1606_RS1‹ | ompetence-da | 1.12 | 2.7E-02 |
| PCL1606_RS1( | sugar dehydro | 1.12 | 2.6E-02 |
| PCL1606_RS1: | porin | 1.12 | 3.2E-02 |
| PCL1606_RS1( | hypothetical p | 1.12 | 2.7E-02 |
| PCL1606_RS0. | amino acid de | 1.12 | 2.7E-02 |
| PCL1606_RS1‹ | peptidase | 1.12 | 3.0E-02 |
| PCL1606_RS0: | dehydrogenas | 1.12 | 3.5E-02 |
| PCL1606_RS1! | 2,3-dehydroac | 1.12 | 3.2E-02 |
| PCL1606_RS1. | 3-(3-hydroxy-p | 1.12 | 3.5E-02 |
| PCL1606_RS1‹ | SAM-depende | 1.12 | 3.5E-02 |
| PCL1606_RS2‹ | L-serine amm | 1.11 | 7.4E-03 |
| PCL1606_RS1. | cytochrome c | 1.11 | 4.4E-02 |
| PCL1606_RS1‹ | enterobactin r | 1.11 | 3.5E-02 |
| PCL1606_RS1. | RND transport | 1.11 | 4.3E-02 |
| PCL1606_RS0. | conjugal trans | 1.11 | 1.8E-02 |
| PCL1606_RS1! | phenylacetate | 1.11 | 3.6E-02 |
| PCL1606_RS1. | transposase | 1.10 | 4.0E-02 |
| PCL1606_RS1. | C4-dicarboxyl | 1.10 | 3.9E-02 |
| PCL1606_RS2. | hydroxypyruv | 1.10 | 3.9E-02 |
| PCL1606_RS1( | glucarate deh | 1.10 | 4.3E-02 |
| PCL1606_RS1. | RND transport | 1.10 | 4.2E-02 |
| PCL1606_RS1. | ABC transport | 1.10 | 4.8E-02 |
| PCL1606_RS1. | 2-methylcitrat | 1.09 | 4.4E-02 |
| PCL1606_RS0! | type II secreti | 1.09 | 4.8E-02 |
| PCL1606_RS1‹ | siderophore A | 1.09 | 3.5E-02 |
| PCL1606_RS0! | cytochrome-c | 1.09 | 4.7E-02 |

| | | | |
|---|---|---|---|
| PCL1606_RS2 | isovaleryl-CoA | 1.09 | 4.1E-02 |
| PCL1606_RS1 | hypothetical | 1.09 | 4.8E-02 |
| PCL1606_RS0 | malonate dec | 1.09 | 4.5E-02 |
| PCL1606_RS0 | transcriptiona | 1.09 | 7.1E-03 |
| PCL1606_RS1 | 2-(1,2-epoxy- | 1.09 | 3.9E-02 |
| PCL1606_RS1 | transporter | 1.09 | 3.6E-02 |
| PCL1606_RS2 | amino acid AB | 1.09 | 1.3E-09 |
| PCL1606_RS1 | biopolymer tr | 1.08 | 4.8E-02 |
| PCL1606_RS0 | nitric oxide di | 1.08 | 1.5E-02 |
| PCL1606_RS1 | ABC transport | 1.08 | 4.7E-02 |
| PCL1606_RS2 | MexA family r | 1.08 | 2.6E-02 |
| PCL1606_RS1 | benzoylforma | 1.08 | 3.7E-02 |
| PCL1606_RS1 | type II secreti | 1.08 | 2.3E-02 |
| PCL1606_RS2 | anthranilate d | 1.08 | 3.0E-02 |
| PCL1606_RS1 | alkyl hydroper | 1.08 | 5.0E-02 |
| PCL1606_RS1 | CoA transfera | 1.08 | 3.8E-02 |
| PCL1606_RS1 | aldehyde deh | 1.07 | 3.7E-02 |
| PCL1606_RS1 | monooxygena | 1.07 | 1.1E-02 |
| PCL1606_RS1 | NUDIX domair | 1.07 | 4.7E-02 |
| PCL1606_RS0 | 5-methyltetra | 1.07 | 4.7E-02 |
| PCL1606_RS1 | CusA/CzcA far | 1.07 | 4.3E-02 |
| PCL1606_RS1 | NAD(P)-deper | 1.07 | 4.3E-02 |
| PCL1606_RS1 | hypothetical | 1.06 | 4.3E-02 |
| PCL1606_RS1 | alcohol dehyd | 1.06 | 4.0E-02 |
| PCL1606_RS1 | Vanillate O-de | 1.06 | 4.1E-03 |
| PCL1606_RS2 | urea transpor | 1.06 | 4.7E-02 |
| PCL1606_RS1 | TonB-depende | 1.06 | 2.1E-05 |
| PCL1606_RS0 | type II secreti | 1.06 | 4.6E-02 |
| PCL1606_RS1 | alcohol dehyd | 1.05 | 2.3E-02 |
| PCL1606_RS1 | type II secreti | 1.05 | 3.9E-02 |
| PCL1606_RS0 | (Fe-S)-binding | 1.05 | 4.3E-02 |
| PCL1606_RS1 | amino acid tra | 1.05 | 2.0E-02 |
| PCL1606_RS1 | multidrug effl | 1.05 | 4.4E-02 |
| PCL1606_RS1 | MexE family n | 1.05 | 1.1E-04 |
| PCL1606_RS1 | glycolate oxid | 1.05 | 2.2E-02 |
| PCL1606_RS1 | DNA-binding r | 1.05 | 4.6E-02 |
| PCL1606_RS0 | conjugal trans | 1.04 | 2.8E-02 |
| PCL1606_RS0 | peptidase C69 | 1.04 | 3.1E-02 |
| PCL1606_RS1 | antibiotic bios | 1.04 | 4.6E-02 |
| PCL1606_RS1 | hypothetical | 1.04 | 4.1E-02 |
| PCL1606_RS2 | fumarylacetoa | 1.04 | 4.7E-02 |
| PCL1606_RS1 | 5-carboxymet | 1.04 | 7.3E-03 |
| PCL1606_RS1 | LuxR family tr | 1.04 | 3.6E-02 |
| PCL1606_RS1 | quercetin 2,3- | 1.04 | 1.1E-02 |
| PCL1606_RS0 | sulfurase | 1.04 | 3.4E-04 |
| PCL1606_RS1 | peptidase | 1.04 | 2.0E-02 |
| PCL1606_RS1 | DUF421 doma | 1.03 | 2.0E-02 |

| | | | |
|---|---|---|---|
| PCL1606_RS1( | MFS transport | 1.03 | 4.8E-02 |
| PCL1606_RS1( | RNA polymera | 1.03 | 6.7E-03 |
| PCL1606_RS14 | ABC transport | 1.03 | 4.1E-02 |
| PCL1606_RS0! | potassium-tra | 1.03 | 3.9E-02 |
| PCL1606_RS1( | muconolactor | 1.03 | 3.2E-02 |
| PCL1606_RS1. | copper-translo | 1.03 | 1.1E-03 |
| PCL1606_RS0. | RepA replicati | 1.03 | 3.1E-02 |
| PCL1606_RS1( | hypothetical p | 1.02 | 2.6E-02 |
| PCL1606_RS14 | ABC transport | 1.02 | 4.1E-02 |
| PCL1606_RS14 | acid phosphat | 1.02 | 4.0E-02 |
| PCL1606_RS0! | paraquat-indu | 1.02 | 6.5E-03 |
| PCL1606_RS2. | MFS transport | 1.02 | 5.9E-03 |
| PCL1606_RS0. | malonate tran | 1.01 | 1.8E-02 |
| PCL1606_RS0: | flavin reducta: | 1.01 | 3.0E-02 |
| PCL1606_RS1( | phosphohydro | 1.01 | 2.0E-02 |
| PCL1606_RS1( | hypothetical p | 1.01 | 4.9E-02 |
| PCL1606_RS1( | tRNA (adenine | 1.01 | 2.7E-02 |
| PCL1606_RS2. | hypothetical p | 1.01 | 2.0E-02 |
| PCL1606_RS1: | MarR family tr | 1.01 | 8.9E-04 |
| PCL1606_RS1{ | DUF4865 dom | 1.00 | 4.1E-02 |
| PCL1606_RS0. | N-methylproli | 1.00 | 2.7E-02 |
| PCL1606_RS0! | paraquat-indu | 1.00 | 2.1E-02 |
| PCL1606_RS2: | ABC transport | -1.00 | 8.9E-09 |
| PCL1606_RS2: | tail protein | -1.01 | 2.8E-04 |
| PCL1606_RS0( | type IV secreti | -1.01 | 7.4E-05 |
| PCL1606_RS2! | hypothetical p | -1.02 | 2.3E-25 |
| PCL1606_RS3( | hypothetical p | -1.03 | 9.1E-05 |
| PCL1606_RS2. | MFS transport | -1.04 | 5.7E-04 |
| PCL1606_RS2: | 2-hydroxyacid | -1.04 | 7.9E-05 |
| PCL1606_RS0( | type VI secreti | -1.04 | 2.9E-04 |
| PCL1606_RS1! | lipoprotein | -1.04 | 7.8E-05 |
| PCL1606_RS1: | HlyD family ty | -1.04 | 3.4E-02 |
| PCL1606_RS2: | tail sheath pro | -1.05 | 1.5E-13 |
| PCL1606_RS0( | hypothetical p | -1.06 | 1.3E-06 |
| PCL1606_RS2: | hypothetical p | -1.06 | 7.6E-03 |
| PCL1606_RS1: | aminotransfer | -1.07 | 6.6E-06 |
| PCL1606_RS1{ | tryptophan sy | -1.07 | 3.1E-05 |
| PCL1606_RS0. | pilus assembly | -1.08 | 2.7E-02 |
| PCL1606_RS1. | hypothetical p | -1.08 | 1.3E-03 |
| PCL1606_RS0( | hypothetical p | -1.10 | 5.2E-09 |
| PCL1606_RS2! | type I secretio | -1.12 | 1.2E-17 |
| PCL1606_RS0( | type VI secreti | -1.12 | 1.4E-08 |
| PCL1606_RS2: | hypothetical p | -1.13 | 1.5E-08 |
| PCL1606_RS2! | hypothetical p | -1.13 | 1.7E-07 |
| PCL1606_RS1: | flavin reducta: | -1.14 | 1.5E-07 |
| PCL1606_RS0. | sugar ABC trar | -1.15 | 1.7E-05 |
| PCL1606_RS0( | membrane pr | -1.16 | 1.4E-11 |

| | | | |
|---|---|---|---|
| PCL1606_RS2: | phage tail pro | -1.16 | 1.2E-11 |
| PCL1606_RS0: | type I glyceral | -1.17 | 1.5E-02 |
| PCL1606_RS1: | monodechloro | -1.18 | 1.7E-05 |
| PCL1606_RS0( | protein phosp | -1.23 | 2.0E-09 |
| PCL1606_RS1{ | serine 3-dehy | -1.23 | 4.4E-04 |
| PCL1606_RS1: | chitin-binding | -1.26 | 3.8E-17 |
| PCL1606_RS1: | peptidase | -1.26 | 8.2E-03 |
| PCL1606_RS2: | ArsR family tra | -1.27 | 1.8E-06 |
| PCL1606_RS1: | polyurethanas | -1.28 | 9.7E-03 |
| PCL1606_RS1( | non-ribosomal | -1.29 | 2.0E-03 |
| PCL1606_RS1: | outer membra | -1.29 | 2.0E-03 |
| PCL1606_RS1: | 3-ketoacyl-AC | -1.29 | 5.1E-06 |
| PCL1606_RS1: | potassium tra | -1.32 | 1.2E-05 |
| PCL1606_RS1: | protease | -1.34 | 5.4E-03 |
| PCL1606_RS2! | DUF2474 dom | -1.36 | 5.4E-04 |
| PCL1606_RS1: | serine proteas | -1.37 | 1.0E-03 |
| PCL1606_RS1: | flavin reducta: | -1.38 | 5.5E-10 |
| PCL1606_RS1: | polyurethanas | -1.39 | 2.3E-07 |
| PCL1606_RS1! | sugar ABC trar | -1.39 | 3.4E-06 |
| PCL1606_RS2{ | glutamine AB( | -1.43 | 6.5E-08 |
| PCL1606_RS2: | membrane pr | -1.45 | 1.4E-03 |
| PCL1606_RS2: | leucyl aminop | -1.46 | 6.4E-05 |
| PCL1606_RS1: | FAD-depende | -1.46 | 1.1E-09 |
| PCL1606_RS1: | chitinase | -1.47 | 5.5E-21 |
| PCL1606_RS1: | (2Fe-2S) ferre | -1.49 | 2.5E-07 |
| PCL1606_RS0 | ubiquinol oxid | -1.62 | 1.0E-10 |
| PCL1606_RS2! | tryptophan sy | -1.63 | 1.4E-21 |
| PCL1606_RS2( | copper chaper | -1.72 | 1.2E-07 |
| PCL1606_RS0 | cytochrome u | -1.74 | 2.6E-10 |
| PCL1606_RS2! | tryptophan sy | -1.78 | 5.7E-21 |
| PCL1606_RS2( | peptidase | -1.79 | 1.6E-08 |
| PCL1606_RS1: | serine 3-dehy | -1.87 | 5.6E-04 |
| PCL1606_RS2( | hypothetical p | -1.90 | 3.5E-25 |
| PCL1606_RS2( | TonB-depende | -1.99 | 2.9E-09 |
